# Getting to know each other: PPIMem, a novel approach for predicting transmembrane protein-protein complexes

**DOI:** 10.1101/871590

**Authors:** Georges Khazen, Aram Gyulkhandanian, Tina Issa, Rachid C. Maroun

## Abstract

Because of their considerable number and diversity, membrane proteins and their macromolecular complexes represent the functional units of cells. Their quaternary structure may be stabilized by interactions between the α-helices of different proteins in the hydrophobic region of the cell membrane. Membrane proteins also represent potential pharmacological targets par excellence for various diseases. Unfortunately, their experimental 3D structure and that of their complexes with intramembrane interacting partners are scarce due to technical difficulties. To overcome this key problem, we devised PPIMem, a computational approach for the specific prediction of higher-order structures of α-helical transmembrane proteins. The novel approach involves identification of the amino acid residues at the interface of complexes with a 3D structure. The identified residues compose then interaction motifs that are conveniently expressed as mathematical regular expressions. These are used for motif search in databases, and for the prediction of intramembrane protein-protein complexes. Our template interface-based approach predicted 21, 544 binary complexes between 1, 504 eukaryotic plasma membrane proteins across 39 species. We compared our predictions to experimental datasets of protein-protein interactions as a first validation method. The PPIMem online database with the annotated predicted interactions is implemented as a web server and can be accessed directly at https://transint.shinyapps.io/transint/.

## INTRODUCTION

Proteins are the core of cell machinery in organisms, and membrane proteins (MPs) encompass a broad variety of functions, including but not limited to activation, transport, degradation, stabilization, apoptosis, cell signaling and participation in the production of other proteins. Because protein complexes represent functional units of cells, it is fundamental to understand the interactions between their components. More specifically, knowledge of the 3D structure of the MPs and interfaces involved in macromolecular complex formation remains a fundamental phenomenon governing all processes of life (1). Therefore, MPs represent ultimate potential pharmacological targets in human health and disease because they include many families involved in protein-protein interaction (PPI) networks, leading to different physiological processes. Current estimates suggest that about half of all drugs target MPs. Moreover, transmembrane (TM) helix-helix interactions, through the lipid-embedded domains, lead to oligomer formation and guide the assembly and function of many cell proteins and receptors (2). In addition, the assembly of MPs may lead to emergent properties, as a relationship exists between oligomerization and function (3, 4). On the other hand, it is important to understand the functional effects of mutations at the interface because these may modify the assembly of MPs to form multiprotein complexes, leading to disruptions in networks of interactions and phenotypic changes. These mutations may possibly lead to the development of various genetic diseases (5). Thus, the role of TM domains in protein function is crucial. These TM proteins span the cell membrane in its entirety and represent approximately one-third of the proteomes of organisms (6). In eukaryotes, they come mostly in the form of α-helical bundles that cross different types of cell membranes and are approximately 3 × 10^5^ in number, excluding polymorphisms or rare mutations. The class of α-helical TM proteins has a higher diversity than their β-barrel counterparts. Thus, the number of possible binary and multiple interactions between them is vastly larger (7). Estimates of the total number of human PPIs range from 130, 000 to 600, 000 (8–10), to 3, 000, 000 (11), which is several orders of magnitude larger than that of the *D. melanogaster* interactome. In this study, the plasma membrane protein interactome is the complete set of direct interactions that take place in the membrane proteome of α-helical integral-to-membrane transmembrane proteins.

High-throughput experimental and theoretical approaches are being used to build protein-protein interaction (PPI) networks. The data covering PPIs are increasing exponentially. Indeed, in the year 2012 more than 235, 000 binary interactions were reported (12). Most protein interaction databases (DBs) (11–31) offer general information about experimentally validated PPIs of all types. The IMEx Consortium and the PSICQUIC service (http://www.ebi.ac.uk) group all data dealing with non-redundant protein interactions at one interface (32). Nevertheless, the DBs are mostly concerned with water-soluble globular proteins and the juxtamembrane interactions of MPs. However, unlike globular proteins, MPs are water-insoluble, their exterior is much more hydrophobic than the interior because of allosteric interactions with the lipid environment, and they lose their native structure when removed from their natural membrane environment. Their thorough investigation has thus lagged due to this technical difficulty (33).

Proteome-wide maps of the human interactome network have been generated in the past (24, 28, 34). Traditional <u>experimental</u> techniques such as yeast two-hybrid (Y2H) assays (35) are not well suited for identifying MP interactions. Other assays, such as Y2H, are depleted of MPs or unreliable (36). A new biochemical technique has been developed (MYTH); however, only a limited number of membrane complexes have been hitherto determined employing it (28, 34). This procedure has been significantly extended (MaMTH) (37). However, to the best of our knowledge, MaMTH has not been used as a systematic screening assay to map the human MP interactome. Another approach for the identification of integral membrane PPIs in yeast used integral membrane proteins as baits (38). An alternative method advanced a novel MYTH yeast proteomics technology, allowing the characterization of interaction partners of full-length GPCRs in a drug-dependent manner (39).

On the other hand, a variety of recent methods for the <u>prediction</u> of PPIs have flourished based on: i) machine learning and classifiers based on sequence alone (40, 41), and deep learning (42); ii) template structures (43, 44). Again, most of the approaches are parameterized on soluble globular proteins only, as in (45). *Ab-initio* prediction of MP interfaces is rendered difficult, as these include amino acid compositions that are not radically different from the rest of the protein surface, even though they are better conserved (46). Indeed, MP interfaces are decorated with hydrophobic residues on the membrane-exposed surface.

To circumvent this problem, we developed a 3D structure knowledge-based approach to predict complexes between TM proteins. Indeed, we incorporate the 3D structure as it provides additional information (surface accessibility, residue neighbors, etc.) not readily present in the amino acid sequence alone. The approach is based on the detection of α-helical TM non-bonded contact residues between different chains in experimental 3D structures of MP multimers reported in the PDB DB (47), and validated by the OPM DB (48). Querying the PDBsum DB (49) thereafter, we obtained the atomic details of the membrane protein-membrane protein interfaces, that is, the contacts at the intermolecular interface of the complexes, through the PDBsum DB. We then gathered those amino acids at the recognition interface to generate regular expressions or patterns that represented in a linear form the interaction motifs in space. The regular expressions include wildcard regions between the interface contact residues representing the residues not in contact, even though they may be exposed to the membrane lipids. With this information in hand, we proceeded to search in the UniProtKB DB (50) of protein sequences the obtained motifs in other MPs. To extend our search, we allowed certain degrees of mutation in the membrane-embedded interface contact residues. Our assumptions are that (i) the interface residues in the experimentally determined structures of complexes between α-helical transmembrane proteins are responsible for the interaction; and (ii) homologs of precisely the benchmark template interface motifs are expected to interact analogously (51). This latter assumption is opposed to a global approach that limits the search to functionally related partners or to overall sequence homolog partners without paying attention to the specific sequence at the interface. In all cases, it is reasonable to assume that the number of interface motifs is limited in nature (52). Thus, we do not focus on the overall sequence homology, such as in other template-based predictions (43, 53). The linear 1D motifs we obtain are useful for quick local sequence searches, represent 3D epitopes implicitly and have the advantage of encompassing implicitly experimental tertiary and quaternary structures, as opposed to other approaches using only the primary structure.

In this work, we focus on the plasma membrane of eukaryotes, ensuring that we probe proteins within the same subcellular localization. In addition, the MPs that compose a complex belong always to the same organism.

## MATERIALS AND METHODS

### Algorithm

#### Data mining and filtering

Figure 1 summarizes the steps followed for collecting, filtering, and processing the input data from several sources and generating the output data based on regular expressions. We started by obtaining a list of all eukaryote “reviewed” proteins from UniProtKB DB, a manually annotated and reviewed DB of proteins with experimental evidence of their existence (50). We mined proteins with cellular component annotations that matched the following GO annotations (54): “integral component of membrane” (GO:0016021) <u>and</u> “plasma membrane” (GO:0005886); <u>or</u> “integral component of plasma membrane” (GO:0005887). Thus, we considered only proteins characterized as membrane proteins. From the resulting list, we identified the subset of proteins with an experimental 3D structure in the PDB DB, a resource containing the 3D shapes of biological macromolecules in the form of a complex spanning the TM region. We retained those with at least six buried interacting residues in each monomer (eight buried residues is typical of biological interfaces, (55)). This amount is the minimum for presenting an accessible surface area leading to quaternary structure. To obtain a high quality homogeneous dataset, we adopted stringent selection conditions. Thus, for the PDB structure to be considered valid, it had to have a high-resolution limit of 3.5 Å or better if it was obtained by X-ray crystallography or cryoEM. We took in all different conformational states, regardless of the presence or absence of ligand(s), pH, symmetry group, apo or holo form, and allosteric effects, unless the differences modified the set of interface residues. We eliminated PDB MPs presenting engineered mutations, insertions, and deletions in the TM segment with respect to the wild-type or natural variant sequence in UniProtKB, just like chimeras in which the xenophobic part is TM. We also ignored redundant MPs, those whose 3D structures showed no significant TM segments, and pair interactions that are redundant due to symmetry. Manual curation excluded non-parallel configurations of the protomers, head-to-head or head-to-tail orientations (resulting most probably from crystal interfaces), out-of-membrane interactions only, and TM segments that do not interact, as dictated by the cell membrane. Intramolecular interactions within each protomer of an oligomer, as well as with the lipids and detergents environment are included implicitly. The PDBsum DB contains the PDB molecules that make up a complex and the interactions between them. This information delivers what interface residues of protomer A interact with what interface residues of protomer or chain B, and what protomer A interacts precisely with what protomer B and not another.

**Figure 1.**
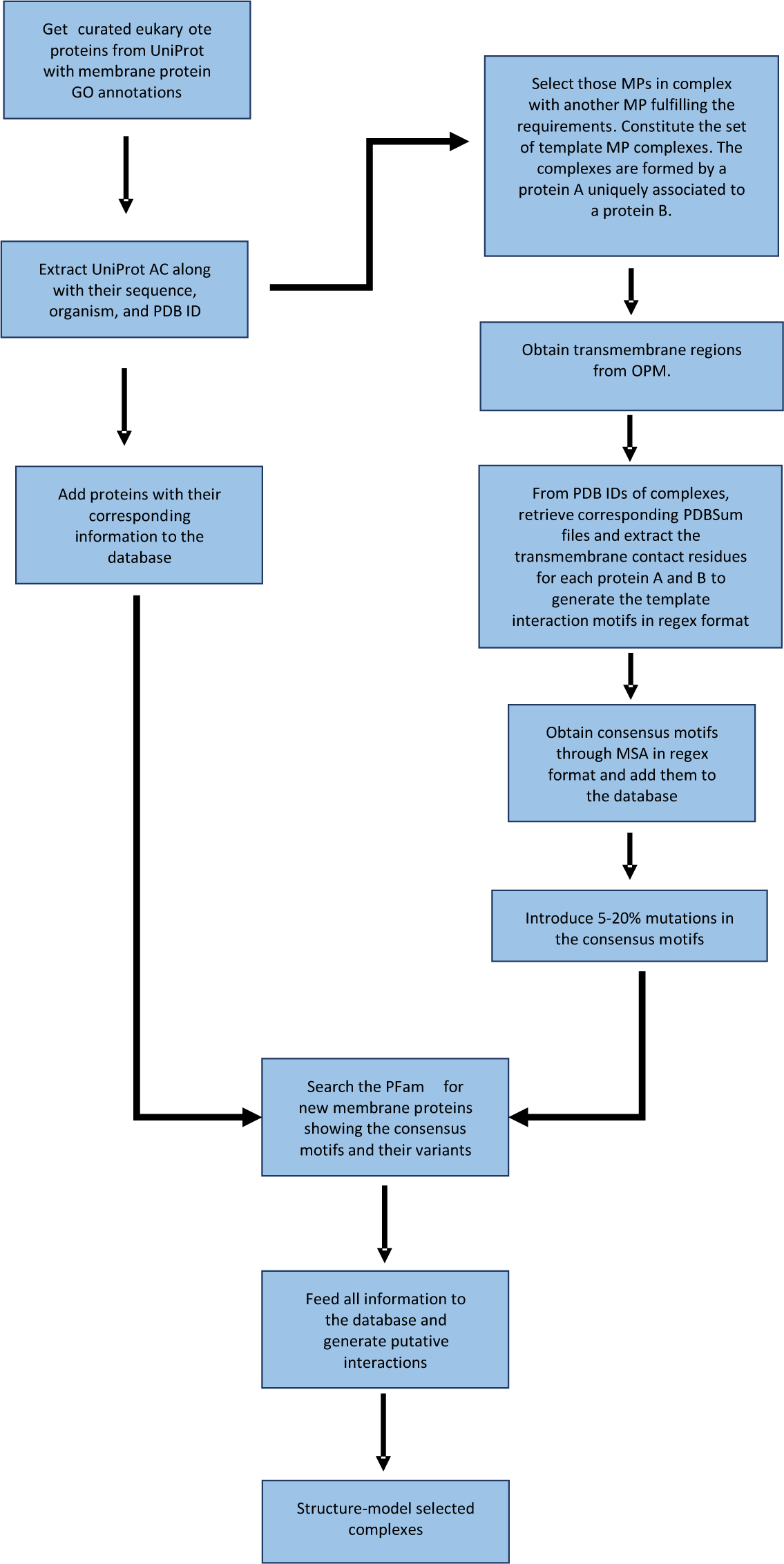
Flowchart illustrating the PPIMem algorithm: from information retrieval to detection of recognition motifs to generation of putative interactions, to 3D modeling of complexes.

Finally, to ensure that the oligomer structures we considered are quaternary structures with biological sense, we used EPPIC, a protein-protein interface classifier (55–57), and PRODIGY, a classifier of biological interfaces in protein complexes (58, 59) to distinguish between crystallographic and biological assemblies.

#### Motif extraction

To choose the PDB structures to work on, we referred to the OPM DB (60), which provides the orientation of known spatial arrangements of unique structures of representative MPs coming from the PDB with respect to the hydrocarbon core of the lipid bilayer. We chose all the PDB structures that mapped to the UniProtKB extracted MPs and pulled out all available PDBsum files of these structures. We double-checked the chosen PDB structures with the MPStruc DB of MPs of known 3D structure (https://blanco.biomol.uci.edu/mpstruc/). PDBsum contains atomic non-bonded contacts between amino acid residues at the interface of molecules in a multimer complex. We used the information in PDBsum to extract the intermolecular contact residues. We filled in with the non-contact residues in the sequence, that is, those between the contact residues, as wildcards. Thereafter, we defined the binding motifs by one to its partner protein B. Because we are only interested in the recognition site at the TM interface region, we ensured that each interacting residue belonged to the TM part of the sequence. We represented our motifs using the Regex format (https://www.regular-expressions.info/) and denoted the TM contact residues by their one-letter symbol, the residues in between (representing the wildcard regions) by a dot, and the curly braces for the consecutive occurrences of the wildcard. As an example, we take UniProtKB membrane protein P04626 (receptor tyrosine-protein kinase ErbB2). The code of the spatial structure of the ErbB2 dimeric transmembrane domain in the PDB is 2N2A. Its PDBsum entry contains non-bonded contacts across the surface for each of the chains A and B. The residues in between constitute the non-contact residues wildcard and are represented by curly braces. PDBsum lists chain A residues I12, V16, L20, V23, L24, V27, and L31 as establishing contacts with chain B. The result of the contact motif for chain A is ^12^IX_3_VX_3_LX_2_VLX_2_VX_3_L^31^ (Figure 2), equivalent in the regular expression notation to I.{3}V.{3}L.{2}VL.{2}V.{3}L. As this is a homodimer and the motif is the only one, the contact motif is the same for chain B. The wild cards {} represent extramembrane loops or inner protein subdomains of different lengths that do not establish membrane interface contacts. We do not focus on these variable segments although of course, those loops may be involved in interactions in the cytosol or in the extracellular milieu; consequently, we do not look in our data mining at whether or not those segments are conserved in the amino acid sequence. We focus only on the intramembrane interface contact residues. If the resulting motif is an intramembrane binding motif, the types of variable residues or the length of the interdomains should not be essential.

**Figure 2.**
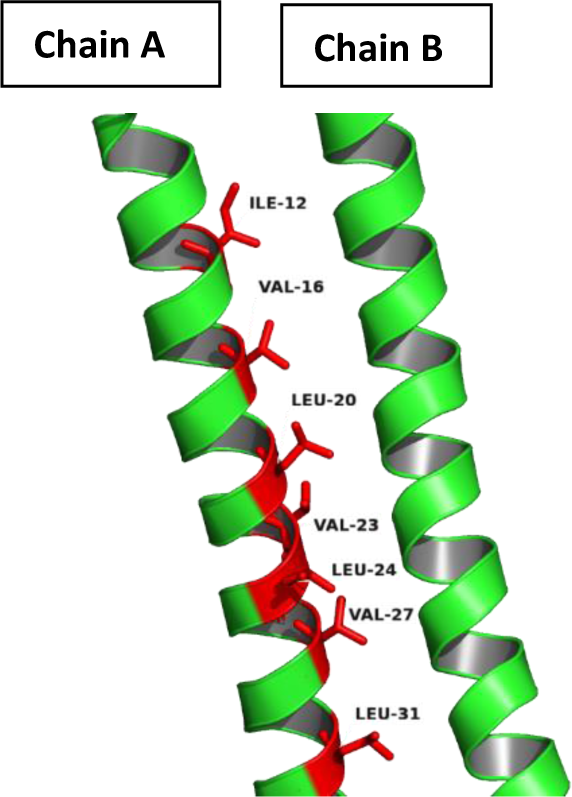
Contact interface residues of chain A of the ErbB2 dimeric transmembrane domain (UniProtKB P04626) of sequence ^12^**Ile**-Ser-Ala-Val-**Val**-Gly-Ile-Leu-**Leu**-Val-Val-**Val**-**Leu**-Gly-Val-**Val**-Phe-Gly-Ile-**Leu**^31^. Contact residues are in bold and compose the motif I.{3}V.{3}L.{2}VL.{2}V.{3}L.

Finally, to verify the consistency of our algorithm and the validity of the obtained binding motifs (that they will not match with many sequences just by random), we sent several BLAST (https://blast.ncbi.nlm.nih.gov) searches against the PDB of several motifs or their parts thereof (in case they were separated by long stretches of non-contact amino acid residues between the contact residues). We always recovered the corresponding template PDB files, as well as related PDB codes, among the maxima BLAST scores and with query covers of 100 (not shown).

#### Searching for identified motifs in other protein sequences

To eliminate redundancy in our data, we grouped motifs derived from the application of mutation rates to the native motifs to build a consensus motif for each cluster derived from multiple sequence alignments (MSA). This allowed us to retain conserved interface residues, avoiding potentially deleterious effects on protein assembly. The mutation rates of the contact residues ranged from 0% (exact match) to 20%, with increments of 5%. In this way, we wished to amplify our search, allowing us to find other protein sequences with homologous motifs. The mutation procedure was applied independently to each protomer, on each motif. Most of the time, the two motifs from a homodimer are identical. However, in some cases, one of the monomers of a homodimer may reveal another interacting surface. In this case, the corresponding motif is different.

Thus, we queried our consensus motifs against the Pfam dataset (https://pfam.xfam.org/). We defined a COST parameter as the number of amino acid mutations allowed anywhere in the motif, depending on the number of contact residues it contains and the mutation rate for the run; we assigned a score of 0 to both insertions and deletions to ensure that no contact residue is lost. For instance, when generating new sites from a valid motif with eight contact residues, COST = 2 for a mutation rate of 20% (8 × 0.20 = 1.6 rounded up to 2). The values of COST_A and COST_B vary from 0 to 4, implying that there are motifs with up to 20 contact residues. In which case, the contact residues may be shared by several submotifs in TM α-helices.

To keep track of which A motif interacts with which motif B, we kept the motifs in separate pools. In other words, the rationale for keeping the two sets A and B of motifs distinct is that a given motif A matches a given motif B and not just any other one. Subsequently, we paired the predicted motifs from novel interactions based on the PDBsum validated interactions. In this way, we were sure that pattern A from the new protein A “binds” to its complementary pattern B of new protein B. Because motifs can be found fortuitously anywhere in the sequence, we considered only those motifs belonging to the TM region. We also checked which MPs showed the motifs in their PDB structures, if available.

### Implementation

#### The PPIMem database

Thereafter, we built a heterogeneous DB named PPIMem, which contains all the found interactions. The DB is the result of the implementation of the fully automated template-based recognition site search pipeline. We used a MySQL DB to maintain all the information collected in a structured manner. To access the DB and search for our data, we built a web interface using PHP and HTML5, which allows the user to query the needed information. Users can query the DB for obtaining motifs by entering a UniProtKB AC, a gene name, a type of organism, a mutation percentage, or a motif of interest or part thereof using the Regex format. A link to the UniProtKB DB site for each UniProtKB AC is available. The user can choose more than one filter option when querying and will exclusively obtain interactions thought to occur in TM regions between plasma membrane proteins of the same species. The user can also adjust the values of the following parameters: Number of contact residues, Mutation rate, COST and Valid, as an interval or as a fixed value. Homodimers of the TMs dealt with do not appear in the database, as all PPIMem TM proteins are expected to form them. We will update our DB following each release of the UniProtKB and OPM datasets and then regenerate all statistics.

#### Molecular docking

The predicted interface residues can be used as constraints to reconstruct the structure of dimers through docking. We consider successful docking as a partial validation procedure. To illustrate our approach, we generated 3D binary complexes derived from selected predicted pairs of MPs with the goal of looking at their internal architecture. To do so, we searched for a protein-protein docking program that would allow us to perform a steered docking simulation using the epitopes extracted from the molecular interface of the complex. We processed and analyzed a considerable number of experimental 3D MP structure files using several docking programs. As we know the docking interfaces, we did not perform in general *ab initio* molecular calculations. Neither were we concerned whether the docking program was trained on sets composed primarily of soluble proteins.

Even though several tested docking programs were sufficiently precise, we decided to use GRAMM-X (61, 62) for creating the novel protein-protein 3D complexes. GRAMM-X has an option in which the user can submit those residues that might form the interface between the “receptor” and the “ligand.” The program also has options for listing the residues required to be in contact. To verify the performance of GRAMM-X for MPs, we benchmarked it against several MP complexes in the PDB. GRAMM-X was indeed able to reproduce many of the experimental MP complexes. For molecular docking simulation and identification of the PPIMem-predicted complexes, we chose examples in which the 3D PDB structures of proteins were already available or represented highly homologous templates. After docking, we manually curated the output by filtering out non-parallel, perpendicular, or oblique protomer pairs, regardless of the calculated energy. We also considered the topology of the MP in the membrane. We visualized the obtained 3D structures of the complexes using PyMol 2.4 (www.pymol.org).

## RESULTS

### Automated pipeline to predict and explore thousands of novel transmembrane protein-protein direct interactions (MPPI)

UniProtKB provided us with 13, 352 MPs that include the GO annotations mentioned in the S&M section. Overall, these proteins mapped to 954 distinct oligomer MP PDB structures. As we focused on structures that satisfy the requirements signified in the S&M section, we were able to validate only ∼50 PDB files of MP-MP complexes. After checking which corresponding PDBsum files to consider, we ended up with 53 non-redundant template interactions, associated with 48 unique reviewed UniProtKB entries across species and forming our benchmark. Fifty of these interactions correspond to structural homomers and three to structural heteromers (Table S1, yellow highlighting). The set includes X-ray, solution NMR, and electron microscopy structures. The MPs include families like receptor tyrosine kinases, TLRs, ion channels, Cys-Loop and immune receptors, gap junctions, transporters, and GPCRs. In these, besides *H. sapiens*, a taxonomically diverse set of organisms across the eukaryotic kingdom of the tree of life is represented: *A. thaliana, B. taurus, G. gallus, M. musculus, O. sativa, R. norvegicus, S. cerevisiae*, and several others. On another hand, there are 21 structures of complexes between bitopic proteins, 32 between polytopic proteins, and one mixed bitopic-polytopic protein complex in our template set. The oligomerization order is adequately represented, with 23 homo 2-mers, five homo 3-mers, 12 homo 4-mers, one hetero 4-mer, four homo 5-mers, three hetero 6-mers, one hetero 10-mer, and one homo 12-mer. The occurrences of the number of TM helices for each of the protomers of the reference MP complexes shows that bitopic single-span helix complexes are the most preponderant; these are followed by polytopic MPs composed of 6, 4, 7, 9, 2, 8, and 10 and 12 helices. The following EC numbers are represented in this set: EC 2 (10 transferases), EC 3 (one hydrolase), and EC 7 (one translocase). When verifying the protein-protein interfaces in the complexes for the type of assembly they form (crystallographic or biological), we found that all the X-ray or electron microscopy complexes were classified as biological (Table S3). We did not submit the NMR-determined complexes to the test. Therefore, this is the number of experimentally solved structures of MP protein complexes that we used as the benchmark set.

The PPIMem pipeline results indicate that there are 1504 unique MPs involved in the predicted complexes, going from UniProtKB accession codes A0PJK1 to Q9Y6W8. Of these, 417 are human (Table S5). The total number of motifs found after removing redundancies due to different chains of the same structures interacting more than once was 98 (Table 1).

**Table 1.**
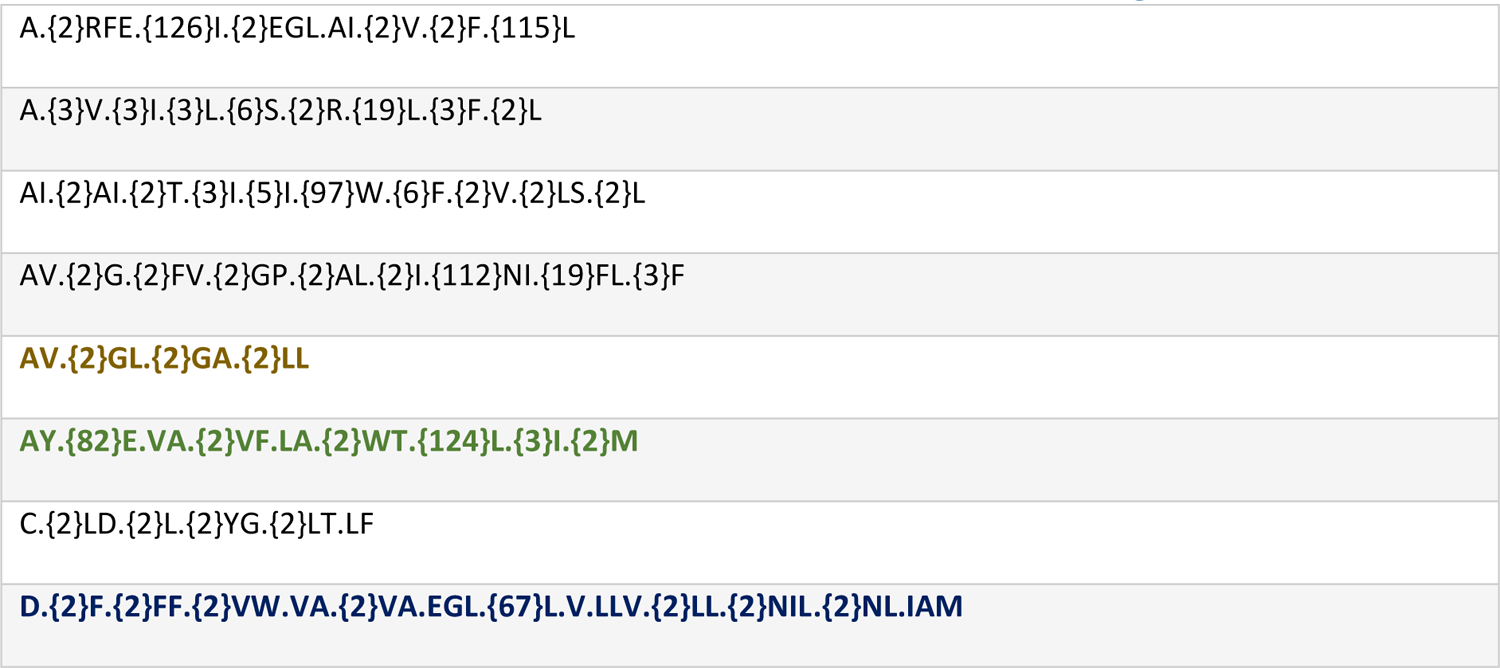
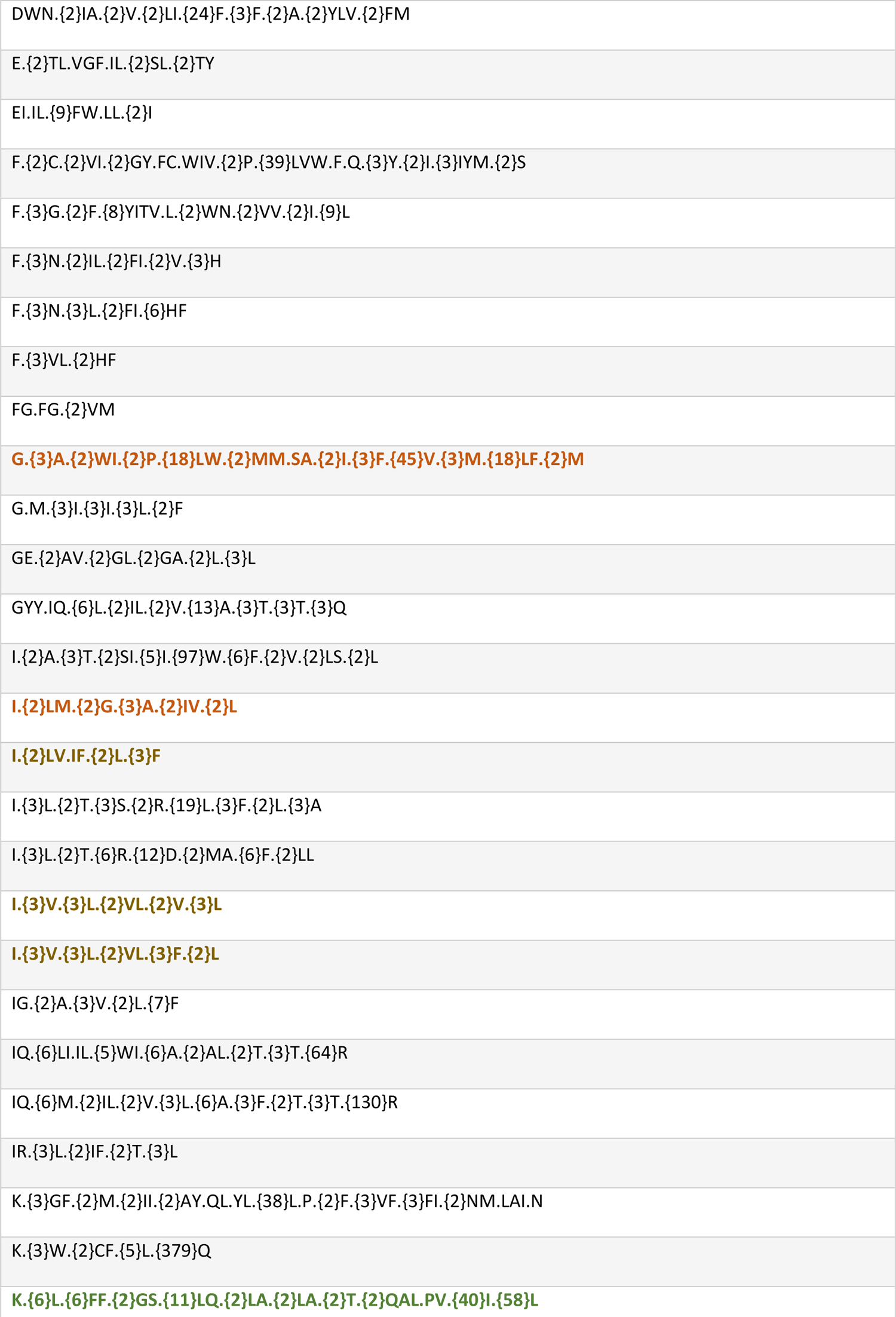
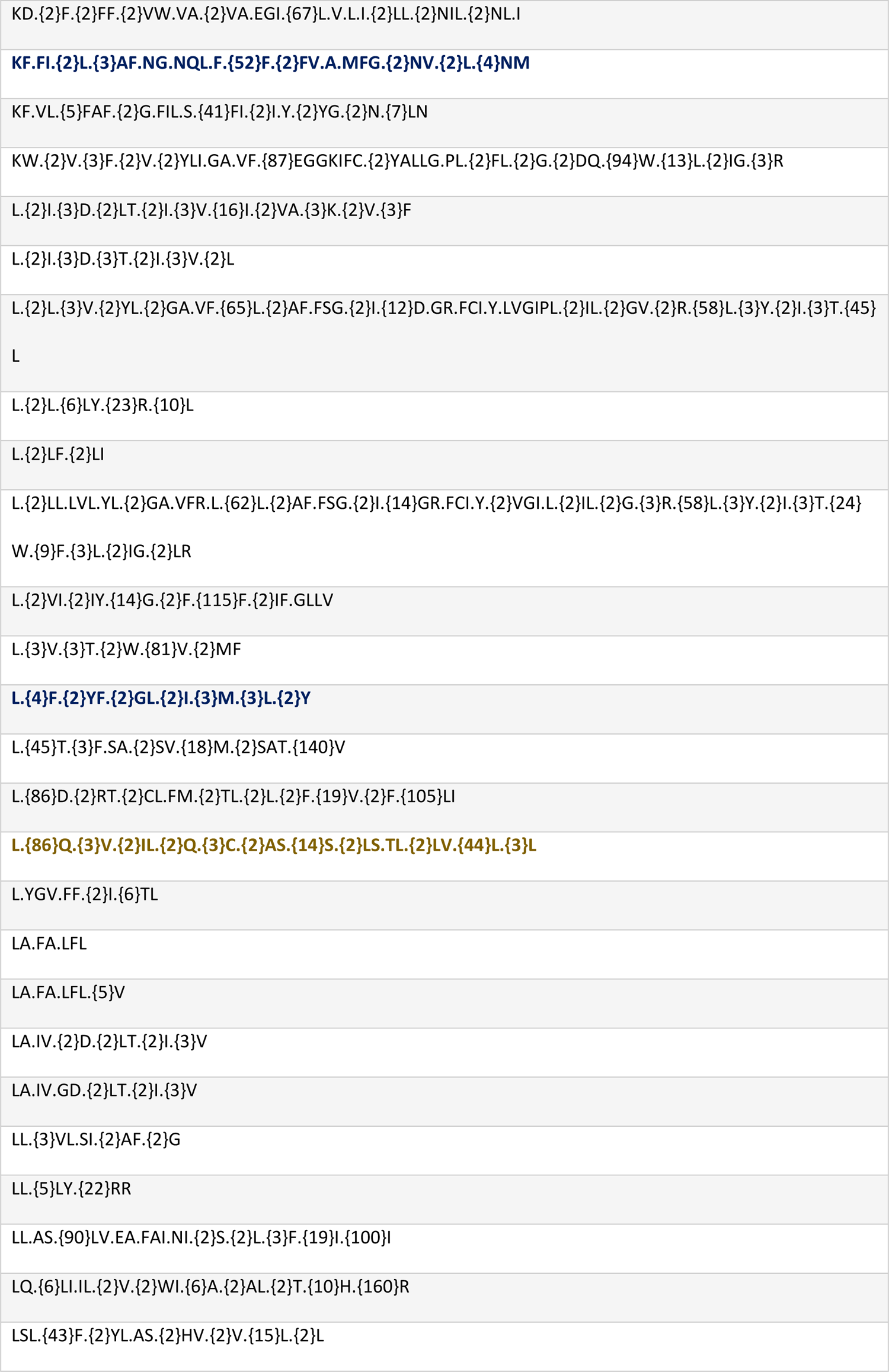
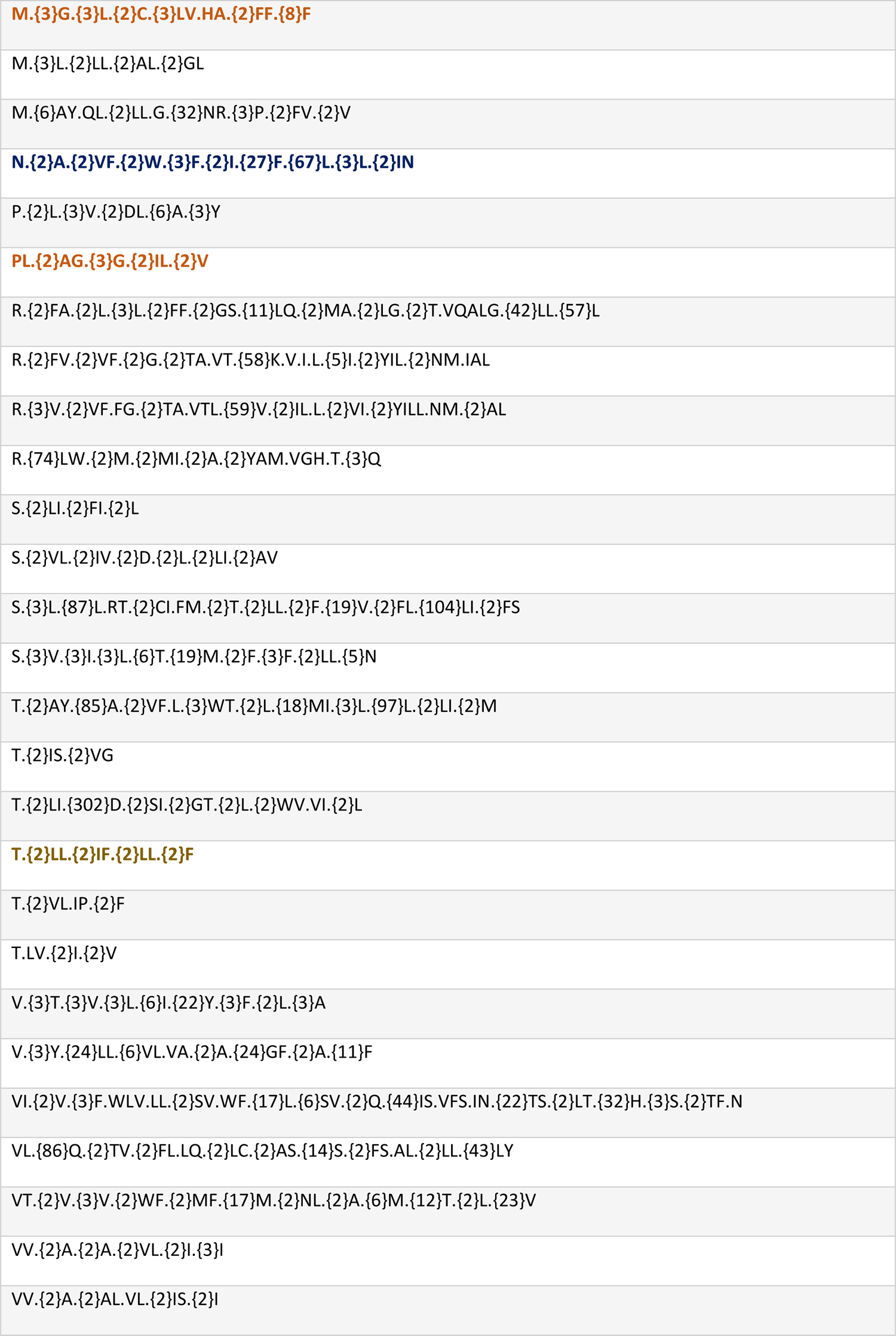
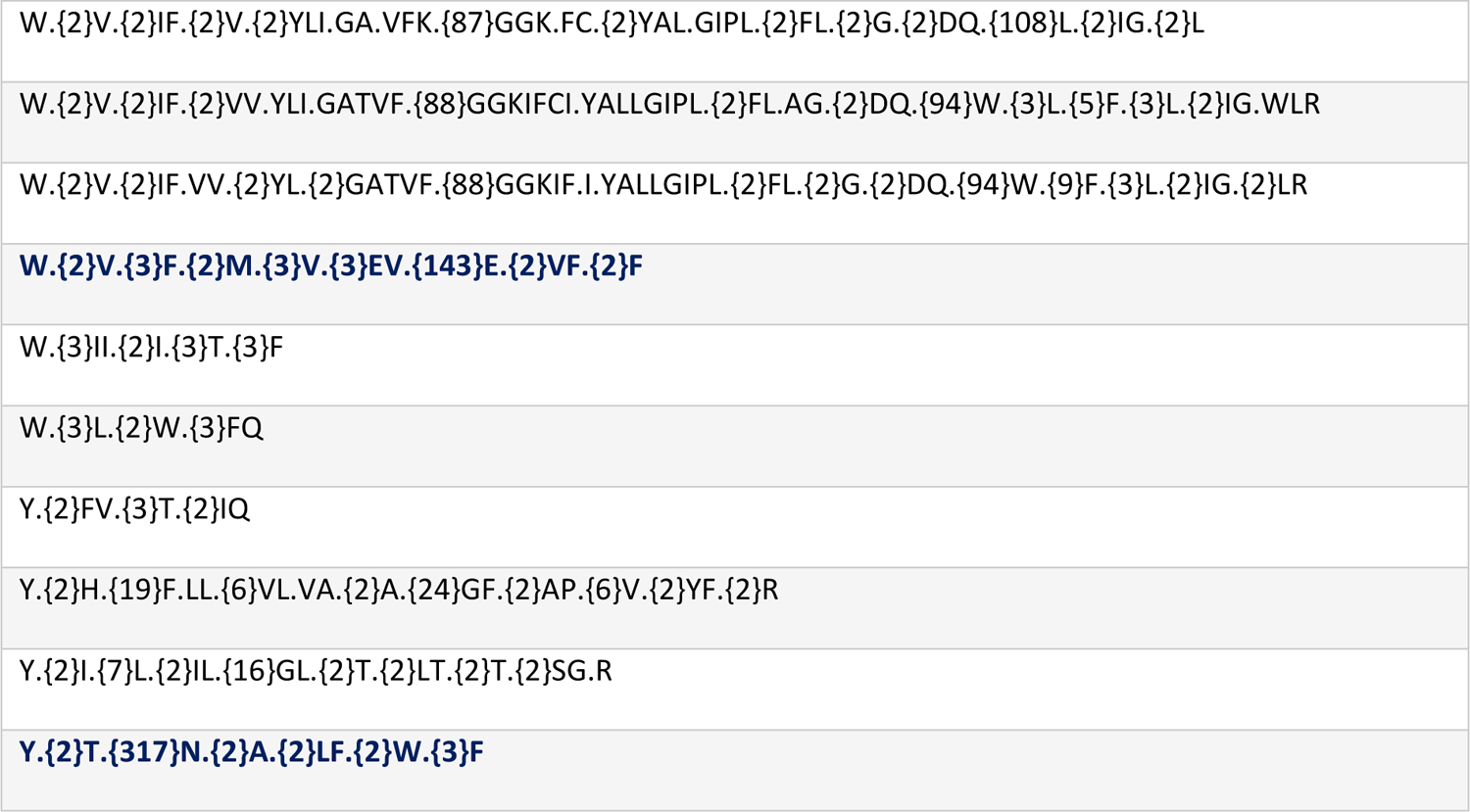
Non-redundant transmembrane α-helix PPIMem recognition motifs

PPIMem binding motifs containing the G.{3}G packing submotif are in bold orange; A.{2}T.{2}QA or A.{2}VF.LA in bold green; Aromatic.{2}Aromatic in bold blue; and L.{6}L heptad in bold gold.

In many complexes, the motifs in each subunit are identical, i.e., motif A is coupled to an identical motif B. But in other instances, the two paired motifs are different (Table S4). In this case, the complexes are usually formed of different monocatenary subunits, although, if both subunits are the same protein, but contain more than one interacting surface, the complex is a homodimer in which the contact surfaces are not the same. Of course, even when motifs A and B are the same, the complex is not necessarily a homodimer.

We observed that some amino acid residues were more favored than others in the TM recognition sites. For instance, the hydrophobic side chains Leu, Ile, Val, and Phe were the most abundant, followed by Ala and Gly. This residue distribution has been found previously (63). Leu was found more than 300 times, making about one fifth of all contact residues, like reported before (64) (Figure S1). As expected, the physicochemical properties of MPPI-binding sites are different from the exposed sites of soluble proteins. The amino acid residue abundances for helix interactions we found in the motifs match those in the literature (65, 66). Figure S2 shows, as a “heat map”, couples of contact residues at the interface for our template set of complexes, as they come out from PDBsum. We can see that the largest value corresponds to the contact between two Leu residues, followed by contact between Phe residues, and then the Leu-Phe pair. The least observed interactions include His-His for a homotypic pair and Trp-Cys for a heterotypic pair. This outcome suggests that residues tend to contact other residues sharing the same or similar physicochemical properties, and agrees with the statistics obtained for interresidue interactions in the MP Bundles DB for α-helical MPs (65). The statistical trends in the contacts imply specificity at the interface of the TM helices, as well as correlated mutations of coupled contact residues and paired motifs.

The number of TM atomic non-bonded contacts per complex covers a wide range: 11-87 for single-span TM proteins, and 8-186 for multiple-span TM complexes (Figure 3). As expected, the latter show in average a larger number of non-bonded contacts.

**Figure 3.**
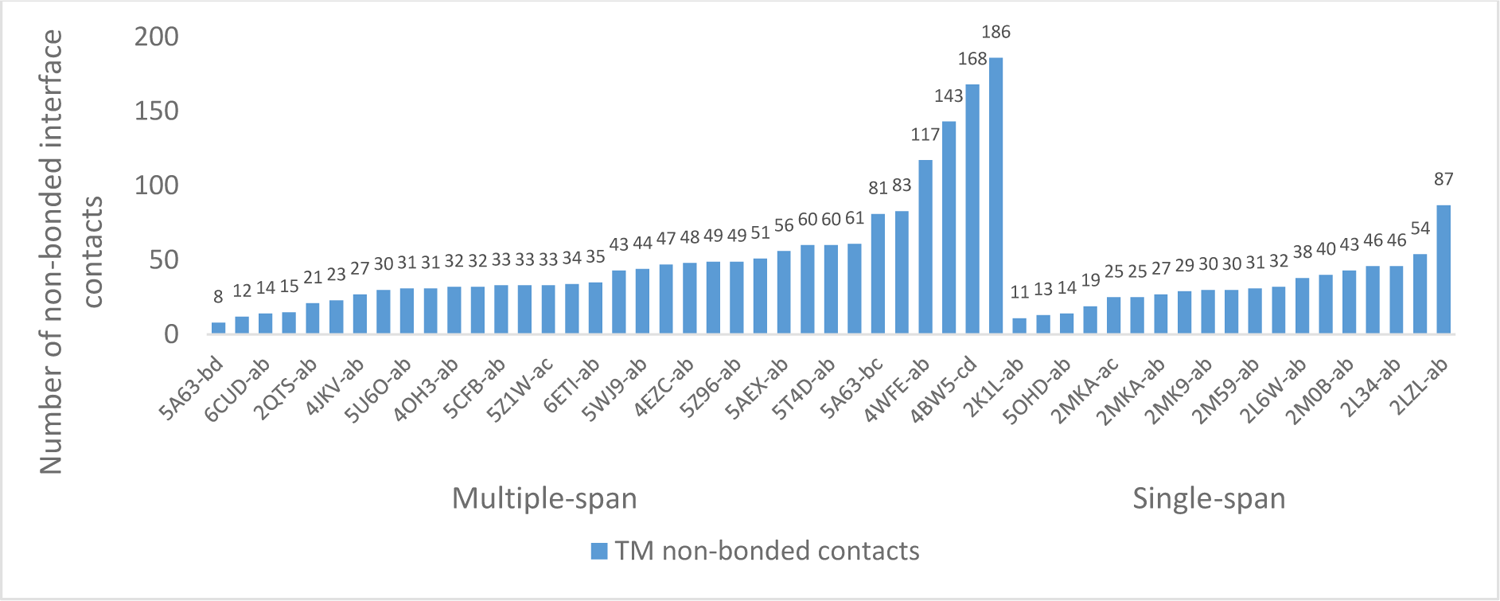
Number of PDBsum non-bonded interface contacts for each of the experimental template complex PDB structures of Table S1.

Finally, we looked at the Pfam-A set of protein families (67) to which the 63 different proteins in the template complexes of Table S1 belong. We found that the most populated families were PF07714 (Pfam family PK_Tyr_Ser-Thr) and PF00520 (Pfam family Ion_trans), with occurrences of 10/110 and 6/110, respectively. Forty-two proteins out of the 63 had only one Pfam occurrence each (not shown).

### α-helix packing motifs

Walters and DeGrado (68) have examined helix-packing motifs in membrane proteins involved in the folding of a helical membrane protein, i.e. interactions between helices within the same protein. As we are dealing with interhelical interactions between different proteins embedded in the membrane, helix-pairing motifs are not directly comparable. However, we can look at the pairing motifs of TM proteins composed of a single TM α-helical domain, such as glycophorin A, presenting a G.{3}G packing submotif (69–71), in which Gly or other small residue space four residues that mediate parallel interhelix protein interactions. Indeed, we find this small motif and its variants as part of the extracted binding motifs from our set of benchmark MP complexes (Table S1). The structures include parallel single α-helix homodimers (Table S1, PDB 2L2T and 2LOH), multiple α-helix homo-oligomers with a different motif in each component (Table S1; PDB 5AEX, chain A; PDB 5CTG, chain A). The G.{3}G submotif appears as well in the non-redundant PPIMem motifs (Table 1, in bold orange).

An antiparallel coiled-coil submotif of α-helices with Ala or another small residue in every seventh position has been termed Alacoil (72). It can be found in human aquaporin 5 (PDB 3D9S) as the PPIMem binding motif A.{2}T.{2}QA, and as A.{2}VF.LA in the TRPV1 ion channel in complex with double-knot toxin and resiniferatoxin (PDB 5IRX) of our benchmark protein set (Table S1; Table 1 in bold green).

Another specific pattern is the Aromatic.{2}Aromatic motif (73), like in F.{2}F of connexin-26 (PDB 2ZW3), of the mouse TRPC4 ion channel (PDB 5Z96), and of TRPM4 (PDB 6BCO, 6BQV). In our benchmark set, this submotif appears as part of the TrkA transmembrane domain (PDB 2N90). In the SWEET transporter (PDB 5CTG), it takes the form F.{2}Y; in polycystic kidney disease protein 2 (PKD2) (PDB 5T4D) and polycystic kidney disease-like channel PKD2L1 (PDB 5Z1W), it becomes F.{2}W (Table S1; Table 1 bold blue). Another submotif that appears very frequently is the L.{6}L heptad, analogous to the Leu zipper that mediates protein complex formation in water-soluble proteins. As this pattern is important, it is shown in Table S2 and Table 1 (bold gold).

**Table 2.** Benchmark of experimental structures of membrane protein-membrane protein complexes and their unbound components.

| Complex | Unbound | Unbound | Type |
| --- | --- | --- | --- |
| PDB code | PDB code protein A | PDB code protein B | Homo- (Hm) or<br>Hetero- (Ht) mer |
| 2LOH | 2LLM | 2LLM | Hm |
| 2QTS | 3IJ4<br>4FZ1<br>4NTW<br>4NTX<br>4NTY<br>4NYK | 3IJ4<br>4FZ1<br>4NTW<br>4NTX<br>4NTY<br>4NYK | Hm |
| 2ZW3 | 5ER7<br>5ERA | 5ER7<br>5ERA | Hm |
| 3SYQ | 3SYA<br>3SYC<br>3SYO<br>3SYP<br>4KFM | 3SYA<br>3SYC<br>3SYO<br>3SYP<br>4KFM | Hm |
| 4JKV | 4N4W<br>4O9R<br>4QIM<br>4QIN | 4N4W<br>4O9R<br>4QIM<br>4QIN | Hm |
| 4NEF | 4OJ2 | 4OJ2 | Hm |
| 4WO1 | 2L34<br>4WOL | 2L34<br>4WOL | Hm |
| 4X5T | 2M6B | 2M6B | Hm |
| 5A63 | 2N7Q<br>2N7R | 2N7Q<br>(for Q92542)<br>2N7R (for Q92542) | Ht |
| 5O9H | 6C1Q<br>6C1R | 6C1Q<br>6C1R | Hm |

PPIMem reveals original packing motifs through its non-redundant consensus motifs, such as the frequent motif A.{2}(A/L/V)(L/F) (Table 1). The following motifs with Ile are also well populated: for position i-1, (F/L/V/W)I; for positions i-2 and i-1, (F/L)CI; for positions i+3 and following, I.{2}(A/L/V), I.{3}(L/V), I.{5}I, and I.{6}A. PPIMem also uncovers Ile composite motifs, like I.{2}(A/F)I.{2}(T/L), and I.{3}I.{3}L. The most frequent motifs are those beginning with Leu: L.{2}(A/F/G/I/L/N/T/V)(A/G/F/I/L/V), and L.{3}(F/V). Leu composite motifs include L.{2}(F/I)L.{2}G, L.{2}IF.{2}LL.{2}F, L.{3}L.{2}FF, and L.{3}L.{2}INP.{2}L.{3}V. The occurrence of these patterns is reflected in the interhelical contacts between proteins of the Ile-Ile, Leu-Leu and Ile-Leu residue pairs (Figure S2). The composition of these residues shows, as if it was necessary, that hydrophobic residues are enriched at the interface between MPs.

A statistical analysis of amino acid patterns in TM helices (71, 74) results in 30 over-represented (p <<<, odds ratio >1), and 30 under-represented (p <<<, odds ratio <1) pairs. Thus, G.{3}G (GG4 in Senes et al. notation) is the most significant pair among over-represented pairs. Indeed, this and other significant over-represented pairs (21 out of 30), such as I.{3}I, G.{3}A, IG, I.G, V.G, I.{3}V, IP, V.{3}V, V.{3}I, AV, and G.{2}L, have their equivalents in non-redundant PPIMem motifs (Table 1). On the other hand, several under-represented pairs (10 out of 30) are in fact absent from PPIMem motifs in Table 1, such as I.I, F.{3}I, and I.{3}G. Moreover, several most significant over-represented triplets of Senes et al. appear in our motifs, lending support to our template-based approach.

We then wished to look at the number of UniProtKB ACs resulting for each number n of contact residues for the biological species recorded in the PDB and UniProtKB. As seen in Figure 4, the count occurrence of contact residues in the motifs is largest for n = 7 for both protein A and protein B of the interacting pair. The quasi-periodicity of the interface recognition residues reflects a spatial arrangement corresponding to a heptad α-helical pattern. This is true also for motifs separated by values of the linker residues ≥4. For example, the long motif DWN.{2}IA.{2}V.{2}LI.{24}F.{3}F.{2}A.{2}YLV.{2}FM (Table 1) contains two motifs of 14 and 17 residues, separated by 24 linker residues. A motif may attain a total of 20 residues, like in E.{2}TL.VGF.IL.{2}SL.{2}TY (Table 1), possessing enough length to span a bilayer. Our motifs therefore may be rather long, implying residue correlations beyond the pair level. Moreover, an additional information is given by the PPIMem motifs: the residues that compose them may be not only on the same face of the helix but are interfacial residues. The corresponding statistics for *H. sapiens* show similar trends: small motifs with six to seven buried contact residues are the most abundant (peak at n = 7, combined number of redundant proteins A and B, 9655). The range of contact residues in the motifs was 6-32. As the number of contact residues increased in the motifs, the number of hits decreased drastically, with only one to four predictions for both proteins in the 18-33 contact residue range (Figure 4). These findings suggest a limited set of binding motifs in nature.

**Figure 4.**
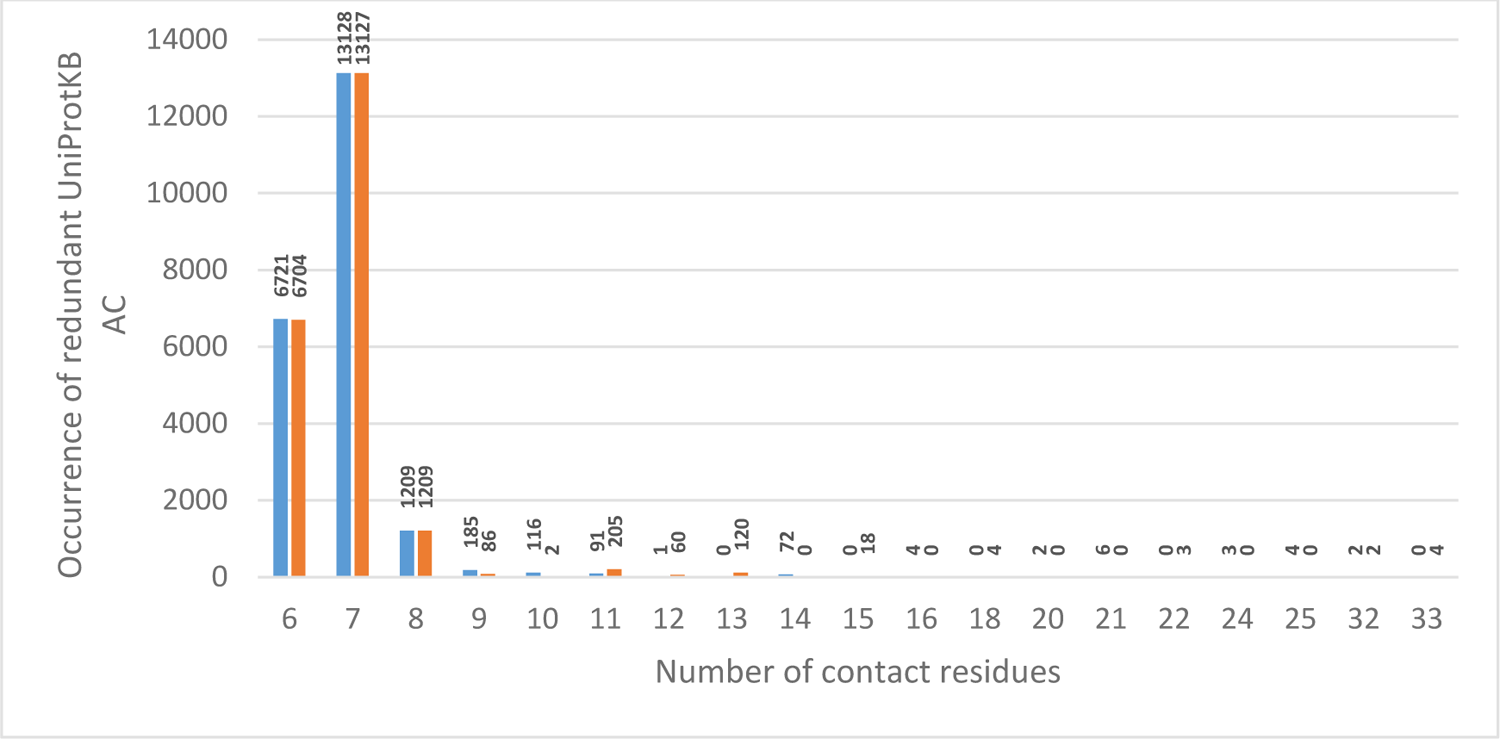
Occurrence of (redundant) UniProtKB ACs as a function of interface contact residues for Proteins A (blue) and corresponding Proteins B (orange). Mutation rate 0-20%. Organisms extracted from the PDB and UniProtKB.

The protein-protein docking benchmark 5.0 PPDB (75) assembles non-redundant high-resolution complex structures, for which the X-ray or NMR unbound structures of the constituent proteins are also available (https://zlab.umassmed.edu/benchmark/). However, none of the complexes in PPDB v5.0 correspond to MPs. On our side, from our 53 non-redundant template structures of MP-MP complexes, we were able to extract a subset of them along with the unbound structures of their components to define a benchmark. This benchmark is made up of ten sets of structures (Table 2).

The first column shows MP complexes of known structure. The 2^nd^ and 3^rd^ columns list the experimental structures of the isolated chains that compose them. The 4^th^ column gives the type of oligomer (homo- or heteromer). PDB structure 5A63 represents the sole heteromer of the benchmark. All the other complexes are homomers. Note: This table is derived from Table S3, Column C.

When comparing our benchmark to the “Dimeric complexes” table of the Membrane Protein Complex Docking Benchmark, MemCplxDB (76), we only recover the PDB 5A63 complex. The reason is that MemCplxDB shows many interactions between MP and non-MP, soluble proteins (antibodies, peripheral proteins, etc.), which we do not deal with. MemCplxDB includes interactions as well between oligomers within a multimer complex, and prokaryotic MPs of β-barrel structure. Our benchmark represents thus a gold-standard set of positives for integral-to-membrane proteins interacting through their α-helical TM segments. As the 3D structures of more TM complexes appear, the benchmark will grow and could serve for a machine learning approach of prediction of membrane PPIs.

### Predicted interactions

The number of PPIMem-predicted membrane heteromer interactions for the 39 species dealt with is 21, 544 among 1, 504 MPs. The homodimers are thus 1, 504 in number and represent 6.5% of all complexes. Of the total heteromer interactions, 9, 797 among 417 MPs correspond to *H. sapiens*, the homodimers representing 4.1%. PPIMem predicts interactions thus for 417 human genes, including many disease genes. The Mendelian Inheritance in Man (MIM) number contains information on the known mendelian disorders caused by variants affecting the gene represented in the entry and focuses on phenotype-genotype relationships (https://www.omim.org/). Table S5 shows that 101 out of 417 PPIMem MPs are involved in human disease according to MIM. This set provides 83 PPIMem MPs extracted from complexes with nil mutations in their binding motifs. We looked at their missense variants in the index of human variants curated from literature reports in UniProtKB (https://www.uniprot.org/docs/humsavar), and focused on the following categories as defined by the ACMG/AMP terminology (77): likely pathogenic, pathogenic or of uncertain significance. Table S5 shows that several mutants involved PPIMem interface contact residues belonging to TM α-helices, suggesting a destabilization of the complex as the molecular basis of the disease.

At different mutation rates, we defined and identified many potential recognition sites and novel binary complexes. As mentioned in the S&M section, we built a consensus amino acid contact motif for all the matched sequences of a given binding site in the 0-20 % mutation range considering the contact residues only. This led us to detect conserved amino acid residues among the contact residues of the binding motifs. The most prevalent consensus motif A found was AV.{2}GL.{2}GA.{2}L, illustrated by the instance sequence ^423^AVFSGLICVAMYL^436^ of the sodium-dependent proline transporter (UniProtKB Q99884) at a 15% mutation allowance (Figure 5a). For consensus motif B, the same motif was the most frequent (Figure 5b). The least frequent motifs are characterized by hefty linker segments between the contact residues and thus by substantial sequence lengths. An example is motif B: LL.AS.{90}LV.EA.FAI.NI.{2}S.{2}L.{3}F.{19}I.{100}I, found in the short transient receptor potential channel 4 (UniProtKB Q9UBN4). In general, the interface contact residues are part of conserved sequences of TM regions.

**Figure 5.**
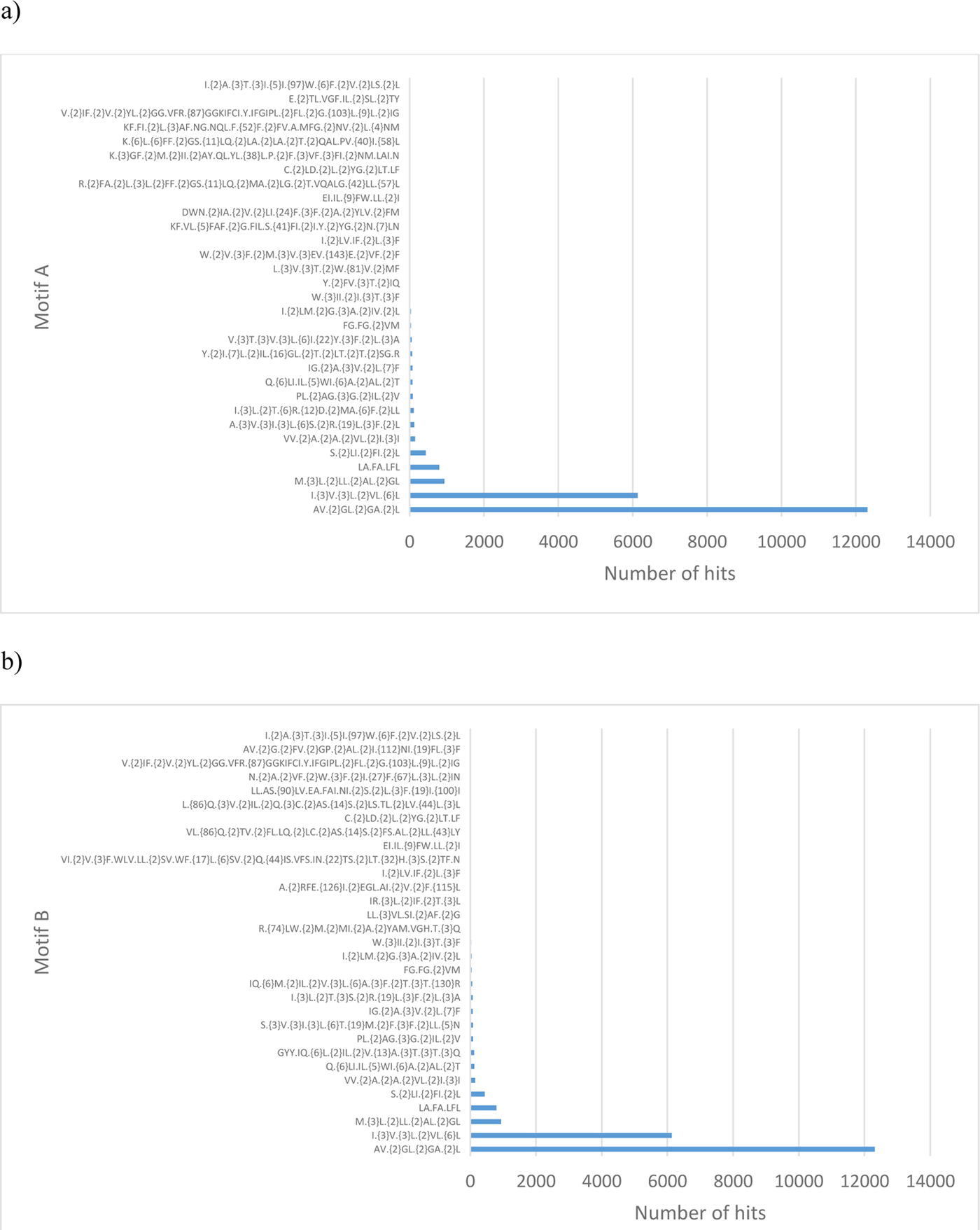
Number of hits per motif for all species. a) motif A, b) motif B. Mutation rate 0-20%.

We cannot know at this stage whether or not the amino acids composing our binding motifs represent hot-spots, i.e., those that contribute to the binding free energy (78).

Mutation rates of 0 % for motif A and motif B result in those proteins whose contact residue sequences conform exactly to the consensus motifs. The PPIMem outcome gives 79 entries of hetero-oligomer complexes across species, of which 25 are *H. sapiens’*. Retaining motif A only, PPIMem predicts 346 interactions for *H. sapiens* at a mutation rate of 0%. For the mutations rates 5, 10, 15, and 20%, the number of interactions is 16, 4439, 4272, and 724, respectively.

As mentioned, we derived a consensus sequence from multiple alignments of the contact residues of all sequences found for a given motif. Thus, for example, the consensus sequence resulting from up to a 20% mutation rate of the VV.{2}A.{2}A.{2}VL.{2}I.{3}I motif of length 19 is shown in Figure 6, leading to VVX_2_AX_2_AX_2_VL as the fingerprint of the binding motif.

**Figure 6.**
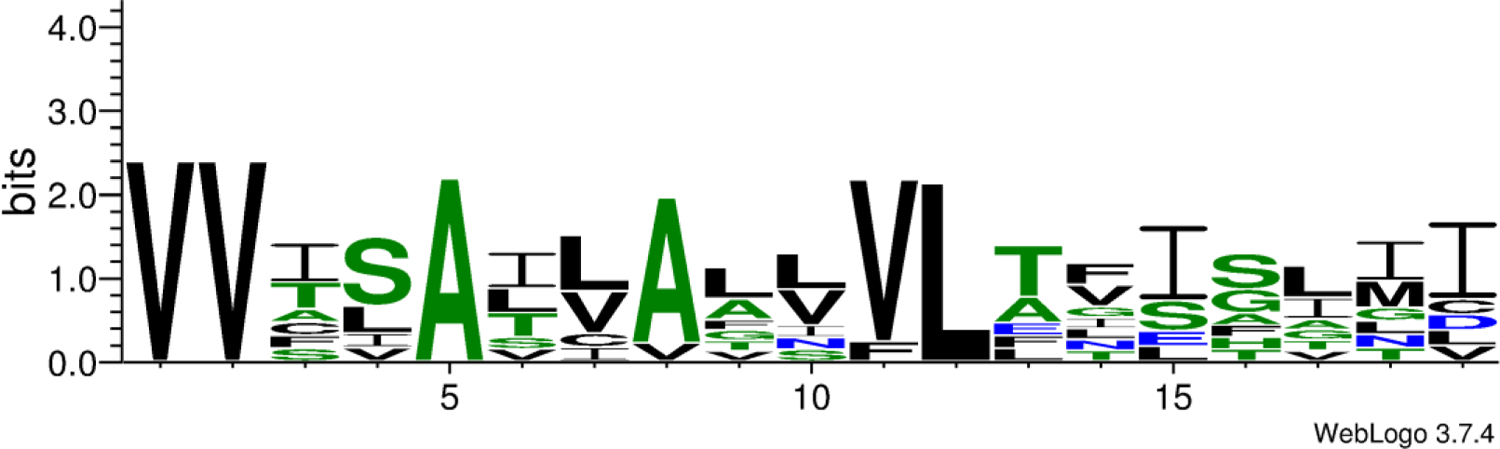
Multiple sequence alignment WebLogo representation (https://weblogo.berkeley.edu/) showing the consensus sequence for the VV.{2}A.{2}A.{2}VL.{2}I.{3}I binding motif of length 19. Black denotes hydrophobic residues; green, polar residues.

### PPIMem and other datasets

The PPI template-based prediction algorithm and server PRISM2.0 (79) also uses the 3D structural similarity of interfaces with respect to templates to propose other protein complexes. PRISM is not specific for MPs and requires the 3D structure of the targets to propose an interaction, whereas PPIMem is specific for MPs and does not require the 3D structure of the subunits composing the putative complex. Thus, when having an interface template corresponding to an MP, PRISM may propose not only TM protein complexes but also globular protein complexes. As a consequence, many of our MP template interfaces are absent in the PRISM dataset.

Although most datasets based on experimental approaches cover the entire human proteome, again, MPs are under-represented (for an overview of the major high-throughput experimental methods used to detect PPIs, see (80, 81)). Thus, the experimental work of Rolland and colleagues on the human interactome network (24) found only 41 interactions between MPs. Twenty-eight of these proteins were found to interact in the IntAct DB. Nevertheless, none of the interactions we extracted from the structural PDBsum DB were found among the 41 interactions above. It seems that some of these interactions bear between the juxtamembrane regions of the MPs reported by Rolland et al. for MPs. We did not find either any of our predictions in their results. In the HI-III-20/HuRI updated dataset (11), none of the high-quality binary PPIs are MPs (log_2_ odds ratio < 0, Extended Data Figure 7a of Luck et al.)!

**Figure 7.**
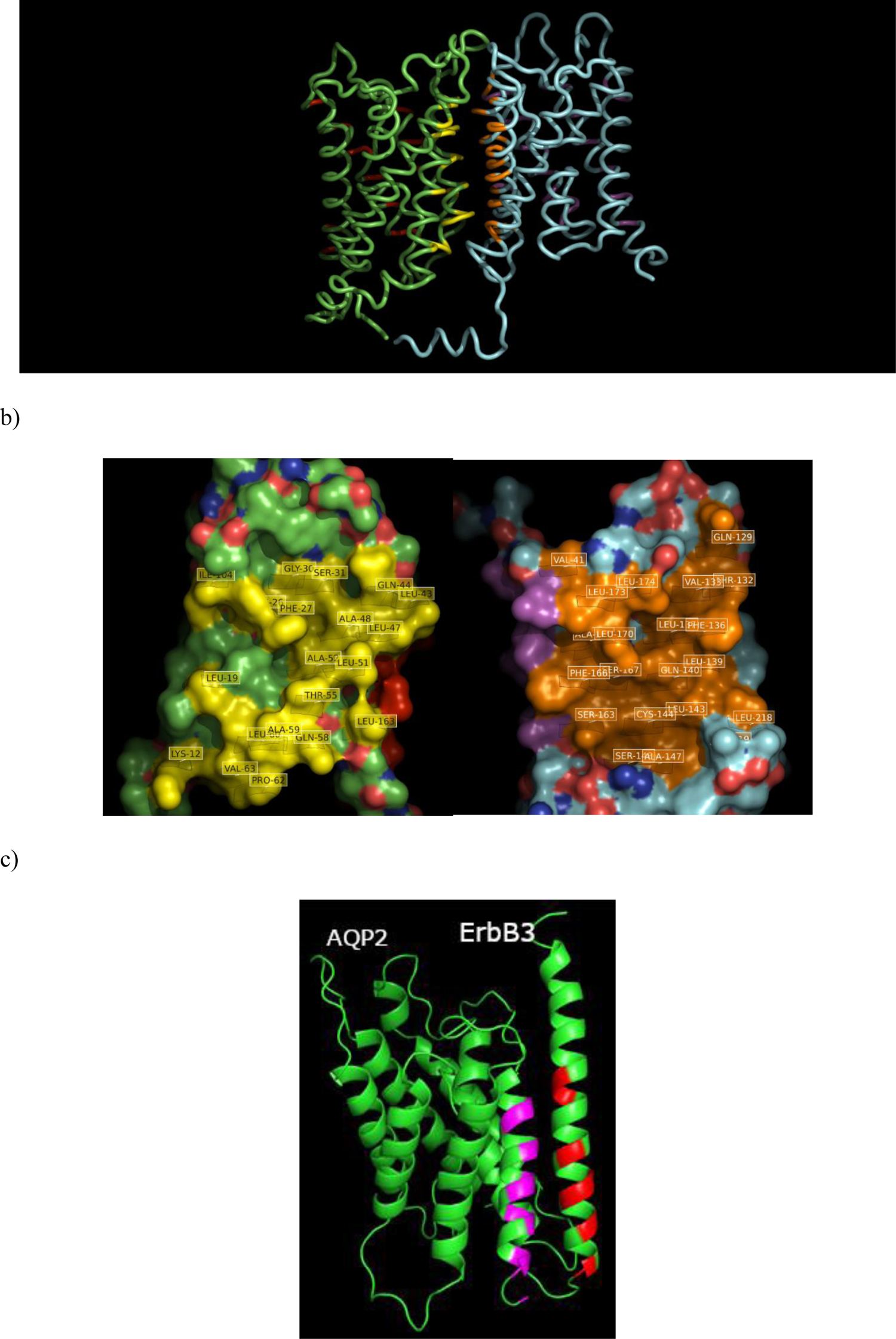
Low-resolution cartoon structural model of predicted PPIMem MP-MP complexes obtained by molecular docking. a) H. sapiens’ protomers of aquaporin 5 (UniProtKB P55064, PDB 3D9S, green) and aquaporin 2 (P41181, 4NEF, cyan) in complex, with PPIMem interface residues in yellow and orange, respectively. The binding motif for the former is K.{6}L.{6}FF.{2}GS.{11}LQ.{2}LA.{2}LA.{2}T.{2}QAL.PV.{40}I.{58}L, whereas that for the latter is R.{2}FA.{2}L.{3}L.{2}FF.{2}GS.{11}LQ.{2}MA.{2}LG.{2}T.VQALG.{42}LL.{57}L. Regions in red and magenta in the rear side of the molecules correspond to a second binding motif; b) the same complex in a solvent-accessible surface representation in which each chain has been rotated 90° towards the viewer, revealing the contact interface residues (labeled) - yellow for P55064 and orange for P41181; c) 3D model of the docking complex between R. norvegicus’ protomers of aquaporin 2 (P34080) and the receptor tyrosine-protein kinase ErbB-3 (Q62799) in which the AV.{2}GL.{2}GA.{2}L binding motif, present on both protein surfaces, was used to direct the docking. AQP2 interface residues are in purple and ErbB-3 residues in red.

When relating our PPIMem predictions at a 0-20% mutation rate for human proteins to several datasets like FPClass, BioPlex 3.0, MENTHA, IID, HMRI, and BIPS, we find several consistencies, listed in Table 3.

**Table 3.** Predicted interactions common to PPIMem (0-20% mutation rate, *H. sapiens*) and several datasets

Common interactions
| UniProtKB AC A | UniProtKB AC B | FPClass total score |
| --- | --- | --- |
| P18507 | P47870 | 0.4029 |
| P34903 | Q16445 | 0.2915 |
| P47869 | Q16445 | 0.2738 |
| P47870 | P48169 | 0.2901 |
| P47870 | Q16445 | 0.2837 |
| P48169 | P47870 | 0.2901 |
| P48169 | Q16445 | 0.2786 |
| Q01814 | Q16720 | 0.3235 |
| Q16445 | P47870 | 0.2837 |
| Q16445 | P48169 | 0.2786 |
| Q16720 | Q01814 | 0.3235 |

Common interactions
| **UniProtKB AC A** | **UniProtKB AC B** | **BioPlex**<br>**Probability of**<br>**interaction** |
| --- | --- | --- |
| Q9Y5G3 | Q9Y5G2 | 1.0 |
| Q9UN71 | Q9Y5G0 | 1.0 |
| P47870 | P28472 | 0.99 |
| P51795 | P51790 | 0.99 |
| Q01814 | P23634 | 0.99 |
| Q13507 | Q9HCX4 | 0.99 |
| Q16445 | P28472 | 0.99 |
| Q9H313 | O95180 | 0.99 |
| Q9H313 | O95427 | 0.99 |
| Q9H313 | Q8IYS0 | 0.99 |
| Q9H313 | Q96D96 | 0.99 |
| Q9Y5G2 | Q9UN71 | 0.99 |
| Q9Y5G2 | Q9Y5F8 | 0.99 |
| Q9Y5G2 | Q9Y5F9 | 0.99 |
| Q9Y5G2 | Q9Y5G0 | 0.99 |
| Q9Y5G2 | Q9Y5G1 | 0.99 |
| Q9Y5G3 | Q9UN71 | 0.99 |
| Q9Y5G3 | Q9Y5G0 | 0.99 |
| Q9Y5G3 | Q9Y5G1 | 0.99 |
| P34903 | P31644 | 0.99 |
| Q01814 | Q16720 | 0.99 |
| P36021 | Q8IYS0 | 0.99 |
| Q9H313 | Q9UP95 | 0.99 |
| Q9UN71 | Q9Y5G1 | 0.99 |
| Q9UNQ0 | O15126 | 0.97 |
| Q9Y5G0 | Q9Y5G1 | 0.89 |
| Q8NGN1 | Q9UP95 | 0.88 |
| P25090 | O95427 | 0.86 |
| Q9UBK5 | Q96D53 | 0.86 |
| Q96FT7 | Q68CP4 | 0.84 |
| P47871 | Q96D53 | 0.83 |
| O75311 | P28472 | 0.78 |
| O75311 | P28472 | 0.78 |
| O75311 | P28472 | 0.78 |
| Q15758 | Q8TCT6 | 0.78 |

This table comparing our PPIMem predictions to experiments represents a validation of our method. For example, PPIMem predicts the interaction between isoform 2 of the human γ-aminobutyric acid receptor subunit β-3 (GABRB3, P28472-2) and subunit α-6 (GABRA6, Q16445). In the BioPlex 3.0 dataset, the interaction is detected with a probability >0.99. The ComplexPortal DB https://www.ebi.ac.uk/complexportal/home) reports the same interaction as Complex AC: CPX-2164 and CPX-2951. If we consider that BioPlex reports 2, 000 MP interactions, then the present non-specific PPIMem prediction rate is of ∼ 2%, considering the 35 interactions described above and the fact that the BioPlex and PPIMem sets are independent, i.e., there is no intended intersection between them.

In the following, we illustrate our data for *H. sapiens* with other datasets:

– The MENTHA experimentally-determined direct protein interactions DB presents also the P28472-Q16445, as well as the P28472-P47870 (GABRB2) interaction.
– The IID DB validates experimentally the Vascular endothelial growth factor receptor 2 (P35968) – receptor 3 (P35916) interaction.
– The HMRI DB, which seems to list only heteromers and for which not all the interactions are between MPs, shows a correspondence for the heterotypic pair TYROBP-KLRC2 (p value = 0, 034341).
– BIPS (82) predicts putative interactions and is based on sequence homology between proteins found in PPI DBs and templates. We find several correlations between BIPS and PPIMem. For instance, we propose an interaction between T-cell surface glycoprotein CD3 ζ chain (P20963) and high immunity immunoglobulin ε receptor subunit γ (P30273). BIPS predicts a similar pair between T-cell surface glycoprotein CD3 ζ chain (P20963) and low-affinity immunoglobulin γ Fc region receptor III-A (P08637).

Finally, as the benchmark set is so small to serve as a training set, we could not evaluate quantitatively interface predictions using standard criteria (ROC plot, TP/FP, etc.).

### Interaction networks

Networks link overlapping pairs of proteins, from which it is possible to propose multimer complexes if the binding sites are independent and non-overlapping. The architecture of a network reflects biological processes and molecular functions. Thus, from the predictions, *de novo* connections can be found, linking the network to a disease pathway, and proposing innovative possible cellular roles for some of the complexes. We illustrate below a subnetwork of PPIMem-predicted MPPIs for *H. sapiens*:

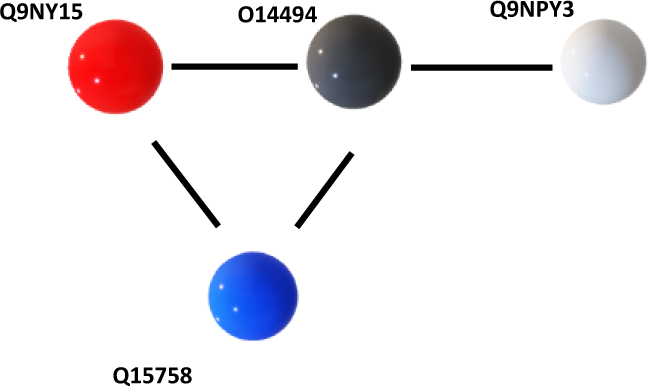

Q9NY15 (*STAB1*) Stabilin 1; O14494 (*PLPP1*) Phospholipid phosphatase 1; Q15758 (*SLC1A5*) Solute carrier family 1 member 5 or Neutral amino acid transporter B(0); Q9NPY3 (*CD93*) Complement component C1q receptor.

In support of the proposed subnetwork, we found that all four MPs were present in two tissues of *H. sapiens’* — adipose tissue (major) and breast (minor) as reported in the Human Protein Atlas (https://www.proteinatlas.org/). Furthermore, a *CD93* - *STAB1* interaction in humans has been reported in the String DB of PPI networks, and *CD93* and *PLPP1* are co-expressed in *G. gallus* (83).

### Negative interactions

The importance of recording negative results of PPI assays in interatomic DBs, i.e., those indicating the tested proteins do not interact, has been raised (84). However, their identification is less straightforward. This feature should eventually lead us to define a set of true negative interactions with the goal of training a predictor of MPPIs as the sampling of negatives is crucial for optimal performance (85, 86). Looking for negative interactions in the IntAct DB for our PPIMem proteins, we find two negative interactions of two PPIMem proteins, but with two non-MPs. Analogously, the Negatome datasets (87, 88) and Stelzl (89) compile sets of protein pairs that are unlikely to engage in direct physical interactions. We observed that spanning the Negatome dataset with our predicted positive interactions for *H. sapiens* gives no results. In other words, the Negatome DB does not report any of our complexes as a negative interaction. Conversely, several MPs in the Negatome set are absent from PPIMem. As this information is not conclusive, we will continue to probe our dataset as more negative interaction data become available.

### Molecular docking simulations

To support the predictions from a structural point of view, we selected several MP pairs for molecular docking simulations for which PPIMem predicted an interaction based on the interface epitopes. To begin with, we took human AQP5 (P55064) and AQP2 (P41181). In this case, the crystal structures exist for both MPs (PDBs 3D9S and 4NEF, respectively). For the docking simulation, we took one of the PPIMem predicted patterns for AQP5 as the binding site (K.{6}L.{6}FF.{2}GS.{11}LQ.{2}LA.{2}LA.{2}T.{2}QAL.PV.{40}I.{58}L; Table S1), and one of the predicted patterns for AQP2 (R.{2}FA.{2}L.{3}L.{2}FF.{2}GS.{11}LQ.{2}MA.{2}LG.{2}T.VQALG.{42}LL.{57}L; Table S1). It is interesting to note that both motifs show contact residues separated by regions with many wildcard residues (40 and 58 for AQP5; 42 and 57 for AQP2). This is a direct consequence of the spatial distribution of the contact residues that may belong to different helices but still be involved in the same intermolecular interaction. Moreover, not all residues of a given exposed transmembrane helix are necessarily contact residues, but only a few, such as one Ile and one Leu at the end of the AQP5 motif and belonging to different helices. The other example are the three Leu at the end of the AQP2 motif, two of which belong to one helix and the third one to another helix. That is why the primary structure of the PPIMem binding motif codes also for the tertiary structure. Figure 7a shows a heterodimer among the top 10 GRAMM-X docking predictions that involves exactly the membrane-exposed interface regions of each MP. In Figure 7b each chain has been rotated 90° towards the viewer to reveal the contact interface residues.

On another hand, we selected the predicted rat AQP2 (P34080)-ErbB-3 (Q62779) pair, both subunits interacting through the AV.{2}GL.{2}GA.{2}L pattern on both protein surfaces. Since the experimental structures are unavailable for either MP, we homology-modeled them from human AQP2 (PDB 4NEF) and human ErbB-3 (PDB 2L9U), respectively. The sequences of both aquaporins (rat and human), are more than 30% identical in the TM region, just like both ErbB-3’s; the resulting individual 3D models are thus highly reliable. Figure 7c shows that the modeled rat AQP2-ErbB-3 heterodimer respects the query interface, suggesting that the complex is viable. In both complexes, the contact residues are indeed at the interface of the complex and may lead to its formation. a)

Lastly, in the PPIMem user interface, when Valid = 2, the 3D structures of each of the isolated protomers making up a putative complex are available. It is therefore possible to perform the directed docking implemented above to obtain 3D structures of any one of the corresponding 67 complexes (63 for *H. sapiens*). The PDB IDs can be found in the UniProtKB DB through the UniProtKB link in our database.

## DISCUSSION

In this work, we developed PPIMem, an interface residue template-based protocol destined to predict at large scale MP complexes through binary interactions among their α-helical TM segments. PPIMem is a model-driven biological discovery tool to be queried for the discovery of verified and potential interactions, and for obtaining varied types of necessary information about them. It contains TM oligomerization recognition sites based on the assumption that homologous structural interacting motifs always interact in similar orientations and with equivalent interfaces. In this work, we report exclusively interactions taking place in the eukaryotic plasma membrane interactome in which the binding sites specifically involve α-helical TM regions. PPIMem predicts thus an α-helix — α-helix TM protein MP interactome with thousands of *de novo* interactions, including multiple recognition sites, i.e., MPs with more than one interface, important for multimer formation. The obtained sequence motifs identify homo- and heterodimer interface amino acid residues that represent the first step to generating higher-order macromolecular edifices. Albeit our benchmark set is small (Table 2), it is of high quality and does not mix distinct types of organisms or membranes as other sets do. We have not assessed the effects of intramembrane mutations in the structure or function of the MPs, as the goal of our approach implementing different degrees of mutations was that of finding other MPs with homologous interfaces and thus forecasting new complexes.

The uniqueness of the PPIMem approach resides on our focusing on the local, membrane-exposed interface residues, largely responsible of the formation of a complex between transmembrane proteins. The resulting MP interactome represents “first draft” predictions and contains 21, 544 unique entries for all species dealt with, of which 9, 798 for *H. sapiens*. The considerable number of protein partners we uncover suggests that even distantly related TM proteins make use of regions of their surface with similar sequences and arrangements to bind to other proteins. The predicted interaction partners can lead to generating low-resolution 3D structures for several predicted complexes. This shows that complex formation is feasible as the interacting surfaces of the individual proteins manage to face each other in the docked complex and provides a partial validation of the method.

There are many caveats of any analysis comparing PPIs from different sources (90), as there are large discrepancies and dramatic differences in the content between experimental PPI data collected by the same or different techniques, making our attempt to compare our predictions to experimental data more haphazard. Indeed, the intersections between various interaction maps that employ altogether diverse approaches are very small (91, 92). For example, even though the interactions in our template set exist in the PDB structure, most of them do not show in PPI databases. Moreover, the presence of orthologs makes the research more cumbersome. Thus, the comparison between our predicted set of TM complexes and other datasets is necessarily qualitative. Disagreements occur also because MPs are not well probed in experiments. Because of all these factors, we do not have a large, validated dataset to compare to and assess the FPs. Still, to reduce the FPs we considered only motifs with six or more contact residues. It is neither possible to perform proper cross-validation, as our initial starting set (the template complexes) includes only true interactions, and no negative interactions. In few words, we cannot quantify FP, FN and TN. Nevertheless, we found several of our suggested interactions in different experimental PPI DBs, representing part of the validation approach.

Complementary to the sequence-based co-evolution PPI prediction methods (7, 93–95), our approach encodes 3D into 1D and thus adds the spatial dimension to a given MP interactome. This may lead to pioneering biological hypotheses concerning PPIs at the membrane level, to genotype-phenotype relationships, to investigation of the effect of pathological mutations on the interaction between MPs, and to propose molecular mechanisms of action. Recovering PPIMem predictions for human MP complexes in several experimental PPI datasets obtained by different methods (BioPlex 3.0, MENTHA, IID, HMRI) highlights the pertinence of our approach.

Because of their number and diversity, the higher-order structures and interfaces generated in this work represent potential pharmacological targets for the discovery of modulators targeting the intramembrane protein-protein interface. These modulators include membrane insertable, metabolically stable, non-toxic small-molecule active MPPI inhibitors, stabilizers and allosteric binders that influence *in vivo* the assembly of intact MPs involved in various diseases. Examples of these modulators are exogenous peptides and peptidomimetics (96–101). Indeed, many of the MPs in the predicted complexes are involved in a variety of diseases as deficient or enhanced oligomerization is associated with diseases (102, 103). Additionally, our protein interaction data imply direct interactions and can lead to the construction of protein interaction networks. Thus, our results may help understand biochemical pathways and the crosstalk between them, including transactivation; they may suggest as well potential drug targets, and may aid in understanding the functional effects of mutations at the interface (5). The PPIMem approach should be useful as well for the design of MP structure.

The experimental validation of the predicted MP-MP interactions could be tested with techniques such as FRET (104), or by our exclusive Microtubule Bench approach (105). By applying machine learning methods, the PPIMem method can be improved by insuring that the MPs belong to the same developmental stage, tissue, cell type, site of expression, reaction and metabolic pathways, display functional similarity, and do not show a gene distance of more than 20 (106).

Incidentally, the PPIMem algorithm can be applied to soluble proteins and to other cell membranes (mitochondria, nucleus, endoplasmic reticulum, Golgi apparatus) and across the tree of life, provided the 3D structures of the corresponding protein complexes become available. Our methodology can equally be extended by introducing side-chain and main-chain h-bond and electrostatic information at the interface, important for MPPIs (107). Other applications of our approach include homologous protein networks in other organisms.

### Data availability

Figure S3 shows a screen capture of PPIMem’s first page. The corresponding source code and instructions for running the prediction algorithm have been deposited at github.com/PPIMem. The database with the annotated predicted interactions is implemented as a web application that supports sorting and filtering. The output data can be downloaded as a csv file and the predictions can be accessed at https://transint.shinyapps.io/transint/.

Several pertinent notes are to be found in the Supplemental Material.

## Acknowledgments

We thank Prof. D. Pastré for critical reading of the manuscript.

