## Supplemental material for "Getting to know each other: PPIMem, a novel approach for predicting transmembrane protein-protein complexes"

TITLE

### NOTES

Our approach has some limitations derived from the inaccuracies and incompleteness of the original DBs, like the PDB, containing *in vitro* multimers in different media that modulate the oligomerization interface (1); however, the reported complexes could exist in living organisms and be obligate since there are no enzyme-peptide complexes. On the other hand, there may be allosteric mechanisms, as well as ligand-induced binding effects. Neither do we consider the TM interaction driven by extra-membrane interactions, just like the possible control of MPPIs by post-translational modifications (2). Also, we examine only the isoforms of the PDB template structures and those recovered from UniProt canonical sequences, and we assume the MPs to be rigid bodies; fortunately, the conformational changes upon complexation are known not to be major for the TM domains. Within the limits of our assumptions and of our approach, our proposed amino acid residue recognition sites and corresponding molecular complexes can address the possible effects of natural or artificial mutations on PPIs, as well as protein function and disease associations. Thus, the potential interacting MPs identified in our DB may be used as candidates to be validated experimentally.

Finally, because the MPPI data set may be noisy, we intend to employ next machine-learning techniques to design a more exact predictive model, allowing us to better estimate complexes between MPs.

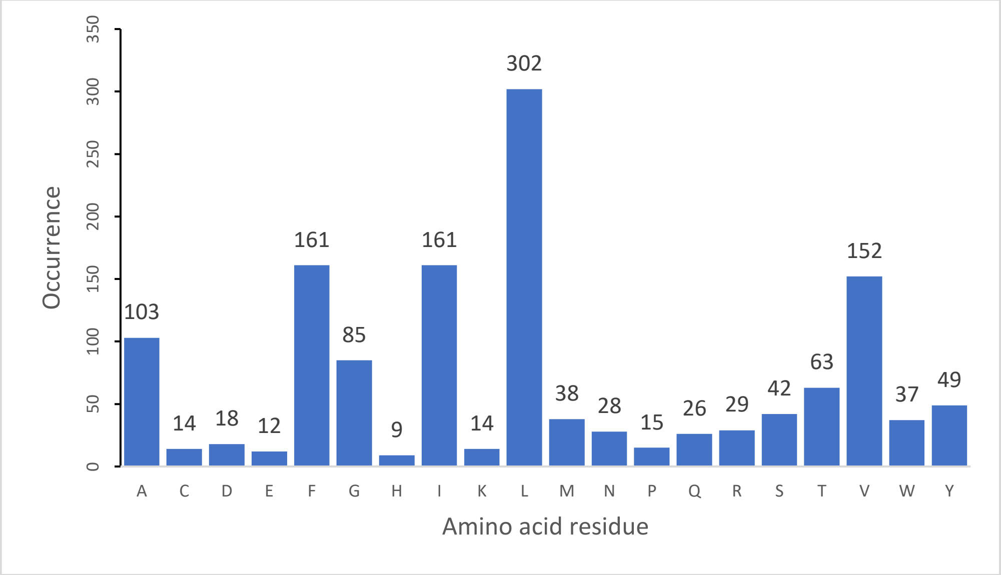

#### Figure S1

Frequency of amino acids in the extracted motifs from the PDBsum contact maps at the interfaces of MP complexes.

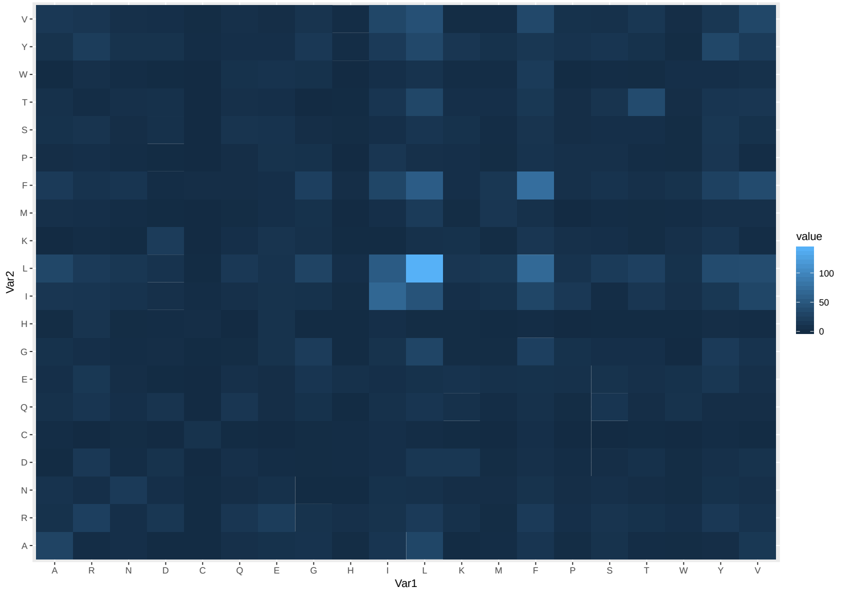

#### Figure S2

Heat map of pairs of contact residues at the MP-MP interface of the template complexes. Amino acid residue names are represented by the one-letter code.

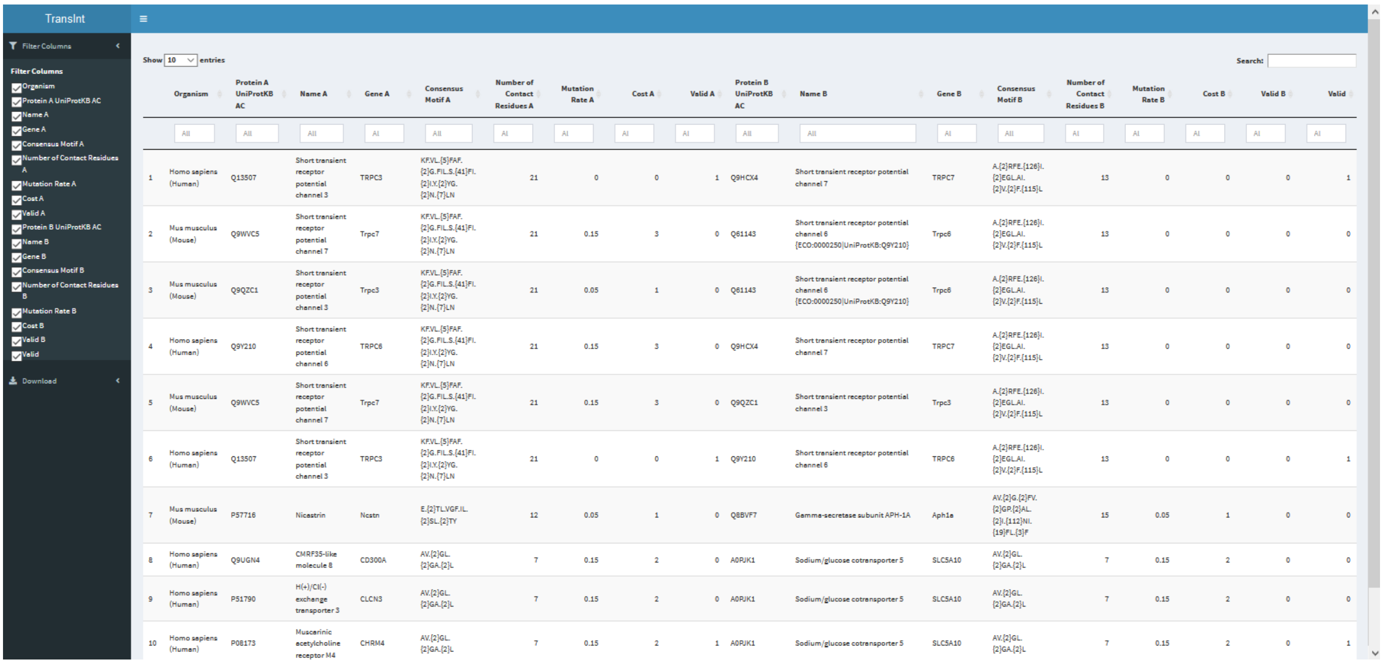

#### Figure S3

Screen caption of the PPIMem database webpage. The menu on the left allows filtering the columns on the right. The first column corresponds to the Organism name. The next eight columns are for Protein A, and represent its UniProtKB AC, its Name, its Gene, its contact Consensus Motif in regular expression format, the Number of Contact Residues in the Consensus Motif, the applied percent Mutation Rate, the Cost parameter (number of allowed amino acid mutations for a given number of contact residues and mutation rate), and the Valid parameter (1 if the experimental structure available in the PDB; 0 otherwise). The values of several of these variables may be modified through a sliding window. Each line represents a TM binary complex between Protein A and Protein B. The following eight columns correspond to the same variables for Protein B. The value of the last column is 0 if Valid A = Valid B = 0; 1 if Valid A or Valid B are 1; and 2 if Valid A = Valid B = 2, that is, when each of the two TM proteins possess an experimental structure. Two options are available in the top of the webpage: “Show X entries,” that is, the number of entries that can be shown in the page (up to 100), and a Search engine for searching any character (motif, digit, rate, protein name, etc.) in the whole database. The bottom of the webpage lists the number of entries resulting from a given choice of parameters (left), and the corresponding number of pages (left). This last option allows to pass from one page of the results to another.

#### Table S1

Membrane protein complexes used as templates in PPIMem

| A | B | C | D | E | F | H | I | J | K | L | M | N | O | Q | R |
|---|---|---|---|---|---|---|---|---|---|---|---|---|---|---|---|
| **P20963** | 2HAC | A | C.{2}LD.{2}L.{2}YG.{2}LT.LF | 10 | CYLLDGILFIYGVILTALF | 32 | 19 | **P20963** | 2HAC | B | C.{2}LD.{2}L.{2}YG.{2}LT.LF | 10 | CYLLDGILFIYGVILTALF | 32 | 19 |
| **P21709** | 2K1L | A | AV.{2}GL.{2}GA.{2}L | 8 | AVIFGLLLGAALL | 550 | 14 | **P21709** | 2K1L | B | AV.{2}GL.{2}GA.{2}L | 8 | AVIFGLLLGAALL | 550 | 14 |
| **P29317** | 2K9Y | A | IG.{2}A.{3}V.{2}L.{7}F | 6 | IGGVAVGVVLLLVLAGVGFF | 538 | 20 | **P29317** | 2K9Y | B | IG.{2}A.{3}V.{2}L.{7}F | 6 | IGGVAVGVVLLLVLAGVGFF | 538 | 20 |
| **Q15303** | 2L2T | A | PL.{2}AG.{3}G.{2}IL.{2}V | 8 | PLIAAGVIGGLFILVIV | 651 | 17 | **Q15303** | 2L2T | B | PL.{2}AG.{3}G.{2}IL.{2}V | 8 | PLIAAGVIGGLFILVIV | 651 | 17 |
| **O43914** | 2L34 | A | LA.IV.{2}D.{2}LT.{2}I.{3}V | 10 | LAGIVMGDLVLTVLIALAV | 43 | 19 | **O43914** | 2L34 | B | LA.IV.{2}D.{2}LT.{2}I.{3}V | 9 | LAGIVMGDLVLTVLIALAV | 43 | 19 |
| **P09619** | 2L6W | A | VV.{2}A.{2}A.{2}VL.{2}I.{3}I | 8 | VVISAILALVVLTIISLII | 533 | 19 | **P09619** | 2L6W | B | VV.{2}A.{2}A.{2}VL.{2}I.{3}I | 10 | VVISAILALVVLTIISLII | 533 | 19 |
| **P21860** | 2L9U | A | I.{2}LV.IF.{2}L.{3}F | 7 | IAGLVVIFMMLGGTF | 649 | 15 | **P21860** | 2L9U | B | I.{2}LV.IF.{2}L.{3}F | 7 | IAGLVVIFMMLGGTF | 649 | 15 |
| **P05067** | 2LOH | A | I.{2}LM.{2}G.{3}A.{2}IV.{2}L | 8 | IIGLMVGGVVIATVIVITL | 702 | 19 | **P05067** | 2LOH | B | I.{2}LM.{2}G.{3}A.{2}IV.{2}L | 8 | IIGLMVGGVVIATVIVITL | 702 | 19 |
| **P22607** | 2LZL | A | L.YGV.FF.{2}I.{6}TL | 9 | LSYGVGFFLFILVVAAVTL | 377 | 19 | **P22607** | 2LZL | B | L.YGV.FF.{2}I.{6}TL | 9 | LSYGVGFFLFILVVAAVTL | 377 | 19 |
| **P00533** | 2M0B | A | M.{3}L.{2}LL.{2}AL.{2}GL | 8 | MVGALLLLLVVALGIGL | 650 | 17 | **P00533** | 2M0B | B | M.{3}L.{2}LL.{2}AL.{2}GL | 8 | MVGALLLLLVVALGIGL | 650 | 17 |

| | | | | | | | | | | | | | | | |
|---|---|---|---|---|---|---|---|---|---|---|---|---|---|---|---|
| **P35968** | 2M59 | A | EI.IL.{9}FW.LL.{2}I | 9 | EIIILVGTAVIAMFFWLLLVII | 764 | 22 | **P35968** | 2M59 | B | EI.IL.{9}FW.LL.{2}I | 9 | EIIILVGTAVIAMFFWLLLVII | 764 | 22 |
| **O15455** | 2MK9 | A | S.{2}LI.{2}FI.{2}L | 6 | SILLIFIFIVLL | 711 | 12 | **O15455** | 2MK9 | B | S.{2}LI.{2}FI.{2}L | 6 | SILLIFIFIVLL | 711 | 12 |
| **O15455** | 2MKA | A | F.{3}N.{2}IL.{2}FI.{2}V.{3}H | 8 | FFMINTSILLIFIFIVLLIH | 705 | 20 | **O15455** | 2MKA | B | F.{3}N.{3}L.{2}FI.{6}HF | 7 | FFMINTSILLIFIFIVLLIHF | 705 | 21 |
| **O15455** | 2MKA | A | T.{2}LL.{2}IF.{2}LL.{2}F | 8 | TSILLIFIFIVLLIHF | 710 | 16 | **O15455** | 2MKA | C | F.{3}VL.{2}HF | 5 | FIFIVLLIHF | 716 | 10 |
| **P04626** | 2N2A | A | I.{3}V.{3}L.{2}VL.{6}L | 7 | ISAVVGILLVVVLGVVFGIL | 655 | 20 | **P04626** | 2N2A | B | I.{3}V.{3}L.{2}VL.{6}L | 7 | ISAVVGILLVVVLGVVFGIL | 655 | 20 |
| **P04629** | 2N90 | A | LA.FA.LFL | 7 | LAVFACLFL | 424 | 9 | **P04629** | 2N90 | B | LA.FA.LFL | 8 | LAVFACLFL | 424 | 15 |
| **Q1XA76** | 2QTS | A | G.M.{3}I.{3}I.{3}L.{2}F | 6 | GQMGLFIGASILTVLELF | 436 | 18 | **Q1XA76** | 2QTS | B | K.{3}W.{2}CF.{5}L.{379}Q | 6 | KRVVWALCFMGSLALLALVCTNRIQYYFLYPHVTKLDEVAATRLTFPAVTFCNLNEFRFSRVTKNDLYHAGELLALLNNRYEIPDTQTADEKQLEILQDKANFRNFKPKPFNMLEFYDRAGHDIREMLLSCFFRGEQCSPEDFKVVFTRYGKCYTFNAGQDGKPRLITMKGGTGNGLEIMLDIQQDEYLPVWGETDETSFEAGIKVQIHSQDEPPLIDQLGFGVAPGFQTFVSCQEQRLIYLPPPWGDCKATTGDSEFYDTYSITACRIDCETRYLVENCNCRMVHMPGDAPYCTPEQYKECADPALDFLVEKDNEYCVCEMPCNVTRYGKELSMVKIPSKASAKYLAKKYNKSEQYIGENILVLDIFFEALNYETIEQKKAYEVAGLLGDIGGQ | 43 | 395 |
| **P29033** | 2ZW3 | A | W.{2}V.{3}F.{2}M.{3}V.{3}EV.{143}E.{2}VF.{2}F | 11 | WLTVLFIFRIMILVVAAKEVWGDEQADFVCNTLQPGCKNVCYDHYFPISHIRLWALQLIFVSTPALLVAMHVAYRRHEKKRKFIKGEIKSEFKDIEEIKTQKVRIEGSLWWTYTSSIFFRVIFEAAFMYVFYVMYDGFSMQRLVKCNAWPCPNTVDCFVSRPTEKTVFTVF | 24 | 171 | **P29033** | 2ZW3 | B | IR.{3}L.{2}IF.{2}T.{3}L | 7 | IRLWALQLIFVSTPALL | 74 | 17 |
| **P55064** | 3D9S | B | K.{6}L.{6}FF.{2}GS.{11}LQ.{2}LA.{2}LA.{2}T.{2}QAL.PV.{40}I.{58}L | 20 | KAVFAEFLATLIFVFFGLGSALKWPSALPTILQIALAFGLAIGTLAQALGPVSGGHINPAITLALLVGNQISLLRAFFYVAAQLVGAIAGAGILYGVAPLNARGNLAVNALNNNTTQGQAMVVELILTFQLALCIFASTDSRRTSPVGSPAL | 12 | 152 | **P55064** | 3D9S | D | L.{86}Q.{3}V.{2}IL.{2}Q.{3}C.{2}AS.{14}S.{2}LS.TL.{2}LV.{44}L.{3}L | 18 | LQIALAFGLAIGTLAQALGPVSGGHINPAITLALLVGNQISLLRAFFYVAAQLVGAIAGAGILYGVAPLNARGNLAVNALNNNTTQGQAMVVELILTFQLALCIFASTDSRRTSPVGSPALSIGLSVTLGHLVGIYFTGCSMNPARSFGPAVVMNRFSPAHWVFWVGPIVGAVLAAILYFYL | 43 | 182 |

| | | | | | | | | | | | | | | | |
|---|---|---|---|---|---|---|---|---|---|---|---|---|---|---|---|
| **O35433** | 3J5P | A | R.{2}FV.{2}VF.{2}G.{2}TA.VT.{58}K.V.I.L.{5}I.{2}YIL.{2}NM.IAL | 23 | RFMFVYLVFLFGFSTAVVTLIEDGKNNSLPMESTPHKCRGSACKPGNSYNSLYSTCLELFKFTIGMGDLEFTENYDFKAVFIILLLAYVILTYILLLNMLIAL | 579 | 103 | **O35433** | 3J5P | B | T.{2}AY.{85}A.{2}VF.L.{3}WT.{2}L.{18}MI.{3}L.{97}L.{2}LI.{2}M | 17 | TAAAYYRPVEGLPPYKLKNTVGDYFRVTGEILSVSGGVYFFFRGIQYFLQRRPSLKSLFVDSYSEILFFVQSLFMLVSVVLYFSQRKEYVASMVFSLAMGWTNMLYYTRGFQQMGIYAVMIEKMILRDLCRFMFVYLVFLFGFSTAVVTLIEDGKNNSLPMESTPHKCRGSACKPGNSYNSLYSTCLELFKFTIGMGDLEFTENYDFKAVFIILLLAYVILTYILLLNMLIALM | 449 | 234 |
| **P48542** | 3SYQ | A | L.{3}V.{3}T.{2}W.{81}V.{2}MF | 7 | LLIFVMVYTVTWLFFGMIWWLIAYIRGDMDHIEDPSWTPCVTNLNGFVSAFLFSIETETTIGYGYRVITDKCPEGIILLLIQSVLGSIVNAFMVGCMF | 95 | 98 | **P48542** | 3SYQ | B | LL.{3}VL.SI.{2}AF.{2}G | 9 | LLIQSVLGSIVNAFMVG | 173 | 17 |
| **P57789** | 4BW5 | C | V.{2}IF.{2}V.{2}YL.{2}GG.VFR.{87}GGKIFCI.Y.IFGIPL.{2}FL.{2}G.{103}L.{9}L.{2}IG | 35 | VVAIFVVVVVYLVTGGLVFRALEQPFESSQKNTIALEKAEFLRDHVCVSPQELETLIQHALDADNAGVSPIGNSSNNSSHWDLGSAFFFAGTVITTIGYGNIAPSTEGGKIFCILYAIFGIPLFGFLLAGIGDQLGTIFGKSIARVEKVFRKKQVSQTKIRVISTILFILAGCIVFVTIPAVIFKYIEGWTALESIYFVVVTLTTVGFGDFVAGGNAGINYREWYKPLVWFWILVGLAYFAAVLSMIG | 72 | 252 | P57789 | 4BW5 | D | V.{2}IF.{2}V.{2}YL.{2}GG.VFR.{87}GGKIFCI.Y.IFGIPL.{2}FL.{2}G.{103}L.{9}L.{2}IG | 39 | VVAIFVVVVVYLVTGGLVFRALEQPFESSQKNTIALEKAEFLRDHVCVSPQELETLIQHALDADNAGVSPIGNSSNNSSHWDLGSAFFFAGTVITTIGYGNIAPSTEGGKIFCILYAIFGIPLFGFLLAGIGDQLGTIFGKSIARVEKVFRKKQVSQTKIRVISTILFILAGCIVFVTIPAVIFKYIEGWTALESIYFVVVTLTTVGFGDFVAGGNAGINYREWYKPLVWFWILVGLAYFAAVLSMIG | 72 | 250 |
| **P28472** | 4COF | A | Q.{6}LI.IL.{5}WI.{6}A.{2}AL.{2}T | 15 | QTYMPSILITILSWVSFWINYDASAARVALGIT | 249 | 206 | **P28472** | 4COF | B | S.{3}V.{3}I.{3}L.{6}T.{19}M.{2}F.{3}F.{2}LL.{5}N | 11 | SAARVALGITTVLTMTTINTHLRETLPKIPYVKAIDMYLMGCFVFVFLALLEYAFVN | 272 | 57 |
| **Q5QF96** | 4EZC | A | L.{45}T.{3}F.SA.{2}SV.{18}M.{2}SAT.{140}V | 12 | LSQDRSAITAGLQGYNATLVGILMAIYSDKGNYFWWLLFPVSAMSMTCPVFSSALNSVLSKWDLPVFTLPFNMALSMYLSATGHYNPFFPSTLITPVTSVPNVTWPDLSALQLLKSLPVGVGQIYGCDNPWTGGIFLGAILLSSPLMCLHAAIGSLLGIIAGLSLSAPFEDIYAGLWGFNSSLACIAIGGTFMALTWQTHLLALACALFTAYLGASMSHVMAV | 105 | 223 | **Q5QF96** | 4EZC | B | LSL.{43}F.{2}YL.AS.{2}HV.{2}V.{15}L.{2}L | 13 | LSLSAPFEDIYAGLWGFNSSLACIAIGGTFMALTWQTHLLALACALFTAYLGASMSHVMAVVGLPSGTWPFCLATLLFLL | 267 | 80 |
| **Q99835** | 4JKV | A | Y.{2}H.{19}F.LL.{6}VL.VA.{2}A.{24}GF.{2}AP.{6}V.{2}YF.{2}R | 18 | YAWHTSFKALGTTYQPLSGKTSYFHLLTWSLPFVLTVAILAVAQVDGDSVSGICFVGYKNYRYRAGFVLAPIGLVLIVGGYFLIR | 337 | 85 | **Q99835** | 4JKV | B | V.{3}Y.{24}LL.{6}VL.VA.{2}A.{24}GF.{2}A.{11}F | 13 | VVLTYAWHTSFKALGTTYQPLSGKTSYFHLLTWSLPFVLTVAILAVAQVDGDSVSGICFVGYKNYRYRAGFVLAPIGLVLIVGGYF | 333 | 86 |
| P41181 | 4NEF | A | R.{2}FA.{2}L.{3}L.{2}FF.{2}GS.{11}LQ.{2}MA.{2}LG.{2}T.VQALG.{42}LL.{57}L | 24 | RAVFAEFLATLLFVFFGLGSALNWPQALPSVLQIAMAFGLGIGTLVQALGHISGAHINPAVTVACLVGCHVSVLRAAFYVAAQLLGAVAGAALLHEITPADIRGDLAVNALSNSTTAGQAVTVELFLTLQLVLCIFASTDERRGENPGTPAL | 11 | 152 | **P41181** | 4NEF | C | VL.{86}Q.{2}TV.{2}FL.LQ.{2}LC.{2}AS.{14}S.{2}FS.AL.{2}LL.{43}LY | 22 | VLQIAMAFGLGIGTLVQALGHISGAHINPAVTVACLVGCHVSVLRAAFYVAAQLLGAVAGAALLHEITPADIRGDLAVNALSNSTTAGQAVTVELFLTLQLVLCIFASTDERRGENPGTPALSIGFSVALGHLLGIHYTGCSMNPARSLAPAVVTGKFDDHWVFWIGPLVGAILGSLLY | 41 | 179 |
| **Q05085** | 4OH3 | A | I.{2}A.{3}T.{3}I.{5}I.{97}W.{6}F.{2}V.{2}LS.{2}L | 12 | IFAAIQATGVSILTLSTIIPGLRPPRCNPTTSSHCEQASGIQLTVLYLALYLTALGTGGVKASVSGFGSDQFDETEPKERSKMTYFFNRFFFCINVGSLLAVTVLVYVQDDVGRKWGYGICAFAIVLALSVFL | 104 | 133 | **Q05085** | 4OH3 | B | I.{2}A.{3}T.{3}I.{5}I.{97}W.{6}F.{2}V.{2}LS.{2}L | 13 | IFAAIQATGVSILTLSTIIPGLRPPRCNPTTSSHCEQASGIQLTVLYLALYLTALGTGGVKASVSGFGSDQFDETEPKERSKMTYFFNRFFFCINVGSLLAVTVLVYVQDDVGRKWGYGICAFAIVLALSVFL | 104 | 134 |
| **O95069** | 4TWK | A | W.{2}V.{2}IF.{2}V.{2}YLI.GA.VF.{88}GGKI | 39 | WKTVSTIFLVVVLYLIIGATVFKALEQPHEISQRTTIVIQKQTFISQHSCVNSTELDELIQQIVAAINAGIIPLGNTSNQISHWDLGSSFFFAGTVITTIGFGNISPRTEGGKIFCIIYALLGIPLFGFLLAGVGDQLGTIFGKGIAKVEDTFIKWNVSQTKIRIISTIIFILFGCV | 59 | 254 | **O95069** | 4TWK | B | W.{2}V.{2}IF.{2}V.{2}YLI.GA.VF.{88}GGKIFC.{2}YALLGIPL.{2}FL.{2}G.{2}DQ.{94}W.{ | 44 | WKTVSTIFLVVVLYLIIGATVFKALEQPHEISQRTTIVIQKQTFISQHSCVNSTELDELIQQIVAAINAGIIPLGNTSNQISHWDLGSSFFFAGTVITTIGFGNISPRTEGGKIFCIIYALLGIPLFGFLLAGVGDQLGTIFGKGIAKVEDTFIKWNVSQTKI | 59 | 254 |

| | | | | | | | | | | | | | | | |
|---|---|---|---|---|---|---|---|---|---|---|---|---|---|---|---|
| | | | F C.{2}YALLGIPL.{2}FL.{2}G.{2}D Q.{94}W.{13}L.{2}IG.{2}LR | | LFVALPAIIFKHIEGWSALDAIYFVVITLTTIGFGDYVAGGSDIEYLDFYKPVVVWFWILVGLAYFAAVLSMIGDWLR | | | | | | 13}L.{2}IG.{2}LR | | RIISTIIFILFGCVLFVALPAIIFKHIEGWSALDAIYFVVITLTTIGFGDYVAGGSDIEYLDFYKPVVVWFWILVGLAYFAAVLSMIGDWLR | | |
| **Q9NYG8** | 4WFE | A | L.{2}L.{3}V.{2}YL.{2}GA.VF.{65}L.{2}AF.FSG.{2}I.{12}D.GR.FCI.Y.LVGIPL.{2}IL.{2}GV.{2}R.{58}L.{3}Y.{2}I.{3}T.{45}L | 39 | LLALLALVLLYLVSGALVFRALEQPHEQQAQRELGEVREKFLRAHPCVSDQELGLLIKEVADALGGGADPETNSTSNSSHSAWDLGSAFFFSGTIITTIGYGNVALRTDAGRLFCIFYALVGIPLFGILLAGVGDRLGSSLRHGIGHIEAIFLKWHVPPELVRVLSAMLFLLIGCLLFVLTPTFVFCYMEDWSKLEAIYFVIVTLTTVGFGDYVAGADPRQDSPAYQPLVWFWILLGLAYFASVLTTIGNWL | 6 | 252 | **Q9NYG8** | 4WFE | B | L.{2}LL.LVL.YL.{2}GA.VFR.L.{62}L.{2}AF.FSG.{2}I.{14}GR.FCI.Y.{2}VGI.L.{2}IL.{2}G.{3}R.{58}L.{3}Y.{2}I.{3}T.{24}W.{9}F.{3}L.{2}IG.{2}LR | 46 | LLALLALVLLYLVSGALVFRALEQPHEQQAQRELGEVREKFLRAHPCVSDQELGLLIKEVADALGGGADPETNSTSNSSHSAWDLGSAFFFSGTIITTIGYGNVALRTDAGRLFCIFYALVGIPLFGILLAGVGDRLGSSLRHGIGHIEAIFLKWHVPPELVRVLSAMLFLLIGCLLFVLTPTFVFCYMEDWSKLEAIYFVIVTLTTVGFGDYVAGADPRQDSPAYQPLVWFWILLGLAYFASVLTTIGNWLR | 6 | 253 |
| **O43914** | 4WO1 | B | P.{2}L.{3}V.{2}DL.{6}A.{3}Y | 7 | PGVLAGIVMGDLVLTVLIALAVY | 40 | 23 | **O43914** | 4WO1 | C | L.{2}I.{3}D.{3}T.{2}I.{3}V.{2}L | 7 | LAGIVMGDLVLTVLIALAVYFL | 43 | 22 |
| **P23415** | 4X5T | A | Y.{2}I.{7}L.{2}IL.{16}GL.{2}T.{2}LT.{2}T.{2}SG.R | 14 | YYLIQMYIPSLLIVILSWISFWINMDAAPARVGLGITTVLTMTTQSSGSR | 250 | 50 | **P23415** | 4X5T | B | I.{3}L.{2}T.{3}S.{2}R.{19}L.{3}F.{2}L.{3}A | 9 | ITTVLTMTTQSSGSRASLPKVSYVKAIDIWMAVCLLFVFSALLEYA | 285 | 46 |
| **P49768** | 5A63 | B | DWN.{2}IA.{2}V.{2}LI.{24}F.{3}F.{2}A.{2}YLV.{2}FM | 16 | DWNTTIACFVAILIGLCLTLLLLAIFKKALPALPISITFGLVFYFATDYLVQPFM | 403 | 55 | **Q96BI3** | 5A63 | C | VI.{2}V.{3}F.WLV.LL.{2}SV.WF.{17}L.{6}SV.{2}Q.{44}IS.VFS.IN.{22}TS.{2}LT.{32}H.{3}S.{2}TF.N | 33 | VIILVAGAFFWLVSLLLASVVWFILVHVTDRSDARLQYGLLIFGAAVSVLLQEVFRFAYYKLLKKADEGLASLSEDGRSPISIRQMAYVSGLSFGIISGVFSVINILADALGPGVVGIHGDSPYYFLTSAFLTAAIILLHTFWGVVFFDACERRRYWALGLVVGSHLLTSGLTFLN | 32 | 176 |
| **Q92542** | 5A63 | A | E.{2}TL.VGF.IL.{2}SL.{2}TY | 12 | ELITLTVGFGILIFSLIVTY | 669 | 20 | **Q96BI3** | 5A63 | C | AV.{2}G.{2}FV.{2}GP.{2}AL.{2}I.{112}NI.{19}FL.{3}F | 15 | AVFFGCTFVAFGPAFALFLITVAGDPLRVIILVAGAFFWLVSLLLASVVWFILVHVTDRSDARLQYGLLIFGAAVSVLLQEVFRFAYYKLLKKADEGLASLSEDGRSPISIRQMAYVSGLSFGIISGVFSVINILADALGPGVVGIHGDSPYYFLTSAF | 4 | 159 |
| **P49768** | 5A63 | B | F.{3}G.{2}F.{8}YITV.L.{2}WN.{2}VV.{2}I.{9}L | 14 | FIYLGEVFKTYNVAVDYITVALLIWNFGVVGMISIHWKGPLRL | 179 | 43 | **Q9NZ42** | 5A63 | D | W.{3}L.{2}W.{3}FQ | 5 | WVIVLTSWITIFQ | 67 | 13 |
| **P41948** | 5AEX | A | G.{3}A.{2}WI.{2}P.{18}LW.{2}MM.SA.{2}I.{3}F.{45}V.{3}M.{18}LF.{2}M | 18 | GVASAGVWIMVPGIGLLYSGLSRKKHALSLLWASMMASAVCIFQWFFWGYSLAFSHNTRGNGFIGTLEFFGFRNVLGAPSSVSSLPDILFAVYQGMFAAVTGALMLGGACERARLFPMM | 36 | 119 | **P41948** | 5AEX | B | VT.{2}V.{3}V.{2}WF.{2}MF.{17}M.{2}NL.{2}A.{6}M.{12}T.{2}L.{23}V | 16 | VTSVVLGTVFLWFGWMFFNGGSAGNATIRAWYSIMSTNLAAACGGLTWMVIDYFRCGRKWTTVGLCSGIIAGLVGITPAAGFVPIWSAV | 228 | 89 |
| **O75311** | 5CFB | A | I.{3}L.{2}T.{6}R.{12}D.{2}MA.{6}F.{2}LL | 10 | ITTVLTMTTQSSGSRASLPKVSYVKAIDIWMAVCLLFVFSALL | 290 | 43 | **O75311** | 5CFB | B | GYY.IQ.{6}L.{2}IL.{2}V.{13}A.{3}T.{3}T.{3}Q | 13 | GYYLIQMYIPSLLIVILSWVSFWINMDAAPARVALGITTVLTMTTQ | 254 | 46 |
| **Q5N8** | 5CTG | A | M.{3}G.{3}L.{2}C.{3}LV.HA.{2}FF.{8}F | 11 | MKIIGLLVLVVCGFALVSHASVFFFDQPLRQQF | 102 | 33 | **Q5N8** | 5CTG | B | L.{4}F.{2}YF.{2}GL.{2}I.{3}M.{3}L.{2}Y | 10 | LLLRDFFIYFPNGLGLILGAMQLALYAY | 186 | 28 |

| | | | | | | | | | | | | | | | |
|---|---|---|---|---|---|---|---|---|---|---|---|---|---|---|---|
| **J1** | | | | | | | | **J1** | | | | | | | |
| **O35433** | 5IRX | A | R.{3}V.{2}VF.FG.{2}TA.VTL.{59}V.{2}IL.L.{2}VI.{2}YILL.NM.{2}AL | 25 | RFMFVYLVFLFGFSTAVVTLIEDGKNNSLPMESTPHKCRGSACKPGNSYNSLYSTCLELFKFTIGMGDLEFTENYDFKAVFIILLLAYVILTYILLLNMLIAL | 579 | 103 | **O35433** | 5IRX | C | AY.{82}E.VA.{2}VF.LA.{2}WT.{124}L.{3}I.{2}M | 14 | AYYRPVEGLPPYKLKNTVGDYFRVTGEILSVSGGVYFFFRGIQYFLQRRPSLKSLFVDSYSEILFFVQSLFMLVSVVLYFSQRKEYVASMVFSLAMGWTNMLYYTRGFQQMGIYAVMIEKMILRDLCRFMFVYLVFLFGFSTAVVTLIEDGKNNSLPMESTPHKCRGSACKPGNSYNSLYSTCLELFKFTIGMGDLEFTENYDFKAVFIILLLAYVILTYILLLNMLIALM | 452 | 231 |
| **P31644** | 5O8F | A | V.{3}T.{3}V.{3}L.{6}I.{22}Y.{3}F.{2}L.{3}A | 9 | VPARTVFGVTTVLTMTTLSISARNSLPKVAYATAMDWFIAVCYAFVFSALIEFA | 286 | 54 | **P31644** | 5O8F | B | IQ.{6}M.{2}IL.{2}V.{3}L.{6}A.{3}F.{2}T.{3}T.{130}R | 12 | IQTYLPCIMTVILSQVSFWLNRESVPARTVFGVTTVLTMTTLSISARNSLPKVAYATAMDWFIAVCYAFVFSALIEFATVNYFTKRGWAWDGKKALEAAKIKKKREVILNKSTNAFTTGKMSHPPNIPKEQTPAGTSNTTSVSVKPSEEKTSESKKTYNSISKIDKMSR | 262 | 169 |
| **P21730** | 5O9H | A | L.{6}LY.{23}R | 6 | LLTIPSFLYRVVREEYFPPKVLCGVDYSHDKRR | 166 | 47 | **P21730** | 5O9H | B | L.{6}LY.{23}R | 6 | LLTIPSFLYRVVREEYFPPKVLCGVDYSHDKRR | 166 | 33 |
| **P10912** | 5OEK | A | FG.FG.{2}VM | 6 | FGIFGLTVM | 273 | 9 | **P10912** | 5OEK | B | FG.FG.{2}VM | 6 | FGIFGLTVM | 273 | 9 |
| **P10912** | 5OHD | A | W.{3}II.{2}I.{3}T.{3}F | 6 | WLLIIIFGIFGLTVMLF | 267 | 17 | **P10912** | 5OHD | B | W.{3}II.{2}I.{3}T.{3}F | 6 | WLLIIIFGIFGLTVMLF | 267 | 17 |
| **Q13563** | 5T4D | A | K.{3}GF.{2}M.{2}II.{2}AY.QL.YL.{38}L.P.{2}F.{3}VF.{3}FI.{2}NM.LAI.N | 25 | KDLFGFAIMFFIIFLAYAQLAYLVFGTQVDDFSTFQECIFTQFRIILGDINFAEIEEANRVLGPIYFTTFVFFMFFILLNMFLAIIN | 595 | 87 | **Q13563** | 5T4D | B | N.{2}A.{2}VF.{2}W.{3}F.{2}I.{27}F.{67}L.{3}L.{2}IN | 12 | NIAAVTVFFVWIKLFKFINFNRTMSQLSTTMSRCAKDLFGFAIMFFIIFLAYAQLAYLVFGTQVDDFSTFQECIFTQFRIILGDINFAEIEEANRVLGPIYFTTFVFFMFFILLNMFLAIIN | 560 | 122 |
| **O75311** | 5TIN | A | A.{3}V.{3}I.{3}L.{6}S.{2}R.{19}L.{3}F.{2}L | 9 | APARVALGITTVLTMTTQSSGSRASLPKVSYVKAIDIWMAVCLLFVFSAL | 282 | 50 | **O75311** | 5TIN | B | Q.{6}LI.IL.{5}WI.{6}A.{2}AL.{2}T | 14 | QMYIPSLLIVILSWVSFWINMDAAPARVALGIT | 259 | 103 |
| **O60741** | 5U6O | A | Y.{2}FV.{3}T.{2}IQ | 6 | YAMFVGHATALIQ | 386 | 13 | **O60741** | 5U6O | B | R.{74}LW.{2}M.{2}MI.{2}A.{2}YAM.VGH.T.{3}Q | 15 | RIFNLIGMMLLLCHWDGCLQFLVPLLQDFPPDCWVSLNEMVNDSWGKQYSYALFKAMSHMLCIGYGAQAPVSMSDLWITMLSMIVGATCYAMFVGHATALIQ | 297 | 102 |
| **Q9GZU1** | 5WJ9 | A | F.{2}C.{2}VI.{2}GY.FC.WIV.{2}P.{39}LVW.F.Q.{3}Y.{2}I.{3}IYM.{2}S | 23 | FCCCVAVIYLGYCFCGWIVLGPYHVKFRSLSMVSECLFSLINGDDMFVTFAAMQAQQGRSSLVWLFSQLYLYSFISLFIYMVLS | 428 | 84 | **Q9GZU1** | 5WJ9 | B | T.{2}LI.{302}D.{2}SI.{2}GT.{2}L.{2}WV.VI.{2}L | 14 | TVQLILFGLSNQLAVTFREENTIAFRHLFLLGYSDGADDTFAAYTREQLYQAIFHAVDQYLALPDVSLGRYAYVRGGGDPWTNGSGLALCQRYYHRGHVDPANDTFDIDPMVVTDCIQVDPPERPPPPPSDDLTLLESSSSYKNLTLKFHKLVNVTIHFRLKTINLQSLINNEIPDCYTFSVLITFDNKAHSGRIPISLETQAHIQECKHPSVFQHGDNSFRLLFDVVVILTCSLSFLLCARSLLRGFLLQNEFVGFMWRQRGRVISLWERLEFVNGWYILLVTSDVLTI | 77 | 329 |

| | | | | | | | | | | | | | | | |
|---|---|---|---|---|---|---|---|---|---|---|---|---|---|---|---|
| | | | | | | | | | | | | | SGTIMKIGIEAKNLASYDVCSILLGTSTLLVWVGVIRYL | | |
| **A2A259** | 5Z1W | A | Y.{2}T.{317}N.{2}A.{2}LF.{2}W.{3}F | 8 | YGMTSSSAYYYTKVMSELFLHTPSDSGVSFQTISSMSDFWDFAQGPLLDSLYWTKWYNNQSLGRGSHSFIYYENLLLGAPRLRQLRVRNDSCVVHEDFREDILNCYDVYSPDKEDQLPFGPQNGTAWTYHSQNELGGSSHWGRLTSYSGGGYYLDLPGSRQASAEALQGLQEGLWLDRGTRVVFIDFSVYNANINLFCILRLVVEFPATGGTIPSWQIRTVKLIRYVNNWDFFIVGCEVVFCVFIFYYVVEEILEIHLHRLRYLSSVWNILDLVVILLSIVAVGFHIFRTLEVNRLMGKLLQQPDTYADFEFLAFWQTQYNNMNAVNLFFAWIKIF | 119 | 336 | **A2A259** | 5Z1W | C | M.{6}AY.QL.{2}LL.G.{32}NR.{3}P.{2}FV.{2}V | 14 | MFFIVFFAYAQLGYLLFGTQVENFSTFVKCIFTQFRIILGDFDYNAIDNANRILGPVYFVTYV | 483 | 63 |
| **Q9QUQ5** | 5Z96 | A | KF.FI.{2}L.{3}AF.NG.NQL.F.{52}F.{2}FV.A.MFG.{2}NV.{2}L.{4}NM | 25 | KFLFIYCLVLLAFANGLNQLYFYYEETKGLSCKGIRCEKQNNAFSTLFETLQSLFWSIFGLINLYVTNVKAQHEFTEFVGATMFGTYNVISLVVLLNM | 518 | 98 | **Q9QUQ5** | 5Z96 | B | LL.AS.{90}LV.EA.FAI.NI.{2}S.{2}L.{3}F.{19}I.{100}I | 18 | LLLASQHIDRSDLNRQGPPPTIVEWMILPWVLGFIWGEIKQMWDGGLQDYIHDWWNLMDFVMNSLYLATISLKIVAFVKYSALNPRESWDMWHPTLVAEALFAIANIFSSLRLISLFTANSHLGPLQISLGRMLLDILKFLFIYCLVLLAFANGLNQLYFYYEETKGLSCKGIRCEKQNNAFSTLFETLQSLFWSIFGLINLYVTNVKAQHEFTEFVGATMFGTYNVISLVVLLNMLI | 380 | 238 |
| **Q7TN37** | 6BCO | A | D.{2}F.{2}FF.{2}VW.VA.{2}VA.EG.{68}L.V.L.{4}LL.{2}NIL.{2}NL.I | 26 | DVFFFLFFLCVWLVAYGVATEGILRPQDRSLPSILRRVFYRPYLQIFGQIPQEEMDVALMIPGNCSMERGSWAHPEGPVAGSCVSQYANWLVVLLLIVFLLVANILLLNLLI | 925 | 113 | **Q7TN37** | 6BCO | B | L.{86}D.{2}RT.{2}CL.FM.{2}TL.{2}L.{2}F.{19}V.{2}F.{105}LI | 16 | LVDFQPTKPSVSELLLYFWAFTLLCEELRQGLGGGWGSLASGGRGPDRAPLRHRLHLYLSDTWNQCDLLALTCFLLGVGCRLTPGLFDLGRTVLCLDFMIFTLRLLHIFTVNKQLGPKIVIVSKMMKDVFFFLFFLCVWLVAYGVATEGILRPQDRSLPSILRRVFYRPYLQIFGQIPQEEMDVALMIPGNCSMERGSWAHPEGPVAGSCVSQYANWLVVLLLIVFLLVANILLLNLLI | 798 | 239 |
| **Q8TD43** | 6BQV | A | D.{2}F.{2}FF.{2}VW.VA.{2}VA.EG.{68}L.V.L.{4}LL.{2}NIL.{2}NL.I | 28 | DVFFFLFFLGVWLVAYGVATEGLLRPRDSDFPSILRRVFYRPYLQIFGQIPQEDMDVALMEHSNCSSEPGFWAHPPGAQAGTCVSQYANWLVVLLLVIFLLVANILLVNLLI | 929 | 114 | **Q8TD43** | 6BQV | B | S.{3}L.{87}L.RT.{2}CI.FM.{2}T.{2}LL.{2}F.{19}V.{2}FL.{104}LI.{2}FS | 20 | SRVLLVDFQPAPPGSLELLLYFWAFTLLCEELRQGLSGGGGSLASGGPGPGHASLSQRLRLYLADSWNQCDLVALTCFLLGVGCRLTPGLYHLGRTVLCIDFMVFTVRLLHIFTVNKQLGPKIVIVSKMMKDVFFFLFFLGVWLVAYGVATEGLLRPRDSDFPSILRRVFYRPYLQIFGQIPQEDMDVALMEHSNCSSEPGFWAHPPGAQAGTCVSQYANWLVVLLLVIFLLVANILLVNLLIAMFS | 798 | 247 |
| **P97438** | 6CQ9 | A | W.{2}V.{2}IF.{2}V.{2}YLI.GA.VF.{88}GGKIFC.{2}YALLGIPL.{2}FL.{2}G.{2}DQ.{94}W.{13}L.{2}IG.{2}LR | 36 | WKTVSTIFLVVVLYLIIGATVFKALEQPQEISQRTTIVIQKQTFIAQHACVNSTELDELIQQIVAAINAGIIPLGNSSNQVSHWDLGSSFFFAGTVITTIGFGNISPRTEGGKIFCIIYALLGIPLFGFLLAGVGDQLGTIFGKGIAKVEDTFIKWNVSQTKIRIISTIIFILFGCVLFVALPAVIFKHIEGWSALDAIYFVVITLTTIGFGDYVAGGSDIEYLDFYKPVVWFWILVGLAYFAAVLSMIGDWLR | 59 | 255 | **P97438** | 6CQ9 | B | W.{2}V.{2}IF.{2}V.{2}YLI.GA.VF.{88}GGKIFC.{2}YALLGIPL.{2}FL.{2}G.{2}DQ.{94}W.{13}L.{2}IG.{2}LR | 34 | WKTVSTIFLVVVLYLIIGATVFKALEQPQEISQRTTIVIQKQTFIAQHACVNSTELDELIQQIVAAINAGIIPLGNSSNQVSHWDLGSSFFFAGTVITTIGFGNISPRTEGGKIFCIIYALLGIPLFGFLLAGVGDQLGTIFGKGIAKVEDTFIKWNVSQTKIRIISTIIFILFGCVLFVALPAVIFKHIEGWSALDAIYFVVITLTTIGFGDYVAGGSDIEYLDFYKPVVWFWILVGLAYFAAVLSMIGDWLR | 59 | 253 |
| **Q13507** | 6CUD | A | KF.VL.{5}FAF.{2}G.FIL.S.{41}FI.{2}I.Y.{2}YG.{2}N.{7}LN | 21 | KFMVLFIMVFFAFMIGMFILYSYYLGAKVNAAFTTVEESFKTLFWSIFGLSEVTSVVLKYDHKFIENIGYVLYGIYNVTMVVVLLN | 567 | 86 | **Q13507** | 6CUD | B | A.{2}RFE.{126}I.{2}EGL.AI.{2}V.{2}F.{115}L | 13 | ASDRFEGITTLPNITVTDYPKQIFRVKTTQFTWTEMLIMVWVLGMMWSECKELWLEGPREYILQLWNVLDFGMLSIFIAAFTARFLAFLQATKAQQYVDSYVQESDLSEVTLPPEIQYFTYARDKWLPS | 392 | 263 |

| | | | | | | | | | | | | | | | |
|---|---|---|---|---|---|---|---|---|---|---|---|---|---|---|---|
| | | | | | | | | | | | | | DPQIISEGLYAIAVVLSFSRIAYILPANESFGPLQISLGRTVKDIFKFMVLFIMVFFAFMIGMFILYSYYLGAKVNAAFTTVEESFKTLFWSIFGLSEVTSVVLKYDHKFIENIGYVLYGIYNVTMVVVLLNML | | |
| **Q9UNQ0** | 6ETI | A | L.{2}VI.{2}IY.{14}G.{2}F.{115}F.{2}IF.GLLV | 14 | LGLVIGAIYFGLKNDSTGIQNRAGVLFFLTTNQCFSSVSAVELFVVEKKLFIHEYISGYYRVSSYFLGKLLSDLLPMRMLPSIIFTCIVYFMLGLKPKADAFFVMMFTLMMVAYSASSMALAIAAGQSVVSVATLLMTICFVFMMIFSGLLV | 405 | 152 | **Q9UNQ0** | 6ETI | B | L.{2}VI.{2}IY.{14}G.{2}F.{115}F.{2}IF.GLLV | 14 | LGLVIGAIYFGLKNDSTGIQNRAGVLFFLTTNQCFSSVSAVELFVVEKKLFIHEYISGYYRVSSYFLGKLLSDLLPMRMLPSIIFTCIVYFMLGLKPKADAFFVMMFTLMMVAYSASSMALAIAAGQSVVSVATLLMTICFVFMMIFSGLLV | 405 | 152 |

Column A (J): UniProt accession number of protein A (B)

B (K): PDB accession code of protein A (B)

C (L): Chain name of protein A (B)

D (M): Consensus binding motif of protein A (B)

E (N): Number of interface contact residues of protein A (B)

F (O): Matched motif sequence (motif instance) of protein A (B)

H (Q): Position of first residue of matched motif sequence of protein A (B)

I (R): Length of binding motif of protein A (B)

Rows highlighted in light blue represent homodimers in which each protomer presents a different interface.

Rows highlighted in light yellow represent heterodimers.

#### Table S2

Transmembrane protein oligomers presenting the L heptad pattern

| Pattern | TM MP oligomer | PDB code |
|---|---|---|
| L.{2}YG.{2}L | ζ-ζ | 2HAC |
| L.{2}GA.{2}L | Receptor tyrosine kinase EphA1 | 2K1L |
| LV.IF.{2}L | ErbB3 | 2L9U |
| L.{2}IF.{2}L | Toll-like receptor | 2MKA |
| L.{6}L | ErbB2 | 2N2A |
| LS.TL.{2}L | Human aquaporin 5 | 3D9S |
| L.{2}IG.{2}L | Two-pore domain potassium ion channel TREK1 | 4TWK |
| L.LVL.YL and L.{2}IG.{2}L | Human TRAAK K+ channel | 4WFE |
| L.{3}F.{2}L | α1 glycine receptor | 4X5T |
| L.VGF.IL | γ-secretase complex | 5A63 |
| L.{2}C.{3}L | SWEET transporter | 5CTG |
| L.NM.{2}AL | TRPV1 | 5IRX |
| L.{2}V.{3}L | β3-α5 GABAA receptor | 5O8F |
| L.{6}L | C5a anaphylatoxin chemotactic receptor 1 | 5O9H |
| L.{3}F.{2}L | Glycine receptor α-3 | 5TIN |
| L.FM.{2}TL | TRPM4 | 6BCO |
| L.{2}IG.{2}L | TREK-1 | 6CQ9 |

#### Table S3

PPIMem binding motif statistics for the membrane protein complexes used as templates in PPIMem

| A | B | C | D | E | F | G | H | I | J | K | L | M | N | O | P | Q | R |
|---|---|---|---|---|---|---|---|---|---|---|---|---|---|---|---|---|---|
| **2HAC (NMR)** | Low (Bio/Crystallographic) | - | P20963 | CD247 | P20963 | | A | B | 1 | C.{2}LD.{2}L.{2}YG.{2}LT.LF | C.{2}LD.{2}L.{2}YG.{2}LT.LF | 10 | 10 | 19 | 19 | MKWKALFTAAILQAQLPITEAQSFGLLDPKLCYLLDGILFIYGVILTALFLRVKFSRSADAPAYQQGQNQLYNELNLGRREEYDVLDKRRGRDPEMGGKPQRRKNPQEGLYNELQKDKMAEAYSEIGMKGERRRGKGHDGLYQGLSTATKDTYDALHMQALPPR | MKWKALFTAAILQAQLPITEAQSFGLLDPKLCYLLDGILFIYGVILTALFLRVKFSRSADAPAYQQGQNQLYNELNLGRREEYDVLDKRRGRDPEMGGKPQRRKNPQEGLYNELQKDKMAEAYSEIGMKGERRRGKGHDGLYQGLSTATKDTYDALHMQALPPR |
| **2JWA (NMR)** | Low (Bio/Crystallograp | - | P04626 | ERBB2 | P04626 | | A | B | 1 | T.{2}IS.{2}VG | T.{2}IS.{2}VG | 5 | 5 | 9 | 9 | MELAALCRWGLLLALLPPGAASTQVCTGTDMKLRLPASPETHLDMLRHLYQGCQVVQGNLELTYLPTNASLSFLQDIQEVQGYVLIAHNQVRQVPLQRLRIVRGTQLFEDNYALAVLDNGDPLNNTTPVTGASPGGLRELQLRSLTEILKGGVLIQRNPQLCYQDTILWKDIFHKNNQLALTLIDTNRSRACHPCSPMCKGSRCWGESSEDCQSLTRTVCAGGCARCKGPLPTDCCHEQCAAGCTGPKHSDCLACLHFNHSGICELHCPALVTYNTDTFESMPNPEGRYTFGASCVTACPYNYLSTDVGSCTLVCPLHNQEVTAEDGTQRCEKCSKPCARVCYGLGMEHLREVRAVTSANIQEFAGCKKIFGSLAF | MELAALCRWGLLLALLPPGAASTQVCTGTDMKLRLPASPETHLDMLRHLYQGCQVVQGNLELTYLPTNASLSFLQDIQEVQGYVLIAHNQVRQVPLQRLRIVRGTQLFEDNYALAVLDNGDPLNNTTPVTGASPGGLRELQLRSLTEILKGGVLIQRNPQLCYQDTILWKDIFHKNNQLALTLIDTNRSRACHPCSPMCKGSRCWGESSEDCQSLTRTVCAGGCARCKGPLPTDCCHEQCAAGCTGPKHSDCLACLHFNHSGICELHCPALVTYNTDTFESMPNPEGRYTFGASCVTACPYNYLSTDVGSCTLVCPLHNQEVTAEDGTQRCEKCSKPCARVCYGLGMEHLREVRAVTSANIQEFAGCKKIFGSLAF |

| | | | | | | | | | | | | | | | | | |
|---|---|---|---|---|---|---|---|---|---|---|---|---|---|---|---|---|---|
| | hic) | | | | | | | | | | | | | | | LPESFDGDPASNTAPLQPEQLQVFETLEEITGYLYISAWPDSLPDLSVFQNLQVIRGRILHNGAYSLTLQGLGISWLGLRSLRELGSGLALIHHNTHLCFVHTVPWDQLFRNPHQALLHTANRPEDECVGEGLACHQLCARGHCWGPGPTQCVNCSQFLRGQECVEECRVLQGLPREYVNARHCLPCHPECQPQNGSVTCFGPEADQCVACAHYKDPPFCVARCPSGVKPDLSYMPIWKFPDEEGACQPCPINCTHSCVDLDDKGCPAEQRASPLTSIISAVVGILLVVVLGVVFGILIKRRQQKIRKYTMRRLLQETELVEPLTPSGAMPNQAQMRILKETELRKVKVLGSGAFGTVYKGIWIPDGENVKIPVAIKVLRENTSPKANKEILDEAYVMAGVGSPYVSRLLGICLTSTVQLVTQLMPYGCLLDHVRENRGRLGSQDLLNWCMQIAKGMSYLEDVRLVHRDLAARNVLVKSPNHVKITDFGLARLLDIDETEYHADGGKVPIKWMALESILRRRFTHQSDVWSYGVTVWELMTFGAKPYDGIPAREIPDLLEKGERLPQPPICTIDVYMIMVKCWMIDSECRPRFRELVSEFSRMARDPQRFVVIQNEDLGPASPLDSTFYRSLLEDDDMGDLVDAEEYLVPQQGFFCPDPAPGAGGMVHHRHRSSSTRSGGGDLTLGLEPSEEEAPRSPLAPSEGAGSDVFDGDLGMGAAKGLQSLPTHDPSPLQRYSEDPTVPLPSETDGYVAPLTCSPQPEYVNQPDVRPQPPSPREGPLPAARPAGATLERPKTLSPGKNGVVKDVFAFGGAVENPEYLTPQGGAAPQPHPPPAFSPAFDNLYYWDQDPPERGAPPSTFKGTPTAENPEYLGLDVPV | LPESFDGDPASNTAPLQPEQLQVFETLEEITGYLYISAWPDSLPDLSVFQNLQVIRGRILHNGAYSLTLQGLGISWLGLRSLRELGSGLALIHHNTHLCFVHTVPWDQLFRNPHQALLHTANRPEDECVGEGLACHQLCARGHCWGPGPTQCVNCSQFLRGQECVEECRVLQGLPREYVNARHCLPCHPECQPQNGSVTCFGPEADQCVACAHYKDPPFCVARCPSGVKPDLSYMPIWKFPDEEGACQPCPINCTHSCVDLDDKGCPAEQRASPLTSIISAVVGILLVVVLGVVFGILIKRRQQKIRKYTMRRLLQETELVEPLTPSGAMPNQAQMRILKETELRKVKVLGSGAFGTVYKGIWIPDGENVKIPVAIKVLRENTSPKANKEILDEAYVMAGVGSPYVSRLLGICLTSTVQLVTQLMPYGCLLDHVRENRGRLGSQDLLNWCMQIAKGMSYLEDVRLVHRDLAARNVLVKSPNHVKITDFGLARLLDIDETEYHADGGKVPIKWMALESILRRRFTHQSDVWSYGVTVWELMTFGAKPYDGIPAREIPDLLEKGERLPQPPICTIDVYMIMVKCWMIDSECRPRFRELVSEFSRMARDPQRFVVIQNEDLGPASPLDSTFYRSLLEDDDMGDLVDAEEYLVPQQGFFCPDPAPGAGGMVHHRHRSSSTRSGGGDLTLGLEPSEEEAPRSPLAPSEGAGSDVFDGDLGMGAAKGLQSLPTHDPSPLQRYSEDPTVPLPSETDGYVAPLTCSPQPEYVNQPDVRPQPPSPREGPLPAARPAGATLERPKTLSPGKNGVVKDVFAFGGAVENPEYLTPQGGAAPQPHPPPAFSPAFDNLYYWDQDPPERGAPPSTFKGTPTAENPEYLGLDVPV |
| **2K** | Low | - | P21 | EPH | P21 | | A | B | 1 | GE.{2}AV.{2}GL. | GE.{2}AV.{2}GL. | 10 | 10 | 21 | 21 | MERRWPLGLGLVLLLCAPLPPGARAKEVTLMDTSKAQGELGWLLDPPKDGWSE | MERRWPLGLGLVLLLCAPLPPGARAKEVTLMDTSKAQGELGWLLDPPKDGWSE |

| | | | | | | | | | | | | | | | | | |
|---|---|---|---|---|---|---|---|---|---|---|---|---|---|---|---|---|---|
| **1K (NMR)** | (Bio>Crystallographic) | | 709 | A1 | 709 | | | | | {2}GA.{2}L.{3}L | {2}GA.{2}L.{3}L | | | | | QQQILNGTPLYMYQDCPMQGRRDTDHWLRSNWIYRGEEASRVHVELQFTVRDCKSFPGGAGPLGCKETFNLLYMESDQDVGIQLRRPLFQKVTTVAADQSFTIRDLVSGSVKLNVERCSLGRLTRRGLYLAFHNPGACVALVSVRVFYQRCPETLNGLAQFPDTLPGPAGLVEVAGTCLPHARASPRPSGAPRMHCSPDGEWLVPVGRCHCEPGYEEGGSGEACVACPSGSYRMDMDTPHCLTCPQQSTAESEGATICTCESGHYRAPGEGPQVACTGPPSAPRNLSFSASGTQLSLRWEPPADTGGRQDVRYSVRCSQCQGTAQDGGPCQPCGVGVHFSPGARGLTTPAVHVNGLEPYANYTFNVEAQNGVSGLGSSGHASTSVSISMGHAESLSGLSLRLVKKEPRQLELTWAGSRPRSPGANLTYELHVLNQDEERYQMVLEPRVLLTELQPDTTYIVRVRMLTPLGPGPFSPDHEFRTSPPVSRGLTGGEIVAVIFGLLLGAALLLGILVFRSRRAQRQRQQRQRDRATDVDREDKLWLKPYVDLQAYEDPAQGALDFTRELDPAWLMVDTVIGEGEFGEVYRGTLRLPSQDCKTVAIKTLKDTSPGGQWWNFLREATIMGQFSHPHILHLEGVVTKRKPIMIITEFMENGALDAFLREREDQLVPGQLVAMLQGIASGMNYLSNHNYVHRDLAARNILVNQNLCCKVSDFGLTRLLDDFDGTYETQGGKIPIRWTAPEAIAHRIFTTASDVWSFGIVMWEVLSFGDKPYGEMSNQEVMKSIEDGYRLPPPVDCPAPLYELMKNCWAYDRARRPHFQKLQAHLEQLLANPHSLRTIANFDPRMTLRLPSLSGSDGIPYRTVSEWLESIRMKRYILHFHSAGLDTMECVLELTAEDLTQMGITLPGHQKRILCSIQGFKD | QQQILNGTPLYMYQDCPMQGRRDTDHWLRSNWIYRGEEASRVHVELQFTVRDCKSFPGGAGPLGCKETFNLLYMESDQDVGIQLRRPLFQKVTTVAADQSFTIRDLVSGSVKLNVERCSLGRLTRRGLYLAFHNPGACVALVSVRVFYQRCPETLNGLAQFPDTLPGPAGLVEVAGTCLPHARASPRPSGAPRMHCSPDGEWLVPVGRCHCEPGYEEGGSGEACVACPSGSYRMDMDTPHCLTCPQQSTAESEGATICTCESGHYRAPGEGPQVACTGPPSAPRNLSFSASGTQLSLRWEPPADTGGRQDVRYSVRCSQCQGTAQDGGPCQPCGVGVHFSPGARGLTTPAVHVNGLEPYANYTFNVEAQNGVSGLGSSGHASTSVSISMGHAESLSGLSLRLVKKEPRQLELTWAGSRPRSPGANLTYELHVLNQDEERYQMVLEPRVLLTELQPDTTYIVRVRMLTPLGPGPFSPDHEFRTSPPVSRGLTGGEIVAVIFGLLLGAALLLGILVFRSRRAQRQRQQRQRDRATDVDREDKLWLKPYVDLQAYEDPAQGALDFTRELDPAWLMVDTVIGEGEFGEVYRGTLRLPSQDCKTVAIKTLKDTSPGGQWWNFLREATIMGQFSHPHILHLEGVVTKRKPIMIITEFMENGALDAFLREREDQLVPGQLVAMLQGIASGMNYLSNHNYVHRDLAARNILVNQNLCCKVSDFGLTRLLDDFDGTYETQGGKIPIRWTAPEAIAHRIFTTASDVWSFGIVMWEVLSFGDKPYGEMSNQEVMKSIEDGYRLPPPVDCPAPLYELMKNCWAYDRARRPHFQKLQAHLEQLLANPHSLRTIANFDPRMTLRLPSLSGSDGIPYRTVSEWLESIRMKRYILHFHSAGLDTMECVLELTAEDLTQMGITLPGHQKRILCSIQGFKD |

| | | | | | | | | | | | | | | | | | |
|---|---|---|---|---|---|---|---|---|---|---|---|---|---|---|---|---|---|
| **2K1L (NMR)** | Low (Bio>Crystallographic) | - | P21709 | EPHA1 | P21709 | | A | B | 1 | AV.{2}GL.{2}GA.{2}LL | AV.{2}GL.{2}GA.{2}LL | 8 | 8 | 14 | 14 | MERRWPLGLGLVLLLCAPLPPGARAKEVTLMDTSKAQGELGWLLDPPKDGWSEQQQILNGTPLYMYQDCPMQGRRDTDHWLRSNWIYRGEEASRVHVELQFTVRDCKSFPGGAGPLGCKETFNLLYMESDQDVGIQLRRPLFQKVTTVAADQSFTIRDLVSGSVKLNVERCSLGRLTRRGLYLAFHNPGACVALVSVRVFYQRCPETLNGLAQFPDTLPGPAGLVEVAGTCLPHARASPRPSGAPRMHCSPDGEWLVPVGRCHCEPGYEEGGSGEACVACPSGSYRMDMDTPHCLTCPQQSTAESEGATICTCESGHYRAPGEGPQVACTGPPSAPRNLSFSASGTQLSLRWEPPADTGGRQDVRYSVRCSQCQGTAQDGGPCQPCGVGVHFSPGARGLTTPAVHVNGLEPYANYTFNVEAQNGVSGLGSSGHASTSVSISMGHAESLSGLSLRLVKKEPRQLELTWAGSRPRSPGANLTYELHVLNQDEERYQMVLEPRVLLTELQPDTTYIVRVRMLTPLGPGPFSPDHEFRTSPPVSRGLTGGEIVAVIFGLLLGAALLLGILVFRSRRAQRQRQQRQRDRATDVDREDKLWLKPYVDLQAYEDPAQGALDFTRELDPAWLMVDTVIGEGEFGEVYRGTLRLPSQDCKTVAIKTLKDTSPGGQWWNFLREATIMGQFSHPHILHLEGVVTKRKPIMIITEFMENGALDAFLREREDQLVPGQLVAMLQGIASGMNYLSNHNYVHRDLAARNILVNQNLCCKVSDFGLTRLLDDFDGTYETQGGKIPIRWTAPEAIAHRIFTTASDVWSFGIVMWEVLSFGDKPYGEMSNQEVMKSIEDGYRLPPPVDCPAPLYELMKNCWAYDRARRPHFQKLQAHLEQLLANPHSLRTIANFDPRMTLRLPSLSGSDGIPYRTVSEWLESIRMKRYILHFHSAGLDT | MERRWPLGLGLVLLLCAPLPPGARAKEVTLMDTSKAQGELGWLLDPPKDGWSEQQQILNGTPLYMYQDCPMQGRRDTDHWLRSNWIYRGEEASRVHVELQFTVRDCKSFPGGAGPLGCKETFNLLYMESDQDVGIQLRRPLFQKVTTVAADQSFTIRDLVSGSVKLNVERCSLGRLTRRGLYLAFHNPGACVALVSVRVFYQRCPETLNGLAQFPDTLPGPAGLVEVAGTCLPHARASPRPSGAPRMHCSPDGEWLVPVGRCHCEPGYEEGGSGEACVACPSGSYRMDMDTPHCLTCPQQSTAESEGATICTCESGHYRAPGEGPQVACTGPPSAPRNLSFSASGTQLSLRWEPPADTGGRQDVRYSVRCSQCQGTAQDGGPCQPCGVGVHFSPGARGLTTPAVHVNGLEPYANYTFNVEAQNGVSGLGSSGHASTSVSISMGHAESLSGLSLRLVKKEPRQLELTWAGSRPRSPGANLTYELHVLNQDEERYQMVLEPRVLLTELQPDTTYIVRVRMLTPLGPGPFSPDHEFRTSPPVSRGLTGGEIVAVIFGLLLGAALLLGILVFRSRRAQRQRQQRQRDRATDVDREDKLWLKPYVDLQAYEDPAQGALDFTRELDPAWLMVDTVIGEGEFGEVYRGTLRLPSQDCKTVAIKTLKDTSPGGQWWNFLREATIMGQFSHPHILHLEGVVTKRKPIMIITEFMENGALDAFLREREDQLVPGQLVAMLQGIASGMNYLSNHNYVHRDLAARNILVNQNLCCKVSDFGLTRLLDDFDGTYETQGGKIPIRWTAPEAIAHRIFTTASDVWSFGIVMWEVLSFGDKPYGEMSNQEVMKSIEDGYRLPPPVDCPAPLYELMKNCWAYDRARRPHFQKLQAHLEQLLANPHSLRTIANFDPRMTLRLPSLSGSDGIPYRTVSEWLESIRMKRYILHFHSAGLDT |

| | | | | | | | | | | | | | | | | | |
|---|---|---|---|---|---|---|---|---|---|---|---|---|---|---|---|---|---|
| | | | | | | | | | | | | | | | | MECVLELTAEDLTQMGITLPGHQKRILCSIQGFKD | MECVLELTAEDLTQMGITLPGHQKRILCSIQGFKD |
| **2K9Y (NMR)** | Low (Bio>Crystallographic) | - | P29317 | EPHA2 | P29317 | | A | B | 1 | IG.{2}A.{3}V.{2}L.{7}F | IG.{2}A.{3}V.{2}L.{7}F | 6 | 6 | 20 | 20 | MELQAARACFALLWGCALAAAAAAQGKEVVLLDFAAAGGELGWLTHPYGKGWDLMQNIMNDMPIYMYSVCNVMSGDQDNWLRTNWVYRGEAERIFIELKFTVRDCNSFPGGASSCKETFNLYYAESDLDYGTNFQKRLFTKIDTIAPDEITVSSDFEARHVKLNVEERSVGPLTRKGFYLAFQDIGACVALLSVRVYYKKCPELLQGLAHFPETIAGSDAPSLATVAGTCVDHAVVPPGGEEPRMHCAVDGEWLVPIGQCLCQAGYEKVEDACQACSPGFFKFEASESPCLECPEHTLPSPEGATSCECEEGFFRAPQDPASMPCTRPPSAPHYLTAVGMGAKVELRWTPPQDSGGREDIVYSVTCEQCWPESGECGPCEASVRYSEPPHGLTRTSVTVSDLEPHMNYTFTVEARNGVSGLVTSRSFRTASVSINQTEPPKVRLEGRSTTSLSVSWSIPPPQQSRVWKYEVTYRKKGDSNSYNVRRTEGFSVTLDDLAPDTTYLVQVQALTQEGQGAGSKVHEFQTLSPEGSGNLAVIGGVAVGVVLLLVLAGVGFFIHRRRKNQRARQSPEDVYFSKSEQLKPLKTYVDPHTYEDPNQAVLKFTTEIHPSCVTRQKVIGAGEFGEVYKGMLKTSSGKKEVPVAIKTLKAGYTEKQRVDFLGEAGIMGQFSHHNIIRLEGVISKYKPMMIITEYMENGALDKFLREKDGEFSVLQLVGMLRGIAAGMKYLANMNYVHRDLAARNILVNSNLVCKVSDFGLSRVLEDDPEATYTTSGGKIPIRWTAPEAISYRKFTSASDVWSFGIVMWEVMTYGERPYWELSNHEVMKAINDGFRLPTPMDCPSAIYQLMMQCWQQERARRPKFADIVSILDKLIRAPDSLKTLADFDPRVSIRLPSTSGSEGVPFRTVSEW | MELQAARACFALLWGCALAAAAAAQGKEVVLLDFAAAGGELGWLTHPYGKGWDLMQNIMNDMPIYMYSVCNVMSGDQDNWLRTNWVYRGEAERIFIELKFTVRDCNSFPGGASSCKETFNLYYAESDLDYGTNFQKRLFTKIDTIAPDEITVSSDFEARHVKLNVEERSVGPLTRKGFYLAFQDIGACVALLSVRVYYKKCPELLQGLAHFPETIAGSDAPSLATVAGTCVDHAVVPPGGEEPRMHCAVDGEWLVPIGQCLCQAGYEKVEDACQACSPGFFKFEASESPCLECPEHTLPSPEGATSCECEEGFFRAPQDPASMPCTRPPSAPHYLTAVGMGAKVELRWTPPQDSGGREDIVYSVTCEQCWPESGECGPCEASVRYSEPPHGLTRTSVTVSDLEPHMNYTFTVEARNGVSGLVTSRSFRTASVSINQTEPPKVRLEGRSTTSLSVSWSIPPPQQSRVWKYEVTYRKKGDSNSYNVRRTEGFSVTLDDLAPDTTYLVQVQALTQEGQGAGSKVHEFQTLSPEGSGNLAVIGGVAVGVVLLLVLAGVGFFIHRRRKNQRARQSPEDVYFSKSEQLKPLKTYVDPHTYEDPNQAVLKFTTEIHPSCVTRQKVIGAGEFGEVYKGMLKTSSGKKEVPVAIKTLKAGYTEKQRVDFLGEAGIMGQFSHHNIIRLEGVISKYKPMMIITEYMENGALDKFLREKDGEFSVLQLVGMLRGIAAGMKYLANMNYVHRDLAARNILVNSNLVCKVSDFGLSRVLEDDPEATYTTSGGKIPIRWTAPEAISYRKFTSASDVWSFGIVMWEVMTYGERPYWELSNHEVMKAINDGFRLPTPMDCPSAIYQLMMQCWQQERARRPKFADIVSILDKLIRAPDSLKTLADFDPRVSIRLPSTSGSEGVPFRTVSEW |

| | | | | | | | | | | | | | | | | | |
|---|---|---|---|---|---|---|---|---|---|---|---|---|---|---|---|---|---|
| | | | | | | | | | | | | | | | | LESIKMQQYTEHFMAAGYTAIEKVVQMTNDDIKRIGVRLPGHQKRIAYSLLGLKDQVNTVGIPI | LESIKMQQYTEHFMAAGYTAIEKVVQMTNDDIKRIGVRLPGHQKRIAYSLLGLKDQVNTVGIPI |
| **2L2T (NMR)** | Low (Bio/Crystallographic) | - | Q15303 | ERBB4 | Q15303 | | A | B | 1 | PL.{2}AG.{3}G.{2}IL.{2}V | PL.{2}AG.{3}G.{2}IL.{2}V | 8 | 8 | 17 | 17 | MKPATGLWVWVSLLVAAGTVQPSDSQSVCAGTENKLSSLSDLEQQYRALRKYYENCEVVMGNLEITSIEHNRDLSFLRSVREVTGYVLVALNQFRYLPLENLRIIRGTKLYEDRYALAIFLNYRKDGNFGLQELGLKNLTEILNGGVYVDQNKFLCYADTIHWQDIVRNPWPSNLTLVSTNGSSGCGRCHKSCTGRCWGPTENHCQTLTRTVCAEQCDGRCYGPYVSDCCHRECAGGCSGPKDTDCFACMNFNDSGACVTQCPQTFVYNPTTFQLEHNFNAKYTYGAFCVKKCPHNFVVDSSSCVRACPSSKMEVEENGIKMCKPCTDICPKACDGIGTGSLMSAQTVDSSNIDKFINCTKINGNLIFLVTGIHGDPYNAIEAIDPEKLNVFRTVREITGFLNIQSWPPNMTDFSVFSNLVTIGGRVLYSGLSLLILKQQGITSLQFQSLKEISAGNIYITDNSNLCYYHTINWTTLFSTINQRIVIRDNRKAENCTAEGMVCNHLCSSDGCWGPGPDQCLSCRRFSRGRICIESCNLYDGEFREFENGSICVECDPQCEKMEDGLLTCHGPGPDNCTKCSHFKDGPNCVEKCPDGLQGANSFIFKYADPDRECHPCHPNCTQGCNGPTSHDCIYYPWTGHSTLPQHARTPLIAAGVIGGLFILVIVGLTFAVYVRRKSIKKKRALRRFLETELVEPLTPSGTAPNQAQLRILKETELKRVKVLGSGAFGTVYKGIWVPEGETVKIPVAIKILNETTGPKANVEFMDEALIMASMDHPHLVRLLGVCLSPTIQLVTQLMPHGCLLEYVHEHKDNIGSQLLLNWCVQIAKGMMYLEERRLVHRDLAARNVLVKSPNHVKITDFGLARLLEGDEKEYNADGGKMPI | MKPATGLWVWVSLLVAAGTVQPSDSQSVCAGTENKLSSLSDLEQQYRALRKYYENCEVVMGNLEITSIEHNRDLSFLRSVREVTGYVLVALNQFRYLPLENLRIIRGTKLYEDRYALAIFLNYRKDGNFGLQELGLKNLTEILNGGVYVDQNKFLCYADTIHWQDIVRNPWPSNLTLVSTNGSSGCGRCHKSCTGRCWGPTENHCQTLTRTVCAEQCDGRCYGPYVSDCCHRECAGGCSGPKDTDCFACMNFNDSGACVTQCPQTFVYNPTTFQLEHNFNAKYTYGAFCVKKCPHNFVVDSSSCVRACPSSKMEVEENGIKMCKPCTDICPKACDGIGTGSLMSAQTVDSSNIDKFINCTKINGNLIFLVTGIHGDPYNAIEAIDPEKLNVFRTVREITGFLNIQSWPPNMTDFSVFSNLVTIGGRVLYSGLSLLILKQQGITSLQFQSLKEISAGNIYITDNSNLCYYHTINWTTLFSTINQRIVIRDNRKAENCTAEGMVCNHLCSSDGCWGPGPDQCLSCRRFSRGRICIESCNLYDGEFREFENGSICVECDPQCEKMEDGLLTCHGPGPDNCTKCSHFKDGPNCVEKCPDGLQGANSFIFKYADPDRECHPCHPNCTQGCNGPTSHDCIYYPWTGHSTLPQHARTPLIAAGVIGGLFILVIVGLTFAVYVRRKSIKKKRALRRFLETELVEPLTPSGTAPNQAQLRILKETELKRVKVLGSGAFGTVYKGIWVPEGETVKIPVAIKILNETTGPKANVEFMDEALIMASMDHPHLVRLLGVCLSPTIQLVTQLMPHGCLLEYVHEHKDNIGSQLLLNWCVQIAKGMMYLEERRLVHRDLAARNVLVKSPNHVKITDFGLARLLEGDEKEYNADGGKMPI |

| | | | | | | | | | | | | | | | | | |
|---|---|---|---|---|---|---|---|---|---|---|---|---|---|---|---|---|---|
| | | | | | | | | | | | | | | | | KWMALECIHYRKFTHQS DVWSYGVTIWELMTFGG KPYDGIPTREIPDLLEKGE RLPQPPICTIDVYMVMVK CWMIDADSRPKFKELAA EFSRMARDPQRYLVIQG DDRMKLPSPNDSKFFQN LLDEEDLEDMMDAEEYL VPQAFNIPPPIYTSRARID SNRSEIGHSPPPAYTPMS GNQFVYRDGGFAAEQGV SVPYRAPTSTIPEAPVAQ GATAEIFDDSCCNGTLRK PVAPHVQEDSSTQRYSA DPTVFAPERSPRGELDEE GYMTPMRDKPKQEYLNP VEENPFVSRRKNGDLQAL DNPEYHNASNGPPKAED EYVNEPLYLNTFANTLGK AEYLKNNILSMPEKAKKA FDNPDYWNHSLPPRSTL QHPDYLQEYSTKYFYKQN GRIRPIVAENPEYLSEFSLK PGTVLPPPPYRHRNTVV | KWMALECIHYRKFTHQS DVWSYGVTIWELMTFGG KPYDGIPTREIPDLLEKGE RLPQPPICTIDVYMVMVK CWMIDADSRPKFKELAA EFSRMARDPQRYLVIQG DDRMKLPSPNDSKFFQN LLDEEDLEDMMDAEEYL VPQAFNIPPPIYTSRARID SNRSEIGHSPPPAYTPMS GNQFVYRDGGFAAEQGV SVPYRAPTSTIPEAPVAQ GATAEIFDDSCCNGTLRK PVAPHVQEDSSTQRYSA DPTVFAPERSPRGELDEE GYMTPMRDKPKQEYLNP VEENPFVSRRKNGDLQAL DNPEYHNASNGPPKAED EYVNEPLYLNTFANTLGK AEYLKNNILSMPEKAKKA FDNPDYWNHSLPPRSTL QHPDYLQEYSTKYFYKQN GRIRPIVAENPEYLSEFSLK PGTVLPPPPYRHRNTVV |
| **2L34 (NMR)** | Low (Bio>Crystallographic) | - | O43914 | TYROBP | O43914 | | A | B | 1 | LA.IV.GD.{2}LT.{2}I.{3}V | LA.IV.{2}D.{2}LT.{2}I.{3}V | 10 | 9 | 19 | 19 | MGGLEPCSRLLLLPLLLAV SGLRPVQAQAQSDCSCST VSPGVLAGIVMGDLVLTV LIALAVYFLGRLVPRGRGA AEAATRKQRITETESPYQE LQGQRSDVYSDLNTQRP YYK | MGGLEPCSRLLLLPLLLAV SGLRPVQAQAQSDCSCST VSPGVLAGIVMGDLVLTV LIALAVYFLGRLVPRGRGA AEAATRKQRITETESPYQE LQGQRSDVYSDLNTQRP YYK |
| **2L35** | Medi | - | P267 | KLR | O439 | TYRO | A | B | 1(A) | L.{2}I.{3}D.{2}LT.{2}I.{3}V.{16}I.{ | S.{2}VL.{2}IV.{2}D.{2}L.{ | 13 | 11 | 51 | 23 | MSKQRGTFSEVSLAQDP KRQQRKPKGNKSSISGTE QEIFQVELNLQNPSLNHQ GIDKIYDCQGLLPPPEKLT | MGGLEPCSRLLLLPLLLAV SGLRPVQAQAQSDCSCST VSPGVLAGIVMGDLVLTV LIALAVYFLGRLVPRGRGA |

| | | | | | | | | | | | | | | | | | |
|---|---|---|---|---|---|---|---|---|---|---|---|---|---|---|---|---|---|
| **(NMR)** | um (Bio) | | 17 | C2 | 14 | BP | | | ,1(B) | 2}VA.{3}K.{2}V.{3}F | 2}LI.{2}AV | | | | | AEVLGIICIVLMATVLKTIVLIPFLEQNNSSPNTRTQKARHCGHCPEEWITYSNSCYYIGKERRTWEESLLACTSKNSSLLSIDNEEEMKFLASILPSSWIGVFRNSSHHPWVTINGLAFKHKIKDSDNAELNCAVLQVNRLKSAQCGSSMIYHCKHKL | AEAATRKQRITETESPYQELQGQRSDVYSDLNTQRPYYK |
| **2L6W (NMR)** | Low (Bio/Crystallographic) | - | P09619 | PDGFRB | P09619 | | A | B | 1 | VV.{2}A.{2}A.{2}VL.{2}I.{3}I | VV.{2}A.{2}AL.VL.{2}IS.{2}I | 8 | 10 | 19 | 19 | MRLPGAMPALALKGELLLLSLLLLLEPQISQGLVVTPPGPELVLNVSSTFVLTCSGSAPVVWERMSQEPPQEMAKAQDGTFSSVLTLTNLTGLDTGEYFCTHNDSRGLETDERKRLYIFVPDPTVGFLPNDAEELFIFLTEITEITIPCRVTDPQLVVTLHEKKGDVALPVPYDHQRGFSGIFEDRSYICKTTIGDREVDSDAYYVYRLQVSSINVSVNAVQTVVRQGENITLMCIVIGNEVVNFEWTYPRKESGRLVEPVTDFLLDMPYHIRSILHIPSAELEDSGTYTCNVTESVNDHQDEKAINITVVESGYVRLLGEVGTLQFAELHRSRTLQVVFEAYPPPTVLWFKDNRTLGDSSAGEIALSTRNVSETRYVSELTLVRVKVAEAGHYTMRAFHEDAEVQLSFQLQINVPVRVLELSESHPDSGEQTVRCRGRGMPQPNIIWSACRDLKRCPRELPPTLLGNSSEEESQLETNVTYWEEEQEFEVVSTLRLQHVDRPLSVRCTLRNAVGQDTQEVIVVPHSLPFKVVVISAILALVVLTIISLIILIMLWQKKPRYEIRWKVIESVSSDGHEYIYVDPMQLPYDSTWELPRDQLVLGRTLGSGAFGQVVEATAHGLSHSQATMKVAVKMLKSTARSSEKQALMSELKIMSHLGPHLNVVNLLGACTKGGPIYIITEYCRYGDLVDYLHRNKHTFLQHHSDKRRPPSAELYSNALPVGLPLPSHVSLTGESDGGYMDMSKDESVDYVPMLDMKGDVKYADIESSNYMAPYDNYVPSAPERTCRATLINESPVLSYMDLV | MRLPGAMPALALKGELLLLSLLLLLEPQISQGLVVTPPGPELVLNVSSTFVLTCSGSAPVVWERMSQEPPQEMAKAQDGTFSSVLTLTNLTGLDTGEYFCTHNDSRGLETDERKRLYIFVPDPTVGFLPNDAEELFIFLTEITEITIPCRVTDPQLVVTLHEKKGDVALPVPYDHQRGFSGIFEDRSYICKTTIGDREVDSDAYYVYRLQVSSINVSVNAVQTVVRQGENITLMCIVIGNEVVNFEWTYPRKESGRLVEPVTDFLLDMPYHIRSILHIPSAELEDSGTYTCNVTESVNDHQDEKAINITVVESGYVRLLGEVGTLQFAELHRSRTLQVVFEAYPPPTVLWFKDNRTLGDSSAGEIALSTRNVSETRYVSELTLVRVKVAEAGHYTMRAFHEDAEVQLSFQLQINVPVRVLELSESHPDSGEQTVRCRGRGMPQPNIIWSACRDLKRCPRELPPTLLGNSSEEESQLETNVTYWEEEQEFEVVSTLRLQHVDRPLSVRCTLRNAVGQDTQEVIVVPHSLPFKVVVISAILALVVLTIISLIILIMLWQKKPRYEIRWKVIESVSSDGHEYIYVDPMQLPYDSTWELPRDQLVLGRTLGSGAFGQVVEATAHGLSHSQATMKVAVKMLKSTARSSEKQALMSELKIMSHLGPHLNVVNLLGACTKGGPIYIITEYCRYGDLVDYLHRNKHTFLQHHSDKRRPPSAELYSNALPVGLPLPSHVSLTGESDGGYMDMSKDESVDYVPMLDMKGDVKYADIESSNYMAPYDNYVPSAPERTCRATLINESPVLSYMDLV |

| | | | | | | | | | | | | | | | | | |
|---|---|---|---|---|---|---|---|---|---|---|---|---|---|---|---|---|---|
| | | | | | | | | | | | | | | | | GFSYQVANGMEFLASKNCVHRDLAARNVLICEGKLVKICDFGLARDIMRDSNYISKGSTFLPLKWMAPESIFNSLYTTLSDVWSFGILLWEIFTLGGTPYPELPMNEQFYNAIKRGYRMAQPAHASDEIYEIMQKCWEEKFEIRPPFSQLVLLLERLLGEGYKKKYQQVDEEFLRSDHPAILRSQARLPGFHGLRSPLDTSSVLYTAVQPNEGDNDYIIPLPDPKPEVADEGPLEGSPSLASSTLNEVNTSSTISCDSPLEPQDEPEPEPQLELQVEPEPELEQLPDSGCPAPRAEAEDSFL | GFSYQVANGMEFLASKNCVHRDLAARNVLICEGKLVKICDFGLARDIMRDSNYISKGSTFLPLKWMAPESIFNSLYTTLSDVWSFGILLWEIFTLGGTPYPELPMNEQFYNAIKRGYRMAQPAHASDEIYEIMQKCWEEKFEIRPPFSQLVLLLERLLGEGYKKKYQQVDEEFLRSDHPAILRSQARLPGFHGLRSPLDTSSVLYTAVQPNEGDNDYIIPLPDPKPEVADEGPLEGSPSLASSTLNEVNTSSTISCDSPLEPQDEPEPEPQLELQVEPEPELEQLPDSGCPAPRAEAEDSFL |
| **2L9U (NMR)** | Low (Bio/Crystallographic) | - | P21860 | ERBB3 | P21860 | | A | B | 1 | I.{2}LV.IF.{2}L.{3}F | I.{2}LV.IF.{2}L.{3}F | 7 | 7 | 15 | 15 | MRANDALQVLGLLFSLARGSEVGNSQAVCPGTLNGLSVTGDAENQYQTLYKLYERCEVVMGNLEIVLTGHNADLSFLQWIREVTGYVLVAMNEFSTLPLPNLRVVRGTQVYDGKFAIFVMLNYNTNSSHALRQLRLTQLTEILSGGVYIEKNDKLCHMDTIDWRDIVRDRDAEIVVKDNGRSCPPCHEVCKGRCWGPGSEDCQTLTKTICAPQCNGHCFGPNPNQCCHDECAGGCSGPQDTDCFACRHFNDSGACVPRCPQPLVYNKLTFQLEPNPHTKYQYGGVCVASCPHNFVVDQTSCVRACPPDKMEVDKNGLKMCEPCGGLCPKACEGTGSGSRFQTVDSSNIDGFVNCTKILGNLDFLITGLNGDPWHKIPALDPEKLNVFRTVREITGYLNIQSWPPHMHNFSVFSNLTTIGGRSLYNRGFSLLIMKNLNVTSLGFRSLKEISAGRIYISANRQLCYHHSLNWTKVLRGPTEERLDIKHNRPRRDCVAEGKVCDPLCSSGGCWGPGPGQCLSCRNYSRGGVCVTHCNFLNGEPREFAHEAECFSCHPECQPMEGTATCNGSGSDTCAQCAHFRDGPHCVSSCPHGVLGAKGPIYKYPDVQNECRPCHENCTQGCKGPELQDCLGQTLVLIGKT | MRANDALQVLGLLFSLARGSEVGNSQAVCPGTLNGLSVTGDAENQYQTLYKLYERCEVVMGNLEIVLTGHNADLSFLQWIREVTGYVLVAMNEFSTLPLPNLRVVRGTQVYDGKFAIFVMLNYNTNSSHALRQLRLTQLTEILSGGVYIEKNDKLCHMDTIDWRDIVRDRDAEIVVKDNGRSCPPCHEVCKGRCWGPGSEDCQTLTKTICAPQCNGHCFGPNPNQCCHDECAGGCSGPQDTDCFACRHFNDSGACVPRCPQPLVYNKLTFQLEPNPHTKYQYGGVCVASCPHNFVVDQTSCVRACPPDKMEVDKNGLKMCEPCGGLCPKACEGTGSGSRFQTVDSSNIDGFVNCTKILGNLDFLITGLNGDPWHKIPALDPEKLNVFRTVREITGYLNIQSWPPHMHNFSVFSNLTTIGGRSLYNRGFSLLIMKNLNVTSLGFRSLKEISAGRIYISANRQLCYHHSLNWTKVLRGPTEERLDIKHNRPRRDCVAEGKVCDPLCSSGGCWGPGPGQCLSCRNYSRGGVCVTHCNFLNGEPREFAHEAECFSCHPECQPMEGTATCNGSGSDTCAQCAHFRDGPHCVSSCPHGVLGAKGPIYKYPDVQNECRPCHENCTQGCKGPELQDCLGQTLVLIGKT |

| | | | | | | | | | | | | | | | | | |
|---|---|---|---|---|---|---|---|---|---|---|---|---|---|---|---|---|---|
| | | | | | | | | | | | | | | | | HLTMALTVIAGLVVIFMM<br>LGGTFLYWRGRRIQNKR<br>AMRRYLERGESIEPLDPSE<br>KANKVLARIFKETELRKLK<br>VLGSGVFGTVHKGVWIP<br>EGESIKIPVCIKVIEDKSGR<br>QSFQAVTDHMLAIGSLD<br>HAHIVRLLGLCPGSSLQLV<br>TQYLPLGSLLDHVRQHRG<br>ALGPQLLLNWGVQIAKG<br>MYYLEEHGMVHRNLAAR<br>NVLLKSPSQVQVADFGV<br>ADLLPPDDKQLLYSEAKTP<br>IKWMALESIHFGKYTHQS<br>DVWSYGVTVWELMTFG<br>AEPYAGLRLAEVPDLLEK<br>GERLAQPQICTIDVYMV<br>MVKCWMIDENIRPTFKE<br>LANEFTRMARDPPRYLVI<br>KRESGPGIAPGPEPHGLT<br>NKKLEEVELEPELDLDLDL<br>EAEEDNLATTTLGSALSLP<br>VGTLNRPRGSQSLLSPSS<br>GYMPMNQGNLGESCQE<br>SAVSGSSERCPRPVSLHP<br>MPRGCLASESSEGHVTGS<br>EAELQEKVSMCRSRSRSR<br>SPRPRGDSAYHSQRHSLL<br>TPVTPLSPPGLEEEDVNG<br>YVMPDTHLKGTPSSREGT<br>LSSVGLSSVLGTEEEDEDE<br>EYEYMNRRRRHSPPHPP<br>RPSSLEELGYEYMDVGSD<br>LSASLGSTQSCPLHPVPIM<br>PTAGTTPDEDYEYMNRQ<br>RDGGGPGGDYAAMGAC<br>PASEQGYEEMRAFQGPG<br>HQAPHVHYARLKTLRSLE<br>ATDSAFDNPDYWHSRLF<br>PKANAQRT | HLTMALTVIAGLVVIFMM<br>LGGTFLYWRGRRIQNKR<br>AMRRYLERGESIEPLDPSE<br>KANKVLARIFKETELRKLK<br>VLGSGVFGTVHKGVWIP<br>EGESIKIPVCIKVIEDKSGR<br>QSFQAVTDHMLAIGSLD<br>HAHIVRLLGLCPGSSLQLV<br>TQYLPLGSLLDHVRQHRG<br>ALGPQLLLNWGVQIAKG<br>MYYLEEHGMVHRNLAAR<br>NVLLKSPSQVQVADFGV<br>ADLLPPDDKQLLYSEAKTP<br>IKWMALESIHFGKYTHQS<br>DVWSYGVTVWELMTFG<br>AEPYAGLRLAEVPDLLEK<br>GERLAQPQICTIDVYMV<br>MVKCWMIDENIRPTFKE<br>LANEFTRMARDPPRYLVI<br>KRESGPGIAPGPEPHGLT<br>NKKLEEVELEPELDLDLDL<br>EAEEDNLATTTLGSALSLP<br>VGTLNRPRGSQSLLSPSS<br>GYMPMNQGNLGESCQE<br>SAVSGSSERCPRPVSLHP<br>MPRGCLASESSEGHVTGS<br>EAELQEKVSMCRSRSRSR<br>SPRPRGDSAYHSQRHSLL<br>TPVTPLSPPGLEEEDVNG<br>YVMPDTHLKGTPSSREGT<br>LSSVGLSSVLGTEEEDEDE<br>EYEYMNRRRRHSPPHPP<br>RPSSLEELGYEYMDVGSD<br>LSASLGSTQSCPLHPVPIM<br>PTAGTTPDEDYEYMNRQ<br>RDGGGPGGDYAAMGAC<br>PASEQGYEEMRAFQGPG<br>HQAPHVHYARLKTLRSLE<br>ATDSAFDNPDYWHSRLF<br>PKANAQRT |
| **2LOH (NMR)** | Low (Bio>Crys | 2LLM | P05067 | APP | P05067 | | A | B | 1 | I.{2}LM.{2}G.{3}A.{2}IV.{2}L | I.{2}LM.{2}G.{3}A.{2}IV.{2}L | 8 | 8 | 19 | 19 | MLPGLALLLLAAWTARAL<br>EVPTDGNAGLLAEPQIA<br>MFCGRLNMHMNVQNG<br>KWDSDPSGTKTCIDTKEG<br>ILQYCQEVYPELQITNVVE<br>ANQPVTIQNWCKRGRK<br>QCKTHPHFVIPYRCLVGE<br>FVSDALLVPDKCKFLHQE<br>RMDVCETHLHWHTVAK<br>ETCSEKSTNLHDYGMLLP<br>CGIDKFRGVEFVCCPLAEE<br>SDNVDSADAEEDDSDVW<br>WGGADTDYADGSEDKV | MLPGLALLLLAAWTARAL<br>EVPTDGNAGLLAEPQIA<br>MFCGRLNMHMNVQNG<br>KWDSDPSGTKTCIDTKEG<br>ILQYCQEVYPELQITNVVE<br>ANQPVTIQNWCKRGRK<br>QCKTHPHFVIPYRCLVGE<br>FVSDALLVPDKCKFLHQE<br>RMDVCETHLHWHTVAK<br>ETCSEKSTNLHDYGMLLP<br>CGIDKFRGVEFVCCPLAEE<br>SDNVDSADAEEDDSDVW<br>WGGADTDYADGSEDKV |

| | | | | | | | | | | | | | | | | | |
|---|---|---|---|---|---|---|---|---|---|---|---|---|---|---|---|---|---|
| | tallographic) | | | | | | | | | | | | | | | VEVAEEEEVAEVEEEEADDDEDDEDGDEVEEEAEEPYEEATERTTSIATTTTTTTESVEEVVREVCSEQAETGPCRAMISRWYFDVTEGKCAPFFYGGCGGNRNNFDTEEYCMAVCGSAMSQSLLKTTQEPLARDPVKLPTTAASTPDAVDKYLETPGDENEHAHFQKAKERLEAKHRERMSQVMREWEEAERQAKNLPKADKKAVIQHFQEKVESLEQEAANERQQLVETHMARVEAMLNDRRRLALENYITALQAVPPRPRHVFNMLKKYVRAEQKDRQHTLKHFEHVRMVDPKKAAQIRSQVMTHLRVIYERMNQSLSLLYNVPAVAEEIQDEVDELLQKEQNYSDDVLANMISEPRISYGNDALMPSLTETKTTVELLPVNGEFSLDDLQPWHSFGADSVPANTENEVEPVDARPAADRGLTTRPGSGLTNIKTEEISEVKMDAEFRHDSGYEVHHQKLVFFAEDVGSNKGAIIGLMVGGVVIATVIVITLVMLKKKQYTSIHHGVVEVDAAVTPEERHLSKMQQNGYENPTYKFFEQMQN | VEVAEEEEVAEVEEEEADDDEDDEDGDEVEEEAEEPYEEATERTTSIATTTTTTTESVEEVVREVCSEQAETGPCRAMISRWYFDVTEGKCAPFFYGGCGGNRNNFDTEEYCMAVCGSAMSQSLLKTTQEPLARDPVKLPTTAASTPDAVDKYLETPGDENEHAHFQKAKERLEAKHRERMSQVMREWEEAERQAKNLPKADKKAVIQHFQEKVESLEQEAANERQQLVETHMARVEAMLNDRRRLALENYITALQAVPPRPRHVFNMLKKYVRAEQKDRQHTLKHFEHVRMVDPKKAAQIRSQVMTHLRVIYERMNQSLSLLYNVPAVAEEIQDEVDELLQKEQNYSDDVLANMISEPRISYGNDALMPSLTETKTTVELLPVNGEFSLDDLQPWHSFGADSVPANTENEVEPVDARPAADRGLTTRPGSGLTNIKTEEISEVKMDAEFRHDSGYEVHHQKLVFFAEDVGSNKGAIIGLMVGGVVIATVIVITLVMLKKKQYTSIHHGVVEVDAAVTPEERHLSKMQQNGYENPTYKFFEQMQN |
| **2LZL (NMR)** | Low (Bio/Crystallograph | - | P22607 | FGFR3 | P22607 | | A | B | 1 | L.YGV.FF.{2}I.{6}TL | L.YGV.FF.{2}I.{6}TL | 9 | 9 | 19 | 19 | MGAPACALALCVAVAIVAGASSESLGTEQRVVGRAAEVPGPEPGQQEQLVFGSGDAVELSCPPPGGGPMGPTVWVKDGTGLVPSERVLVGPQRLQVLNASHEDSGAYSCRQRLTQRVLCHFSVRVTDAPSSGDDEDGEDEAEDTGVDTGAPYWTRPERMDKKLLAVPAANTVRFRCPAAGNPTPSISWLKNGREFRGEHRIGGIKLRHQQWSLVMESVVPSDRGNYTCVVENKFGSIRQTYTLDVLERSPHRPILQAGLPANQTAVLGSDVEFHCKVYSDAQPHIQWLKHVEVNGSKVGPDGTPYVTVLKTAGANTTDKELEVLSLHNVTFEDAGEYTCLAGNSIGFSHHSAWLVVLPAEEELVEADEAGSVYAGILSYGVGFFLFIL | MGAPACALALCVAVAIVAGASSESLGTEQRVVGRAAEVPGPEPGQQEQLVFGSGDAVELSCPPPGGGPMGPTVWVKDGTGLVPSERVLVGPQRLQVLNASHEDSGAYSCRQRLTQRVLCHFSVRVTDAPSSGDDEDGEDEAEDTGVDTGAPYWTRPERMDKKLLAVPAANTVRFRCPAAGNPTPSISWLKNGREFRGEHRIGGIKLRHQQWSLVMESVVPSDRGNYTCVVENKFGSIRQTYTLDVLERSPHRPILQAGLPANQTAVLGSDVEFHCKVYSDAQPHIQWLKHVEVNGSKVGPDGTPYVTVLKTAGANTTDKELEVLSLHNVTFEDAGEYTCLAGNSIGFSHHSAWLVVLPAEEELVEADEAGSVYAGILSYGVGFFLFIL |

| | | | | | | | | | | | | | | | | | |
|---|---|---|---|---|---|---|---|---|---|---|---|---|---|---|---|---|---|
| | ic) | | | | | | | | | | | | | | | VVAAVTLCRLRSPPKKGL<br>GSPTVHKISRFPLKRQVSL<br>ESNASMSSNTPLVRIARLS<br>SGEGPTLANVSELELPAD<br>PKWELSRARLTLGKPLGE<br>GCFGQVVMAEAIGIDKD<br>RAAKPVTVAVKMLKDDA<br>TDKDLSDLVSEMEMMK<br>MIGKHKNIINLLGACTQG<br>GPLYVLVEYAAKGNLREFL<br>RARRPPGLDYSFDTCKPP<br>EEQLTFKDLVSCAYQVAR<br>GMEYLASQKCIHRDLAAR<br>NVLVTEDNVMKIADFGL<br>ARDVHNLDYYKKTTNGRL<br>PVKWMAPEALFDRVYTH<br>QSDVWSFGVLLWEIFTLG<br>GSPYPGIPVEELFKLLKEG<br>HRMDKPANCTHDLYMI<br>MRECWHAAPSQRPTFK<br>QLVEDLDRVLTVTSTDEYL<br>DLSAPFEQYSPGGQDTPS<br>SSSSGDDSVFAHDLLPPA<br>PPSSGGSRT | VVAAVTLCRLRSPPKKGL<br>GSPTVHKISRFPLKRQVSL<br>ESNASMSSNTPLVRIARLS<br>SGEGPTLANVSELELPAD<br>PKWELSRARLTLGKPLGE<br>GCFGQVVMAEAIGIDKD<br>RAAKPVTVAVKMLKDDA<br>TDKDLSDLVSEMEMMK<br>MIGKHKNIINLLGACTQG<br>GPLYVLVEYAAKGNLREFL<br>RARRPPGLDYSFDTCKPP<br>EEQLTFKDLVSCAYQVAR<br>GMEYLASQKCIHRDLAAR<br>NVLVTEDNVMKIADFGL<br>ARDVHNLDYYKKTTNGRL<br>PVKWMAPEALFDRVYTH<br>QSDVWSFGVLLWEIFTLG<br>GSPYPGIPVEELFKLLKEG<br>HRMDKPANCTHDLYMI<br>MRECWHAAPSQRPTFK<br>QLVEDLDRVLTVTSTDEYL<br>DLSAPFEQYSPGGQDTPS<br>SSSSGDDSVFAHDLLPPA<br>PPSSGGSRT |
| **2M0B (NMR)** | Low (Bio>Crystallographic) | - | P00533 | EGFR | P00533 | | A | B | 1 | M.{3}L.{2}LL.{2}AL.{2}GL | M.{3}L.{2}LL.{2}AL.{2}GL | 8 | 8 | 17 | 17 | MRPSGTAGAALLALLAAL<br>CPASRALEEKKVCQGTSN<br>KLTQLGTFEDHFLSLQRM<br>FNNCEVVLGNLEITYVQR<br>NYDLSFLKTIQEVAGYVLI<br>ALNTVERIPLENLQIIRGN<br>MYYENSYALAVLSNYDA<br>NKTGLKELPMRNLQEILH<br>GAVRFSNNPALCNVESIQ<br>WRDIVSSDFLSNMSMDF<br>QNHLGSCQKCDPSCPNG<br>SCWGAGEENCQKLTKIIC<br>AQQCSGRCRGKSPSDCC<br>HNQCAAGCTGPRESDCL<br>VCRKFRDEATCKDTCPPL<br>MLYNPTTYQMDVNPEG<br>KYSFGATCVKKCPRNYVV<br>TDHGSCVRACGADSYEM<br>EEDGVRKCKKCEGPCRKV<br>CNGIGIGEFKDSLSINATNI<br>KHFKNCTSISGDLHILPVA<br>FRGDSFTHTPPLDPQELDI<br>LKTVKEITGFLLIQAWPEN<br>RTDLHAFENLEIIRGRTKQ<br>HGQFSLAVVSLNITSLGLR<br>SLKEISDGDVIISGNKNLCY<br>ANTINWKKLFGTSGQKTK<br>IISNRGENSCKATGQVCH<br>ALCSPEGCWGPEPRDCV | MRPSGTAGAALLALLAAL<br>CPASRALEEKKVCQGTSN<br>KLTQLGTFEDHFLSLQRM<br>FNNCEVVLGNLEITYVQR<br>NYDLSFLKTIQEVAGYVLI<br>ALNTVERIPLENLQIIRGN<br>MYYENSYALAVLSNYDA<br>NKTGLKELPMRNLQEILH<br>GAVRFSNNPALCNVESIQ<br>WRDIVSSDFLSNMSMDF<br>QNHLGSCQKCDPSCPNG<br>SCWGAGEENCQKLTKIIC<br>AQQCSGRCRGKSPSDCC<br>HNQCAAGCTGPRESDCL<br>VCRKFRDEATCKDTCPPL<br>MLYNPTTYQMDVNPEG<br>KYSFGATCVKKCPRNYVV<br>TDHGSCVRACGADSYEM<br>EEDGVRKCKKCEGPCRKV<br>CNGIGIGEFKDSLSINATNI<br>KHFKNCTSISGDLHILPVA<br>FRGDSFTHTPPLDPQELDI<br>LKTVKEITGFLLIQAWPEN<br>RTDLHAFENLEIIRGRTKQ<br>HGQFSLAVVSLNITSLGLR<br>SLKEISDGDVIISGNKNLCY<br>ANTINWKKLFGTSGQKTK<br>IISNRGENSCKATGQVCH<br>ALCSPEGCWGPEPRDCV |

| | | | | | | | | | | | | | | | | | |
|---|---|---|---|---|---|---|---|---|---|---|---|---|---|---|---|---|---|
| | | | | | | | | | | | | | | | | SCRNVSRGRECVDKCNLLEGEPREFVENSECIQCHPECLPQAMNITCTGRGPDNCIQCAHYIDGPHCVKTCPAGVMGENNTLVWKYADAGHVCHLCHPNCTYGCTGPGLEGCPTNGPKIPSIATGMVGALLLLLVVALGIGLFMRRRHIVRKRTLRRLLQERELVEPLTPSGEAPNQALLRILKETEFKKIKVLGSGAFGTVYKGLWIPEGEKVKIPVAIKELREATSPKANKEILDEAYVMASVDNPHVCRLLGICLTSTVQLITQLMPFGCLLDYVREHKDNIGSQYLLNWCVQIAKGMNYLEDRRLVHRDLAARNVLVKTPQHVKITDFGLAKLLGAEEKEYHAEGGKVPIKWMALESILHRIYTHQSDVWSYGVTVWELMTFGSKPYDGIPASEISSILEKGERLPQPPICTIDVYMIMVKCWMIDADSRPKFRELIIEFSKMARDPQRYLVIQGDERMHLPSPTDSNFYRALMDEEDMDDVVDADEYLIPQQGFFSSPSTSRTPLLSSLSATSNNSTVACIDRNGLQSCPIKEDSFLQRYSSDPTGALTEDSIDDTFLPVPEYINQSVPKRPAGSVQNPVYHNQPLNPAPSRDPHYQDPHSTAVGNPEYLNTVQPTCVNSTFDSPAHWAQKGSHQISLDNPDYQQDFFPKEAKPNGIFKGSTAENAEYLRVAPQSSEFIGA | SCRNVSRGRECVDKCNLLEGEPREFVENSECIQCHPECLPQAMNITCTGRGPDNCIQCAHYIDGPHCVKTCPAGVMGENNTLVWKYADAGHVCHLCHPNCTYGCTGPGLEGCPTNGPKIPSIATGMVGALLLLLVVALGIGLFMRRRHIVRKRTLRRLLQERELVEPLTPSGEAPNQALLRILKETEFKKIKVLGSGAFGTVYKGLWIPEGEKVKIPVAIKELREATSPKANKEILDEAYVMASVDNPHVCRLLGICLTSTVQLITQLMPFGCLLDYVREHKDNIGSQYLLNWCVQIAKGMNYLEDRRLVHRDLAARNVLVKTPQHVKITDFGLAKLLGAEEKEYHAEGGKVPIKWMALESILHRIYTHQSDVWSYGVTVWELMTFGSKPYDGIPASEISSILEKGERLPQPPICTIDVYMIMVKCWMIDADSRPKFRELIIEFSKMARDPQRYLVIQGDERMHLPSPTDSNFYRALMDEEDMDDVVDADEYLIPQQGFFSSPSTSRTPLLSSLSATSNNSTVACIDRNGLQSCPIKEDSFLQRYSSDPTGALTEDSIDDTFLPVPEYINQSVPKRPAGSVQNPVYHNQPLNPAPSRDPHYQDPHSTAVGNPEYLNTVQPTCVNSTFDSPAHWAQKGSHQISLDNPDYQQDFFPKEAKPNGIFKGSTAENAEYLRVAPQSSEFIGA |
| **2M59 (NMR)** | Low (Bio/Cryst | - | P35968 | KDR | P35968 | | A | B | 1 | EI.IL.{9}FW.LL.{2}I | EI.IL.{9}FW.LL.{2}I | 9 | 9 | 22 | 22 | MQSKVLLAVALWLCVETRAASVGLPSVSLDLPRLSIQKDILTIKANTTLQITCRGQRDLDWLWPNNQSGSEQRVEVTECSDGLFCKTLTIPKVIGNDTGAYKCFYRETDLASVIYVYVQDYRSPFIASVSDQHGVVYITENKNKTVVIPCLGSISNLNVSLCARYPEKRFVPDGNRISWDSKKGFTIPSYMISYAGMVFCEAKINDESYQSIMYIVVVVGYRIYDVVLSPSHGIELSVGEKLVLNCTARTELNVGI | MQSKVLLAVALWLCVETRAASVGLPSVSLDLPRLSIQKDILTIKANTTLQITCRGQRDLDWLWPNNQSGSEQRVEVTECSDGLFCKTLTIPKVIGNDTGAYKCFYRETDLASVIYVYVQDYRSPFIASVSDQHGVVYITENKNKTVVIPCLGSISNLNVSLCARYPEKRFVPDGNRISWDSKKGFTIPSYMISYAGMVFCEAKINDESYQSIMYIVVVVGYRIYDVVLSPSHGIELSVGEKLVLNCTARTELNVGI |

| | | | | | | | | | | | | | | | | | |
|---|---|---|---|---|---|---|---|---|---|---|---|---|---|---|---|---|---|
| | a<br>ll<br>o<br>g<br>r<br>a<br>p<br>h<br>i<br>c<br>) | | | | | | | | | | | | | | | DFNWEYPSSKHQHKKLV<br>NRDLKTQSGSEMKKFLST<br>LTIDGVTRSDQGLYTCAA<br>SSGLMTKKNSTFVRVHEK<br>PFVAFGSGMESLVEATVG<br>ERVRIPAKYLGYPPPEIKW<br>YKNGIPLESNHTIKAGHVL<br>TIMEVSERDTGNYTVILTN<br>PISKEKQSHVVSLVVYVPP<br>QIGEKSLISPVDSYQYGTT<br>QTLTCTVYAIPPPHHIHW<br>YWQLEEECANEPSQAVS<br>VTNPYPCEEWRSVEDFQ<br>GGNKIEVNKNQFALIEGK<br>NKTVSTLVIQAANVSALY<br>KCEAVNKVGRGERVISFH<br>VTRGPEITLQPDMQPTE<br>QESVSLWCTADRSTFENL<br>TWYKLGPQPLPIHVGELP<br>TPVCKNLDTLWKLNATM<br>FSNSTNDILIMELKNASLQ<br>DQGDYVCLAQDRKTKKR<br>HCVVRQLTVLERVAPTIT<br>GNLENQTTSIGESIEVSCT<br>ASGNPPPQIMWFKDNET<br>LVEDSGIVLKDGNRNLTIR<br>RVRKEDEGLYTCQACSVL<br>GCAKVEAFFIIEGAQEKTN<br>LEIIILVGTAVIAMFFWLLL<br>VIILRTVKRANGGELKTGY<br>LSIVMDPDELPLDEHCERL<br>PYDASKWEFPRDRLKLGK<br>PLGRGAFGQVIEADAFGI<br>DKTATCRTVAVKMLKEG<br>ATHSEHRALMSELKILIHI<br>GHHLNVVNLLGACTKPG<br>GPLMVIVEFCKFGNLSTYL<br>RSKRNEFVPYKTKGARFR<br>QGKDYVGAIPVDLKRRLD<br>SITSSQSSASSGFVEEKSLS<br>DVEEEEAPEDLYKDFLTLE<br>HLICYSFQVAKGMEFLAS<br>RKCIHRDLAARNILLSEKN<br>VVKICDFGLARDIYKDPDY<br>VRKGDARLPLKWMAPET<br>IFDRVYTIQSDVWSFGVLL<br>WEIFSLGASPYPGVKIDEE<br>FCRRLKEGTRMRAPDYTT<br>PEMYQTMLDCWHGEPS<br>QRPTFSELVEHLGNLLQA<br>NAQQDGKDYIVLPISETLS<br>MEEDSGLSLPTSPVSCME<br>EEEVCDPKFHYDNTAGIS | DFNWEYPSSKHQHKKLV<br>NRDLKTQSGSEMKKFLST<br>LTIDGVTRSDQGLYTCAA<br>SSGLMTKKNSTFVRVHEK<br>PFVAFGSGMESLVEATVG<br>ERVRIPAKYLGYPPPEIKW<br>YKNGIPLESNHTIKAGHVL<br>TIMEVSERDTGNYTVILTN<br>PISKEKQSHVVSLVVYVPP<br>QIGEKSLISPVDSYQYGTT<br>QTLTCTVYAIPPPHHIHW<br>YWQLEEECANEPSQAVS<br>VTNPYPCEEWRSVEDFQ<br>GGNKIEVNKNQFALIEGK<br>NKTVSTLVIQAANVSALY<br>KCEAVNKVGRGERVISFH<br>VTRGPEITLQPDMQPTE<br>QESVSLWCTADRSTFENL<br>TWYKLGPQPLPIHVGELP<br>TPVCKNLDTLWKLNATM<br>FSNSTNDILIMELKNASLQ<br>DQGDYVCLAQDRKTKKR<br>HCVVRQLTVLERVAPTIT<br>GNLENQTTSIGESIEVSCT<br>ASGNPPPQIMWFKDNET<br>LVEDSGIVLKDGNRNLTIR<br>RVRKEDEGLYTCQACSVL<br>GCAKVEAFFIIEGAQEKTN<br>LEIIILVGTAVIAMFFWLLL<br>VIILRTVKRANGGELKTGY<br>LSIVMDPDELPLDEHCERL<br>PYDASKWEFPRDRLKLGK<br>PLGRGAFGQVIEADAFGI<br>DKTATCRTVAVKMLKEG<br>ATHSEHRALMSELKILIHI<br>GHHLNVVNLLGACTKPG<br>GPLMVIVEFCKFGNLSTYL<br>RSKRNEFVPYKTKGARFR<br>QGKDYVGAIPVDLKRRLD<br>SITSSQSSASSGFVEEKSLS<br>DVEEEEAPEDLYKDFLTLE<br>HLICYSFQVAKGMEFLAS<br>RKCIHRDLAARNILLSEKN<br>VVKICDFGLARDIYKDPDY<br>VRKGDARLPLKWMAPET<br>IFDRVYTIQSDVWSFGVLL<br>WEIFSLGASPYPGVKIDEE<br>FCRRLKEGTRMRAPDYTT<br>PEMYQTMLDCWHGEPS<br>QRPTFSELVEHLGNLLQA<br>NAQQDGKDYIVLPISETLS<br>MEEDSGLSLPTSPVSCME<br>EEEVCDPKFHYDNTAGIS |

| | | | | | | | | | | | | | | | | | |
|---|---|---|---|---|---|---|---|---|---|---|---|---|---|---|---|---|---|
| | | | | | | | | | | | | | | | | QYLQNSKRKSRPVSVKTFEDIPLEEPEVKVIPDDNQTDSGMVLASEELKTLEDRTKLSPSFGGMVPSKSRESVASEGSNQTSGYQSGYHSDDTDTTVYSSEEAELLKLIEIGVQTGSTAQILQPDSGTTLSSPPV | QYLQNSKRKSRPVSVKTFEDIPLEEPEVKVIPDDNQTDSGMVLASEELKTLEDRTKLSPSFGGMVPSKSRESVASEGSNQTSGYQSGYHSDDTDTTVYSSEEAELLKLIEIGVQTGSTAQILQPDSGTTLSSPPV |
| **2MK9 (NMR)** | Low (Bio/Crystallographic) | - | O15455 | TLR3 | O15455 | | A | B | 1 | S.{2}LI.{2}FI.{2}L | S.{2}LI.{2}FI.{2}L | 6 | 6 | 12 | 12 | MRQTLPCIYFWGGLLPFGMLCASSTTKCTVSHEVADCSHLKLTQVPDDLPTNITVLNLTHNQLRRLPAANFTRYSQLTSLDVGFNTISKLEPELCQKLPMLKVLNLQHNELSQLSDKTFAFCTNLTELHLMSNSIQKIKNNPFVKQKNLITLDLSHNGLSSTKLGTQVQLENLQELLLSNNKIQALKSEELDIFANSSLKKLELSSNQIKEFSPGCFHAIGRLFGLFLNNVQLGPSLTEKLCLELANTSIRNLSLSNSQLSTTSNTTFLGLKWTNLTMLDLSYNNLNVVGNDSFAWLPQLEYFFLEYNNIQHLFSHSLHGLFNVRYLNLKRSFTKQSISLASLPKIDDFSFQWLKCLEHLNMEDNDIPGIKSNMFTGLINLKYLSLSNSFTSLRTLTNETFVSLAHSPLHILNLTKNKISKIESDAFSWLGHLEVLDLGLNEIGQELTGQEWRGLENIFEIYLSYNKYLQLTRNSFALVPSLQRLMLRRVALKNVDSSPSPFQPLRNLTILDLSNNNIANINDDMLEGLEKLEILDLQHNNLARLWKHANPGGPIYFLKGLSHLHILNLESNGFDEIPVEVFKDLFELKIIDLGLNNLNTLPASVFNNQVSLKSLNLQKNLITSVEKKVFGPAFRNLTELDMRFNPFDCTCESIAWFVNWINETHTNIPELSSHYLCNTPPHYHGFPVRLFDTSSCKDSAPFELFFMINTSILLIFIFIVLLIHFEGWRISFYWNVSVHRVLGFKEIDRQTEQFEYAAYIIHAYKDKDWVWEHFSSMEKEDQSLKFCLEERDFEAGVFELEAIVNSIKRSRKIIFVITHHLLKDPLCKRFKVHHAV | MRQTLPCIYFWGGLLPFGMLCASSTTKCTVSHEVADCSHLKLTQVPDDLPTNITVLNLTHNQLRRLPAANFTRYSQLTSLDVGFNTISKLEPELCQKLPMLKVLNLQHNELSQLSDKTFAFCTNLTELHLMSNSIQKIKNNPFVKQKNLITLDLSHNGLSSTKLGTQVQLENLQELLLSNNKIQALKSEELDIFANSSLKKLELSSNQIKEFSPGCFHAIGRLFGLFLNNVQLGPSLTEKLCLELANTSIRNLSLSNSQLSTTSNTTFLGLKWTNLTMLDLSYNNLNVVGNDSFAWLPQLEYFFLEYNNIQHLFSHSLHGLFNVRYLNLKRSFTKQSISLASLPKIDDFSFQWLKCLEHLNMEDNDIPGIKSNMFTGLINLKYLSLSNSFTSLRTLTNETFVSLAHSPLHILNLTKNKISKIESDAFSWLGHLEVLDLGLNEIGQELTGQEWRGLENIFEIYLSYNKYLQLTRNSFALVPSLQRLMLRRVALKNVDSSPSPFQPLRNLTILDLSNNNIANINDDMLEGLEKLEILDLQHNNLARLWKHANPGGPIYFLKGLSHLHILNLESNGFDEIPVEVFKDLFELKIIDLGLNNLNTLPASVFNNQVSLKSLNLQKNLITSVEKKVFGPAFRNLTELDMRFNPFDCTCESIAWFVNWINETHTNIPELSSHYLCNTPPHYHGFPVRLFDTSSCKDSAPFELFFMINTSILLIFIFIVLLIHFEGWRISFYWNVSVHRVLGFKEIDRQTEQFEYAAYIIHAYKDKDWVWEHFSSMEKEDQSLKFCLEERDFEAGVFELEAIVNSIKRSRKIIFVITHHLLKDPLCKRFKVHHAV |

| | | | | | | | | | | | | | | | | | |
|---|---|---|---|---|---|---|---|---|---|---|---|---|---|---|---|---|---|
| | | | | | | | | | | | | | | | | QQAIEQNLDSIILVFLEEIPDYKLNHALCLRRGMFKSHCILNWPVQKERIGAFRHKLQVALGSKNSVH | QQAIEQNLDSIILVFLEEIPDYKLNHALCLRRGMFKSHCILNWPVQKERIGAFRHKLQVALGSKNSVH |
| **2MKA** | Low (Bio/Crystallographic) | - | O15455 | TLR3 | O15455 | | A | C | 1 | T.{2}LL.{2}IF.{2}LL.{2}F | F.{3}VL.{2}HF | 8 | 5 | 16 | 10 | MRQTLPCIYFWGGLLPFGMLCASSTTKCTVSHEVADCSHLKLTQVPDDLPTNITVLNLTHNQLRRLPAANFTRYSQLTSLDVGFNTISKLEPELCQKLPMLKVLNLQHNELSQLSDKTFAFCTNLTELHLMSNSIQKIKNNPFVKQKNLITLDLSHNGLSSTKLGTQVQLENLQELLLSNNKIQALKSEELDIFANSSLKKLELSSNQIKEFSPGCFHAIGRLFGLFLNNVQLGPSLTEKLCLELANTSIRNLSLSNSQLSTTSNTTFLGLKWTNLTMLDLSYNNLNVVGNDSFAWLPQLEYFFLEYNNIQHLFSHSLHGLFNVRYLNLKRSFTKQSISLASLPKIDDFSFQWLKCLEHLNMEDNDIPGIKSNMFTGLINLKYLSLSNSFTSLRTLTNETFVSLAHSPLHILNLTKNKISKIESDAFSWLGHLEVLDLGLNEIGQELTGQEWRGLENIFEIYLSYNKYLQLTRNSFALVPSLQRLMLRRVALKNVDSSPSPFQPLRNLTILDLSNNNIANINDDMLEGLEKLEILDLQHNNLARLWKHANPGGPIYFLKGLSHLHILNLESNGFDEIPVEVFKDLFELKIIDLGLNNLNTLPASVFNNQVSLKSLNLQKNLITSVEKKVFGPAFRNLTELDMRFNPFDCTCESIAWFVNWINETHTNIPELSSHYLCNTPPHYHGFPVRLFDTSSCKDSAPFELFFMINTSILLIFIFIVLLIHFEGWRISFYWNVSVHRVLGFKEIDRQTEQFEYAAYIIHAYKDKDWVWEHFSSMEKEDQSLKFCLEERDFEAGVFELEAIVNSIKRSRKIIFVITHHLLKDPLCKRFKVHHAVQQAIEQNLDSIILVFLEEIPDYKLNHALCLRRGMFKSHCILNWPVQKERIGAFRHKLQVALGSKNSVH | MRQTLPCIYFWGGLLPFGMLCASSTTKCTVSHEVADCSHLKLTQVPDDLPTNITVLNLTHNQLRRLPAANFTRYSQLTSLDVGFNTISKLEPELCQKLPMLKVLNLQHNELSQLSDKTFAFCTNLTELHLMSNSIQKIKNNPFVKQKNLITLDLSHNGLSSTKLGTQVQLENLQELLLSNNKIQALKSEELDIFANSSLKKLELSSNQIKEFSPGCFHAIGRLFGLFLNNVQLGPSLTEKLCLELANTSIRNLSLSNSQLSTTSNTTFLGLKWTNLTMLDLSYNNLNVVGNDSFAWLPQLEYFFLEYNNIQHLFSHSLHGLFNVRYLNLKRSFTKQSISLASLPKIDDFSFQWLKCLEHLNMEDNDIPGIKSNMFTGLINLKYLSLSNSFTSLRTLTNETFVSLAHSPLHILNLTKNKISKIESDAFSWLGHLEVLDLGLNEIGQELTGQEWRGLENIFEIYLSYNKYLQLTRNSFALVPSLQRLMLRRVALKNVDSSPSPFQPLRNLTILDLSNNNIANINDDMLEGLEKLEILDLQHNNLARLWKHANPGGPIYFLKGLSHLHILNLESNGFDEIPVEVFKDLFELKIIDLGLNNLNTLPASVFNNQVSLKSLNLQKNLITSVEKKVFGPAFRNLTELDMRFNPFDCTCESIAWFVNWINETHTNIPELSSHYLCNTPPHYHGFPVRLFDTSSCKDSAPFELFFMINTSILLIFIFIVLLIHFEGWRISFYWNVSVHRVLGFKEIDRQTEQFEYAAYIIHAYKDKDWVWEHFSSMEKEDQSLKFCLEERDFEAGVFELEAIVNSIKRSRKIIFVITHHLLKDPLCKRFKVHHAVQQAIEQNLDSIILVFLEEIPDYKLNHALCLRRGMFKSHCILNWPVQKERIGAFRHKLQVALGSKNSVH |

| | | | | | | | | | | | | | | | | | |
|---|---|---|---|---|---|---|---|---|---|---|---|---|---|---|---|---|---|
| **2MKA (NMR)** | Low (Bio/Crystallographic) | - | O15455 | TLR3 | O15455 | | A | B | 1 | F.{3}N.{2}IL.{2}FI.{2}V.{3}H | F.{3}N.{3}L.{2}FI.{6}HF | 8 | 7 | 20 | 21 | MRQTLPCIYFWGGLLPFG<br>MLCASSTTKCTVSHEVAD<br>CSHLKLTQVPDDLPTNITV<br>LNLTHNQLRRLPAANFTR<br>YSQLTSLDVGFNTISKLEP<br>ELCQKLPMLKVLNLQHNE<br>LSQLSDKTFAFCTNLTELH<br>LMSNSIQKIKNNPFVKQK<br>NLITLDLSHNGLSSTKLGT<br>QVQLENLQELLLSNNKIQ<br>ALKSEELDIFANSSLKKLEL<br>SSNQIKEFSPGCFHAIGRL<br>FGLFLNNVQLGPSLTEKLC<br>LELANTSIRNLSLSNSQLST<br>TSNTTFLGLKWTNLTMLD<br>LSYNNLNVVGNDSFAWL<br>PQLEYFFLEYNNIQHLFSH<br>SLHGLFNVRYLNLKRSFTK<br>QSISLASLPKIDDFSFQWL<br>KCLEHLNMEDNDIPGIKS<br>NMFTGLINLKYLSLSNSFT<br>SLRTLTNETFVSLAHSPLHI<br>LNLTKNKISKIESDAFSWL<br>GHLEVLDLGLNEIGQELT<br>GQEWRGLENIFEIYLSYNK<br>YLQLTRNSFALVPSLQRL<br>MLRRVALKNVDSSPSPFQ<br>PLRNLTILDLSNNNIANIN<br>DDMLEGLEKLEILDLQHN<br>NLARLWKHANPGGPIYFL<br>KGLSHLHILNLESNGFDEI<br>PVEVFKDLFELKIIDLGLN<br>NLNTLPASVFNNQVSLKS<br>LNLQKNLITSVEKKVFGPA<br>FRNLTELDMRFNPFDCTC<br>ESIAWFVNWINETHTNIP<br>ELSSHYLCNTPPHYHGFP<br>VRLFDTSSCKDSAPFELFF<br>MINTSILLIFIFIVLLIHFEG<br>WRISFYWNVSVHRVLGF<br>KEIDRQTEQFEYAAYIIHA<br>YKDKDWVWEHFSSMEK<br>EDQSLKFCLEERDFEAGV<br>FELEAIVNSIKRSRKIIFVIT<br>HHLLKDPLCKRFKVHHAV<br>QQAIEQNLDSIILVFLEEIP<br>DYKLNHALCLRRGMFKS<br>HCILNWPVQKERIGAFRH<br>KLQVALGSKNSVH | MRQTLPCIYFWGGLLPFG<br>MLCASSTTKCTVSHEVAD<br>CSHLKLTQVPDDLPTNITV<br>LNLTHNQLRRLPAANFTR<br>YSQLTSLDVGFNTISKLEP<br>ELCQKLPMLKVLNLQHNE<br>LSQLSDKTFAFCTNLTELH<br>LMSNSIQKIKNNPFVKQK<br>NLITLDLSHNGLSSTKLGT<br>QVQLENLQELLLSNNKIQ<br>ALKSEELDIFANSSLKKLEL<br>SSNQIKEFSPGCFHAIGRL<br>FGLFLNNVQLGPSLTEKLC<br>LELANTSIRNLSLSNSQLST<br>TSNTTFLGLKWTNLTMLD<br>LSYNNLNVVGNDSFAWL<br>PQLEYFFLEYNNIQHLFSH<br>SLHGLFNVRYLNLKRSFTK<br>QSISLASLPKIDDFSFQWL<br>KCLEHLNMEDNDIPGIKS<br>NMFTGLINLKYLSLSNSFT<br>SLRTLTNETFVSLAHSPLHI<br>LNLTKNKISKIESDAFSWL<br>GHLEVLDLGLNEIGQELT<br>GQEWRGLENIFEIYLSYNK<br>YLQLTRNSFALVPSLQRL<br>MLRRVALKNVDSSPSPFQ<br>PLRNLTILDLSNNNIANIN<br>DDMLEGLEKLEILDLQHN<br>NLARLWKHANPGGPIYFL<br>KGLSHLHILNLESNGFDEI<br>PVEVFKDLFELKIIDLGLN<br>NLNTLPASVFNNQVSLKS<br>LNLQKNLITSVEKKVFGPA<br>FRNLTELDMRFNPFDCTC<br>ESIAWFVNWINETHTNIP<br>ELSSHYLCNTPPHYHGFP<br>VRLFDTSSCKDSAPFELFF<br>MINTSILLIFIFIVLLIHFEG<br>WRISFYWNVSVHRVLGF<br>KEIDRQTEQFEYAAYIIHA<br>YKDKDWVWEHFSSMEK<br>EDQSLKFCLEERDFEAGV<br>FELEAIVNSIKRSRKIIFVIT<br>HHLLKDPLCKRFKVHHAV<br>QQAIEQNLDSIILVFLEEIP<br>DYKLNHALCLRRGMFKS<br>HCILNWPVQKERIGAFRH<br>KLQVALGSKNSVH |
| **2N2A** | Low | - | P046 | ERB | P046 | | A | B | 1 | I.{3}V.{3}L.{2}VL.{2}V.{3}L | I.{3}V.{3}L.{2}VL.{3}F.{2}L | 7 | 7 | 20 | 20 | MELAALCRWGLLLALLPP<br>GAASTQVCTGTDMKLRL<br>PASPETHLDMLRHLYQG<br>CQVVQGNLELTYLPTNAS | MELAALCRWGLLLALLPP<br>GAASTQVCTGTDMKLRL<br>PASPETHLDMLRHLYQG<br>CQVVQGNLELTYLPTNAS |

| | | | | | | | | | | | | | | | | | |
|---|---|---|---|---|---|---|---|---|---|---|---|---|---|---|---|---|---|
| **(NMR)** | (Bio/Crystallographic) | | 26 | B2 | 26 | | | | | | | | | | | LSFLQDIQEVQGYVLIAH<br>NQVRQVPLQRLRIVRGT<br>QLFEDNYALAVLDNGDPL<br>NNTTPVTGASPGGLRELQ<br>LRSLTEILKGGVLIQRNPQ<br>LCYQDTILWKDIFHKNNQ<br>LALTLIDTNRSRACHPCSP<br>MCKGSRCWGESSEDCQS<br>LTRTVCAGGCARCKGPLP<br>TDCCHEQCAAGCTGPKH<br>SDCLACLHFNHSGICELHC<br>PALVTYNTDTFESMPNPE<br>GRYTFGASCVTACPYNYL<br>STDVGSCTLVCPLHNQEV<br>TAEDGTQRCEKCSKPCAR<br>VCYGLGMEHLREVRAVT<br>SANIQEFAGCKKIFGSLAF<br>LPESFDGDPASNTAPLQP<br>EQLQVFETLEEITGYLYISA<br>WPDSLPDLSVFQNLQVIR<br>GRILHNGAYSLTLQGLGIS<br>WLGLRSLRELGSGLALIHH<br>NTHLCFVHTVPWDQLFR<br>NPHQALLHTANRPEDEC<br>VGEGLACHQLCARGHC<br>WGPGPTQCVNCSQFLRG<br>QECVEECRVLQGLPREYV<br>NARHCLPCHPECQPQNG<br>SVTCFGPEADQCVACAH<br>YKDPPFCVARCPSGVKPD<br>LSYMPIWKFPDEEGACQ<br>PCPINCTHSCVDLDDKGC<br>PAEQRASPLTSIISAVVGIL<br>LVVVLGVVFGILIKRRQQK<br>IRKYTMRRLLQETELVEPL<br>TPSGAMPNQAQMRILKE<br>TELRKVKVLGSGAFGTVY<br>KGIWIPDGENVKIPVAIKV<br>LRENTSPKANKEILDEAYV<br>MAGVGSPYVSRLLGICLT<br>STVQLVTQLMPYGCLLDH<br>VRENRGRLGSQDLLNWC<br>MQIAKGMSYLEDVRLVH<br>RDLAARNVLVKSPNHVKI<br>TDFGLARLLDIDETEYHAD<br>GGKVPIKWMALESILRRR<br>FTHQSDVWSYGVTVWEL<br>MTFGAKPYDGIPAREIPD<br>LLEKGERLPQPPICTIDVY<br>MIMVKCWMIDSECRPRF<br>RELVSEFSRMARDPQRFV<br>VIQNEDLGPASPLDSTFYR<br>SLLEDDDMGDLVDAEEYL | LSFLQDIQEVQGYVLIAH<br>NQVRQVPLQRLRIVRGT<br>QLFEDNYALAVLDNGDPL<br>NNTTPVTGASPGGLRELQ<br>LRSLTEILKGGVLIQRNPQ<br>LCYQDTILWKDIFHKNNQ<br>LALTLIDTNRSRACHPCSP<br>MCKGSRCWGESSEDCQS<br>LTRTVCAGGCARCKGPLP<br>TDCCHEQCAAGCTGPKH<br>SDCLACLHFNHSGICELHC<br>PALVTYNTDTFESMPNPE<br>GRYTFGASCVTACPYNYL<br>STDVGSCTLVCPLHNQEV<br>TAEDGTQRCEKCSKPCAR<br>VCYGLGMEHLREVRAVT<br>SANIQEFAGCKKIFGSLAF<br>LPESFDGDPASNTAPLQP<br>EQLQVFETLEEITGYLYISA<br>WPDSLPDLSVFQNLQVIR<br>GRILHNGAYSLTLQGLGIS<br>WLGLRSLRELGSGLALIHH<br>NTHLCFVHTVPWDQLFR<br>NPHQALLHTANRPEDEC<br>VGEGLACHQLCARGHC<br>WGPGPTQCVNCSQFLRG<br>QECVEECRVLQGLPREYV<br>NARHCLPCHPECQPQNG<br>SVTCFGPEADQCVACAH<br>YKDPPFCVARCPSGVKPD<br>LSYMPIWKFPDEEGACQ<br>PCPINCTHSCVDLDDKGC<br>PAEQRASPLTSIISAVVGIL<br>LVVVLGVVFGILIKRRQQK<br>IRKYTMRRLLQETELVEPL<br>TPSGAMPNQAQMRILKE<br>TELRKVKVLGSGAFGTVY<br>KGIWIPDGENVKIPVAIKV<br>LRENTSPKANKEILDEAYV<br>MAGVGSPYVSRLLGICLT<br>STVQLVTQLMPYGCLLDH<br>VRENRGRLGSQDLLNWC<br>MQIAKGMSYLEDVRLVH<br>RDLAARNVLVKSPNHVKI<br>TDFGLARLLDIDETEYHAD<br>GGKVPIKWMALESILRRR<br>FTHQSDVWSYGVTVWEL<br>MTFGAKPYDGIPAREIPD<br>LLEKGERLPQPPICTIDVY<br>MIMVKCWMIDSECRPRF<br>RELVSEFSRMARDPQRFV<br>VIQNEDLGPASPLDSTFYR<br>SLLEDDDMGDLVDAEEYL |

| | | | | | | | | | | | | | | | | | |
|---|---|---|---|---|---|---|---|---|---|---|---|---|---|---|---|---|---|
| | | | | | | | | | | | | | | | | VPQQGFFCPDPAPGAGGMVHHRHRSSSTRSGGGDLTLGLEPSEEEAPRSPLAPSEGAGSDVFDGDLGMGAAKGLQSLPTHDPSPLQRYSEDPTVPLPSETDGYVAPLTCSPQPEYVNQPDVRPQPPSPREGPLPAARPAGATLERPKTLSPGKNGVVKDVFAFGGAVENPEYLTPQGGAAPQPHPPPAFSPAFDNLYYWDQDPPERGAPPSTFKGTPTAENPEYLGLDVPV | VPQQGFFCPDPAPGAGGMVHHRHRSSSTRSGGGDLTLGLEPSEEEAPRSPLAPSEGAGSDVFDGDLGMGAAKGLQSLPTHDPSPLQRYSEDPTVPLPSETDGYVAPLTCSPQPEYVNQPDVRPQPPSPREGPLPAARPAGATLERPKTLSPGKNGVVKDVFAFGGAVENPEYLTPQGGAAPQPHPPPAFSPAFDNLYYWDQDPPERGAPPSTFKGTPTAENPEYLGLDVPV |
| **2N90 (NMR)** | Low (Bio/Crystallographic) | - | P04629 | NTRK1 | P04629 | | A | B | 1 | LA.FA.LFL | LA.FA.LFL.{5}V | 7 | 8 | 9 | 15 | MLRGGRRGQLGWHSWAAGPGSLLAWLILASAGAAPCPDACCPHGSSGLRCTRDGALDSLHHLPGAENLTELYIENQQHLQHLELRDLRGLGELRNLTIVKSGLRFVAPDAFHFTPRLSRLNLSFNALESLSWKTVQGLSLQELVLSGNPLHCSCALRWLQRWEEEGLGGVPEQKLQCHGQGPLAHMPNASCGVPTLKVQVPNASVDVGDDVLLRCQVEGRGLEQAGWILTELEQSATVMKSGGLPSLGLTLANVTSDLNRKNVTCWAENDVGRAEVSVQVNVSFPASVQLHTAVEMHHWCIPFSVDGQPAPSLRWLFNGSVLNETSFIFTEFLEPAANETVRHGCLRLNQPTHVNNGNYTLLAANPFGQASASIMAAFMDNPFEFNPEDPIPVSFSPVDTNSTSGDPVEKKDETPFGVSVAVGLAVFACLFLSTLLLVLNKCGRRNKFGINRPAVLAPEDGLAMSLHFMTLGGSSLSPTEGKGSGLQGHIIENPQYFSDACVHHIKRRDIVLKWELGEGAFGKVFLAECHNLLPEQDKMLVAVKALKEASESARQDFQREAELLTMLQHQHIVRFFGVCTEGRPLLMVFEYMRHGDLNRFLRSHGPDAKLLAGGEDVAPGPLGLGQLLAVASQVAAGMVYLAGLHFVHRDLATRNCLVGQGLVVKIGDFGMSRDIYSTDYYRVGGRTML | MLRGGRRGQLGWHSWAAGPGSLLAWLILASAGAAPCPDACCPHGSSGLRCTRDGALDSLHHLPGAENLTELYIENQQHLQHLELRDLRGLGELRNLTIVKSGLRFVAPDAFHFTPRLSRLNLSFNALESLSWKTVQGLSLQELVLSGNPLHCSCALRWLQRWEEEGLGGVPEQKLQCHGQGPLAHMPNASCGVPTLKVQVPNASVDVGDDVLLRCQVEGRGLEQAGWILTELEQSATVMKSGGLPSLGLTLANVTSDLNRKNVTCWAENDVGRAEVSVQVNVSFPASVQLHTAVEMHHWCIPFSVDGQPAPSLRWLFNGSVLNETSFIFTEFLEPAANETVRHGCLRLNQPTHVNNGNYTLLAANPFGQASASIMAAFMDNPFEFNPEDPIPVSFSPVDTNSTSGDPVEKKDETPFGVSVAVGLAVFACLFLSTLLLVLNKCGRRNKFGINRPAVLAPEDGLAMSLHFMTLGGSSLSPTEGKGSGLQGHIIENPQYFSDACVHHIKRRDIVLKWELGEGAFGKVFLAECHNLLPEQDKMLVAVKALKEASESARQDFQREAELLTMLQHQHIVRFFGVCTEGRPLLMVFEYMRHGDLNRFLRSHGPDAKLLAGGEDVAPGPLGLGQLLAVASQVAAGMVYLAGLHFVHRDLATRNCLVGQGLVVKIGDFGMSRDIYSTDYYRVGGRTML |

| | | | | | | | | | | | | | | | | | |
|---|---|---|---|---|---|---|---|---|---|---|---|---|---|---|---|---|---|
| | | | | | | | | | | | | | | | | PIRWMPPESILYRKFTTESDVWSFGVVLWEIFTYGKQPWYQLSNTEAIDCITQGRELERPRACPPEVYAIMRGCWQREPQQRHSIKDVHARLQALAQAPPVYLDVLG | PIRWMPPESILYRKFTTESDVWSFGVVLWEIFTYGKQPWYQLSNTEAIDCITQGRELERPRACPPEVYAIMRGCWQREPQQRHSIKDVHARLQALAQAPPVYLDVLG |
| **2NA6 (NMR)** | Low (Bio/Crystallographic) | - | P25446 | Fas | P25446 | | A | C | 1 | T.LV.{2}I.{2}V | T.{2}VL.IP.{2}F | 5 | 6 | 10 | 11 | MLWIWAVLPLVLAGSQLRVHTQGTNSISESLKLRRRVRETDKNCSEGLYQGGPFCCQPCQPGKKKVEDCKMNGGTPTCAPCTEGKEYMDKNHYADKCRRCTLCDEEHGLEVETNCTLTQNTKCKCKPDFYCDSPGCEHCVRCASCEHGTLEPCTATSNTNCRKQSPRNRLWLLTILVLLIPLVFIYRKYRKRKCWKRRQDDPESRTSSRETIPMNASNLSLSKYIPRIAEDMTIQEAKKFARENNIKEGKIDEIMHDSIQDTAEQKVQLLLCWYQSHGKSDAYQDLIKGLKKAECRRTLDKFQDMVQKDLGKSTPDTGNENEGQCLE | MLWIWAVLPLVLAGSQLRVHTQGTNSISESLKLRRRVRETDKNCSEGLYQGGPFCCQPCQPGKKKVEDCKMNGGTPTCAPCTEGKEYMDKNHYADKCRRCTLCDEEHGLEVETNCTLTQNTKCKCKPDFYCDSPGCEHCVRCASCEHGTLEPCTATSNTNCRKQSPRNRLWLLTILVLLIPLVFIYRKYRKRKCWKRRQDDPESRTSSRETIPMNASNLSLSKYIPRIAEDMTIQEAKKFARENNIKEGKIDEIMHDSIQDTAEQKVQLLLCWYQSHGKSDAYQDLIKGLKKAECRRTLDKFQDMVQKDLGKSTPDTGNENEGQCLE |
| **2QTS (X-ray)** | High (Bio) | 3IJ4 or 4FZ1 or 4NTW or 4NTX | Q1XA76 | ASIC1 | Q1XA76 | | A | B | 2 | G.M.{3}I.{3}I.{3}L.{2}F | K.{3}W.{2}CF.{5}L.{379}Q | 6 | 6 | 18 | 395 | MMDLKVDEEEVDSGQPVSIQAFASSSTLHGISHIFSYERLSLKRVVWALCFMGSLALLALVCTNRIQYYFLYPHVTKLDEVAATRLTFPAVTFCNLNEFRFSRVTKNDLYHAGELLALLNNRYEIPDTQTADEKQLEILQDKANFRNFKPKPFNMLEFYDRAGHDIREMLLSCFFRGEQCSPEDFKVVFTRYGKCYTFNAGQDGKPRLITMKGGTGNGLEIMLDIQQDEYLPVWGETDETSFEAGIKVQIHSQDEPPLIDQLGFGVAPGFQTFVSCQEQRLIYLPPPWGDCKATTGDSEFYDTYSITACRIDCETRYLVENCNCRMVHMPGDAPYCTPEQYKECADPALDFLVEKDNEYCVCEMPCNVTRYGKELSMVKIPSKASAKYLAKKY | MMDLKVDEEEVDSGQPVSIQAFASSSTLHGISHIFSYERLSLKRVVWALCFMGSLALLALVCTNRIQYYFLYPHVTKLDEVAATRLTFPAVTFCNLNEFRFSRVTKNDLYHAGELLALLNNRYEIPDTQTADEKQLEILQDKANFRNFKPKPFNMLEFYDRAGHDIREMLLSCFFRGEQCSPEDFKVVFTRYGKCYTFNAGQDGKPRLITMKGGTGNGLEIMLDIQQDEYLPVWGETDETSFEAGIKVQIHSQDEPPLIDQLGFGVAPGFQTFVSCQEQRLIYLPPPWGDCKATTGDSEFYDTYSITACRIDCETRYLVENCNCRMVHMPGDAPYCTPEQYKECADPALDFLVEKDNEYCVCEMPCNVTRYGKELSMVKIPSKASAKYLAKKY |

| | | | | | | | | | | | | | | | | | |
|---|---|---|---|---|---|---|---|---|---|---|---|---|---|---|---|---|---|
| | | or4NTYor4NYK | | | | | | | | | | | | | | NKSEQYIGENILVLDIFFEALNYETIEQKKAYEVAGLLGDIGGQMGLFIGASILTVLELFDYAYEVIKHRLCRRGKCRKNHKRNNTDKGVALSMDDVKRHNPCESLRGHPAGMTYAANILPHHPARGTFEDFTC | NKSEQYIGENILVLDIFFEALNYETIEQKKAYEVAGLLGDIGGQMGLFIGASILTVLELFDYAYEVIKHRLCRRGKCRKNHKRNNTDKGVALSMDDVKRHNPCESLRGHPAGMTYAANILPHHPARGTFEDFTC |
| **2ZW3 (X-ray)** | High (Bio) | 5ER7or5ERA | P29033 | GJB2 | P29033 | | A | B | 4 | W.{2}V.{3}F.{2}M.{3}V.{3}EV.{143}E.{2}VF.{2}F | IR.{3}L.{2}IF.{2}T.{3}L | 11 | 7 | 171 | 17 | MDWGTLQTILGGVNKHSTSIGKIWLTVLFIFRIMILVVAAKEVWGDEQADFVCNTLQPGCKNVCYDHYFPISHIRLWALQLIFVSTPALLVAMHVAYRRHEKKRKFIKGEIKSEFKDIEEIKTQKVRIEGSLWWTYTSSIFFRVIFEAAFMYVFYVMYDGFSMQRLVKCNAWPCPNTVDCFVSRPTEKTVFTVFMIAVSGICILLNVTELCYLLIRYCSGKSKKPV | MDWGTLQTILGGVNKHSTSIGKIWLTVLFIFRIMILVVAAKEVWGDEQADFVCNTLQPGCKNVCYDHYFPISHIRLWALQLIFVSTPALLVAMHVAYRRHEKKRKFIKGEIKSEFKDIEEIKTQKVRIEGSLWWTYTSSIFFRVIFEAAFMYVFYVMYDGFSMQRLVKCNAWPCPNTVDCFVSRPTEKTVFTVFMIAVSGICILLNVTELCYLLIRYCSGKSKKPV |
| **3D9S (X-ray)** | High (Bio) | - | P55064 | AQP5 | P55064 | | B | D | 6 | K.{6}L.{6}FF.{2}GS.{11}LQ.{2}LA.{2}LA.{2}T.{2}QAL.PV.{40}I.{58}L | L.{86}Q.{3}V.{2}IL.{2}Q.{3}C.{2}AS.{14}S.{2}LS.TL.{2}LV.{44}L.{3}L | 20 | 18 | 152 | 182 | MKKEVCSVAFLKAVFAEFLATLIFVFFGLGSALKWPSALPTILQIALAFGLAIGTLAQALGPVSGGHINPAITLALLVGNQISLLRAFFYVAAQLVGAIAGAGILYGVAPLNARGNLAVNALNNNTTQGQAMVVELILTFQLALCIFASTDSRRTSPVGSPALSIGLSVTLGHLVGIYFTGCSMNPARSFGPAVVMNRFSPAHWVFWVGPIVGAVLAAILYFYLLFPNSLSLSERVAIIKGTYEPDEDWEEQREERKKTMELTTR | MKKEVCSVAFLKAVFAEFLATLIFVFFGLGSALKWPSALPTILQIALAFGLAIGTLAQALGPVSGGHINPAITLALLVGNQISLLRAFFYVAAQLVGAIAGAGILYGVAPLNARGNLAVNALNNNTTQGQAMVVELILTFQLALCIFASTDSRRTSPVGSPALSIGLSVTLGHLVGIYFTGCSMNPARSFGPAVVMNRFSPAHWVFWVGPIVGAVLAAILYFYLLFPNSLSLSERVAIIKGTYEPDEDWEEQREERKKTMELTTR |
| **3J5P (EM)** | High (-) | - | O35433 | Trpv1 | O35433 | | A | B | 6 | R.{2}FV.{2}VF.{2}G.{2}TA.VT.{58}K.V.I.L.{5}I.{2}YIL.{2}NM.IAL | T.{2}AY.{85}A.{2}VF.L.{3}WT.{2}L.{18}MI.{3}L.{97}L.{2}LI.{2}M | 23 | 17 | 103 | 234 | MEQRASLDSEESESPPQENSCLDPPDRDPNCKPPPVKPHIFTTRSRTRLFGKGDSEEASPLDCPYEEGGLASCPIITVSSVLTIQRPGDGPASVRPSSQDSVSAGEKPPRLYDRRSIFDAVAQSNCQELESLLPFLQRSKKRLTDSEFKDPETGKTCLLKAMLNLHNGQNDTIALLLDVARKTDSLKQFVNASYTDSYYKGQTALHIAIERRNMTLVTLLVENGADVQAAANGDFFKK | MEQRASLDSEESESPPQENSCLDPPDRDPNCKPPPVKPHIFTTRSRTRLFGKGDSEEASPLDCPYEEGGLASCPIITVSSVLTIQRPGDGPASVRPSSQDSVSAGEKPPRLYDRRSIFDAVAQSNCQELESLLPFLQRSKKRLTDSEFKDPETGKTCLLKAMLNLHNGQNDTIALLLDVARKTDSLKQFVNASYTDSYYKGQTALHIAIERRNMTLVTLLVENGADVQAAANGDFFKK |

| | | | | | | | | | | | | | | | | | |
|---|---|---|---|---|---|---|---|---|---|---|---|---|---|---|---|---|---|
| | | | | | | | | | | | | | | | | TKGRPGFYFGELPLSLAACTNQLAIVKFLLQNSWQPADISARDSVGNTVLHALVEVADNTVDNTKFVTSMYNEILILGAKLHPTLKLEEITNRKGLTPLALAASSGKIGVLAYILQREIHEPECRHLSRKFTEWAYGPVHSSLYDLSCIDTCEKNSVLEVIAYSSSETPNRHDMLLVEPLNRLLQDKWDRFVKRIFYFNFFVYCLYMIIFTAAAYYRPVEGLPPYKLKNTVGDYFRVTGEILSVSGGVYFFFRGIQYFLQRRPSLKSLFVDSYSEILFFVQSLFMLVSVVLYFSQRKEYVASMVFSLAMGWTNMLYYTRGFQQMGIYAVMIEKMILRDLCRFMFVYLVFLFGFSTAVVTLIEDGKNNSLPMESTPHKCRGSACKPGNSYNSLYSTCLELFKFTIGMGDLEFTENYDFKAVFIILLLAYVILTYILLLNMLIALMGETVNKIAQESKNIWKLQRAITILDTEKSFLKCMRKAFRSGKLLQVGFTPDGKDDYRWCFRVDEVNWTTWNTNVGIINEDPGNCEGVKRTLSFSLRSGRVSGRNWKNFALVPLLRDASTRDRHATQQEEVQLKHYTGSLKPEDAEVFKDSMVPGEK | TKGRPGFYFGELPLSLAACTNQLAIVKFLLQNSWQPADISARDSVGNTVLHALVEVADNTVDNTKFVTSMYNEILILGAKLHPTLKLEEITNRKGLTPLALAASSGKIGVLAYILQREIHEPECRHLSRKFTEWAYGPVHSSLYDLSCIDTCEKNSVLEVIAYSSSETPNRHDMLLVEPLNRLLQDKWDRFVKRIFYFNFFVYCLYMIIFTAAAYYRPVEGLPPYKLKNTVGDYFRVTGEILSVSGGVYFFFRGIQYFLQRRPSLKSLFVDSYSEILFFVQSLFMLVSVVLYFSQRKEYVASMVFSLAMGWTNMLYYTRGFQQMGIYAVMIEKMILRDLCRFMFVYLVFLFGFSTAVVTLIEDGKNNSLPMESTPHKCRGSACKPGNSYNSLYSTCLELFKFTIGMGDLEFTENYDFKAVFIILLLAYVILTYILLLNMLIALMGETVNKIAQESKNIWKLQRAITILDTEKSFLKCMRKAFRSGKLLQVGFTPDGKDDYRWCFRVDEVNWTTWNTNVGIINEDPGNCEGVKRTLSFSLRSGRVSGRNWKNFALVPLLRDASTRDRHATQQEEVQLKHYTGSLKPEDAEVFKDSMVPGEK |
| **3SYQ (X-ray)** | High (Bio) | 3SYAor3SYCor3SYOor3S | P48542 | Kcnj6 | P48542 | | A | B | 2 | L.{3}V.{3}T.{2}W.{81}V.{2}MF | LL.{3}VL.SI.{2}AF.{2}G | 7 | 9 | 98 | 17 | MTMAKLTESMTNVLEGDSMDQDVESPVAIHQPKLPKQARDDLPRHISRDRTKRKIQRYVRKDGKCNVHHGNVRETYRYLTDIFTTLVDLKWRFNLLIFVMVYTVTWLFFGMIWWLIAYIRGDMDHIEDPSWTPCVTNLNGFVSAFLFSIETETTIGYGYRVITDKCPEGIILLLIQSVLGSIVNAFMVGCMFVKISQPKKRAETLVFSTHAVISMRDGKLCLMFRVGDLRNSHIVEASIRAKLIKSKQTSEGEFIPLNQSDINVGYYTGDDRLFLVSPLIISHEINQQSPFWEISKAQLPKEELEIVVILEGIVEATGMTCQARSSYITSEILWGYRFTPVLTMEDGFYEVDYNSFHETYETSTP | MTMAKLTESMTNVLEGDSMDQDVESPVAIHQPKLPKQARDDLPRHISRDRTKRKIQRYVRKDGKCNVHHGNVRETYRYLTDIFTTLVDLKWRFNLLIFVMVYTVTWLFFGMIWWLIAYIRGDMDHIEDPSWTPCVTNLNGFVSAFLFSIETETTIGYGYRVITDKCPEGIILLLIQSVLGSIVNAFMVGCMFVKISQPKKRAETLVFSTHAVISMRDGKLCLMFRVGDLRNSHIVEASIRAKLIKSKQTSEGEFIPLNQSDINVGYYTGDDRLFLVSPLIISHEINQQSPFWEISKAQLPKEELEIVVILEGIVEATGMTCQARSSYITSEILWGYRFTPVLTMEDGFYEVDYNSFHETYETSTP |

| | | | | | | | | | | | | | | | | | |
|---|---|---|---|---|---|---|---|---|---|---|---|---|---|---|---|---|---|
| | | YPor4KFM | | | | | | | | | | | | | | SLSAKELAELANRAEVPLSWSVSSKLNQHAELETEEEEKNPEELTERNGDVANLENESKV | SLSAKELAELANRAEVPLSWSVSSKLNQHAELETEEEEKNPEELTERNGDVANLENESKV |
| **4COF (X-ray)** | High (Bio) | - | P28472 | GABRB3 | P28472 | | A | B | 4 | LQ.{6}LI.IL.{2}V.{2}WI.{6}A.{2}AL.{2}T.{10}H.{160}R | S.{3}V.{3}I.{3}L.{6}T.{19}M.{2}F.{3}F.{2}LL.{5}N | 15 | 11 | 206 | 57 | MWGLAGGRLFGIFSAPVLVAVVCCAQSVNDPGNMSFVKETVDKLLKGYDIRLRPDFGGPPVCVGMNIDIASIDMVSEVNMDYTLTMYFQQYWRDKRLAYSGIPLNLTLDNRVADQLWVPDTYFLNDKKSFVHGVTVKNRMIRLHPDGTVLYGLRITTTAACMMDLRRYPLDEQNCTLEIESYGYTTDDIEFYWRGGDKAVTGVERIELPQFSIVEHRLVSRNVVFATGAYPRLSLSFRLKRNIGYFILQTYMPSILITILSWVSFWINYDASAARVALGITTVLTMTTINTHLRETLPKIPYVKAIDMYLMGCFVFVFLALLEYAFVNYIFFGRGPQRQKKLAEKTAKAKNDRSKSESNRVDAHGNILLTSLEVHNEMNEVSGGIGDTRNSAISFDNSGIQYRKQSMPREGHGRFLGDRSLPHKKTHLRRRSSQLKIKIPDLTDVNAIDRWSRIVFPFTFSLFNLVYWLYYVN | MWGLAGGRLFGIFSAPVLVAVVCCAQSVNDPGNMSFVKETVDKLLKGYDIRLRPDFGGPPVCVGMNIDIASIDMVSEVNMDYTLTMYFQQYWRDKRLAYSGIPLNLTLDNRVADQLWVPDTYFLNDKKSFVHGVTVKNRMIRLHPDGTVLYGLRITTTAACMMDLRRYPLDEQNCTLEIESYGYTTDDIEFYWRGGDKAVTGVERIELPQFSIVEHRLVSRNVVFATGAYPRLSLSFRLKRNIGYFILQTYMPSILITILSWVSFWINYDASAARVALGITTVLTMTTINTHLRETLPKIPYVKAIDMYLMGCFVFVFLALLEYAFVNYIFFGRGPQRQKKLAEKTAKAKNDRSKSESNRVDAHGNILLTSLEVHNEMNEVSGGIGDTRNSAISFDNSGIQYRKQSMPREGHGRFLGDRSLPHKKTHLRRRSSQLKIKIPDLTDVNAIDRWSRIVFPFTFSLFNLVYWLYYVN |
| **4EZC (X-ray)** | Low (Bio) | - | Q5QF96 | SLC14A1 | Q5QF96 | | A | B | 8 | L.{45}T.{3}F.SA.{2}SV.{18}M.{2}SAT.{140}V | LSL.{43}F.{2}YL.AS.{2}HV.{2}V.{15}L.{2}L | 12 | 13 | 223 | 80 | MDDNPTAVKLDQGGNQAPQGRGRRCLPKALGYITGDMKEFANWLKDKPQALQFVDWVLRGISQVVFVSNPISGILILVGLLVQNPWCALNGCVGTVVSTLTALLLSQDRSAITAGLQGYNATLVGILMAIYSDKGNYFWWLLFPVSAMSMTCPVFSSALNSVLSKWDLPVFTLPFNMALSMYLSATGHYNPFFPSTLITPVTSVPNVTWPDLSALQLLKSLPVGVGQIYGCDNPWTGGIFLGAILLSSPLMCLHAAIGSLLGIIAGLSLSAPFEDIYAGLWGFNSSLACIAIGGTFMALTWQTHLLALACALFTAYLGASMSHV | MDDNPTAVKLDQGGNQAPQGRGRRCLPKALGYITGDMKEFANWLKDKPQALQFVDWVLRGISQVVFVSNPISGILILVGLLVQNPWCALNGCVGTVVSTLTALLLSQDRSAITAGLQGYNATLVGILMAIYSDKGNYFWWLLFPVSAMSMTCPVFSSALNSVLSKWDLPVFTLPFNMALSMYLSATGHYNPFFPSTLITPVTSVPNVTWPDLSALQLLKSLPVGVGQIYGCDNPWTGGIFLGAILLSSPLMCLHAAIGSLLGIIAGLSLSAPFEDIYAGLWGFNSSLACIAIGGTFMALTWQTHLLALACALFTAYLGASMSHV |

| | | | | | | | | | | | | | | | | | |
|---|---|---|---|---|---|---|---|---|---|---|---|---|---|---|---|---|---|
| | | | | | | | | | | | | | | | | MAVVGLPSGTWPFCLATLLFLLLTTKNPNIYKMPISKVTYPEENRIFYLQSRKRTVQGPL | MAVVGLPSGTWPFCLATLLFLLLTTKNPNIYKMPISKVTYPEENRIFYLQSRKRTVQGPL |
| **4JKV (X-ray)** | Low (Bio) | 4N4W or 4O9R or 4QIM or 4QIN | Q99835 | SMO | Q99835 | | A | B | 7 (GPCR) | Y.{2}H.{19}F.LL.{6}VL.VA.{2}A.{24}GF.{2}AP.{6}V.{2}YF.{2}R | V.{3}Y.{24}LL.{6}VL.VA.{2}A.{24}GF.{2}A.{11}F | 18 | 13 | 85 | 86 | MAAARPARGPELPLLGLLLLLLLGDPGRGAASSGNATGPGPRSAGGSARRSAAVTGPPPPLSHCGRAAPCEPLRYNVCLGSVLPYGATSTLLAGDSDSQEEAHGKLVLWSGLRNAPRCWAVIQPLLCAVYMPKCENDRVELPSRTLCQATRGPCAIVERERGWPDFLRCTPDRFPEGCTNEVQNIKFNSSGQCEVPLVRTDNPKSWYEDVEGCGIQCQNPLFTEAEHQDMHSYIAAFGAVTGLCTLFTLATFVADWRNSNRYPAVILFYVNACFFVGSIGWLAQFMDGARREIVCRADGTMRLGEPTSNETLSCVIIFVIVYYALMAGVVWFVVLTYAWHTSFKALGTTYQPLSGKTSYFHLLTWSLPFVLTVAILAVAQVDGDSVSGICFVGYKNYRYRAGFVLAPIGLVLIVGGYFLIRGVMTLFSIKSNHPGLLSEKAASKINETMLRLGIFGFLAFGFVLITFSCHFYDFFNQAEWERSFRDYVLCQANVTIGLPTKQPIPDCEIKNRPSLLVEKINLFAMFGTGIAMSTWVWTKATLLIWRRTWCRLTGQSDDEPKRIKKSKMIAKAFSKRHELLQNPGQELSFSMHTVSHDGPVAGLAFDLNEPSADVSSAWAQHVTKMVARRGAILPQDISVTPVATPVPPEEQANLWLVEAEISPELQKRLGRKKKRRKRKKEVCPLAPPPELHPPAPAPSTIPRLPQLPRQKCLVAAGAWGAGDSCRQGAWTLVSNPFCPEPSPPQDPFLPSAPAPVAWAHGRRQGLGPIHSRTNLMDTELMDADSDF | MAAARPARGPELPLLGLLLLLLLGDPGRGAASSGNATGPGPRSAGGSARRSAAVTGPPPPLSHCGRAAPCEPLRYNVCLGSVLPYGATSTLLAGDSDSQEEAHGKLVLWSGLRNAPRCWAVIQPLLCAVYMPKCENDRVELPSRTLCQATRGPCAIVERERGWPDFLRCTPDRFPEGCTNEVQNIKFNSSGQCEVPLVRTDNPKSWYEDVEGCGIQCQNPLFTEAEHQDMHSYIAAFGAVTGLCTLFTLATFVADWRNSNRYPAVILFYVNACFFVGSIGWLAQFMDGARREIVCRADGTMRLGEPTSNETLSCVIIFVIVYYALMAGVVWFVVLTYAWHTSFKALGTTYQPLSGKTSYFHLLTWSLPFVLTVAILAVAQVDGDSVSGICFVGYKNYRYRAGFVLAPIGLVLIVGGYFLIRGVMTLFSIKSNHPGLLSEKAASKINETMLRLGIFGFLAFGFVLITFSCHFYDFFNQAEWERSFRDYVLCQANVTIGLPTKQPIPDCEIKNRPSLLVEKINLFAMFGTGIAMSTWVWTKATLLIWRRTWCRLTGQSDDEPKRIKKSKMIAKAFSKRHELLQNPGQELSFSMHTVSHDGPVAGLAFDLNEPSADVSSAWAQHVTKMVARRGAILPQDISVTPVATPVPPEEQANLWLVEAEISPELQKRLGRKKKRRKRKKEVCPLAPPPELHPPAPAPSTIPRLPQLPRQKCLVAAGAWGAGDSCRQGAWTLVSNPFCPEPSPPQDPFLPSAPAPVAWAHGRRQGLGPIHSRTNLMDTELMDADSDF |
| **4NEF** | High | 4OJ2 | P411 | AQP2 | P411 | | A | C | 6 | R.{2}FA.{2}L.{3}L.{2}FF.{2}GS.{11}LQ.{2} | VL.{86}Q.{2}TV.{2}FL.LQ.{2}LC.{2}AS.{1 | 24 | 22 | 152 | 179 | MWELRSIAFSRAVFAEFLATLLFVFFGLGSALNWPQALPSVLQIAMAFGLGIGTLVQALGHISGAHINPAVTVACLVGCHVSVLRAAFYVA | MWELRSIAFSRAVFAEFLATLLFVFFGLGSALNWPQALPSVLQIAMAFGLGIGTLVQALGHISGAHINPAVTVACLVGCHVSVLRAAFYVA |

| | | | | | | | | | | | | | | | | | |
|---|---|---|---|---|---|---|---|---|---|---|---|---|---|---|---|---|---|
| **(X-ray)** | (Bio) | | 81 | | 81 | | | | | MA.{2}LG.{2}T.VQALG.{42}LL.{57}L | 4}S.{2}FS.AL.{2}LL.{43}LY | | | | | AQLLGAVAGAALLHEITPADIRGDLAVNALSNSTTAGQAVTVELFLTLQLVLCIFASTDERRGENPGTPALSIGFSVALGHLLGIHYTGCSMNPARSLAPAVVTGKFDDHWVFWIGPLVGAILGSLLYNYVLFPPAKSLSERLAVLKGLEPDTDWEEREVRRRQSVELHSPQSLPRGTKA | AQLLGAVAGAALLHEITPADIRGDLAVNALSNSTTAGQAVTVELFLTLQLVLCIFASTDERRGENPGTPALSIGFSVALGHLLGIHYTGCSMNPARSLAPAVVTGKFDDHWVFWIGPLVGAILGSLLYNYVLFPPAKSLSERLAVLKGLEPDTDWEEREVRRRQSVELHSPQSLPRGTKA |
| **4OH3 (X-ray)** | Low (Bio) | - | Q05085 | NPF6.3 | Q05085 | | A | B | 12 | I.{2}A.{3}T.{2}SI.{5}I.{97}W.{6}F.{2}V.{2}LS.{2}L | AI.{2}AI.{2}T.{3}I.{5}I.{97}W.{6}F.{2}V.{2}LS.{2}L | 12 | 13 | 133 | 134 | MSLPETKSDDILLDAWDFQGRPADRSKTGGWASAAMILCIEAVERLTTLGIGVNLVTYLTGTMHLGNATAANTVTNFLGTSFMLCLLGGFIADTFLGRYLTIAIFAAIQATGVSILTLSTIIPGLRPPRCNPTTSSHCEQASGIQLTVLYLALYLTALGTGGVKASVSGFGSDQFDETEPKERSKMTYFFNRFFFCINVGSLLAVTVLVYVQDDVGRKWGYGICAFAIVLALSVFLAGTNRYRFKKLIGSPMTQVAAVIVAAWRNRKLELPADPSYLYDVDDIIAAEGSMKGKQKLPHTEQFRSLDKAAIRDQEAGVTSNVFNKWTLSTLTDVEEVKQIVRMLPIWATCILFWTVHAQLTTLSVAQSETLDRSIGSFEIPPASMAVFYVGGLLLTTAVYDRVAIRLCKKLFNYPHGLRPLQRIGLGLFFGSMAMAVAALVELKRLRTAHAHGPTVKTLPLGFYLLIPQYLIVGIGEALIYTGQLDFFLRECPKGMKGMSTGLLLSTLALGFFFSSVLVTIVEKFTGKAHPWIADDLNKGRLYNFYWLVAVLVALNFLIFLVFSKWYVYKEKRLAEVGIELDDEPSIPMGH | MSLPETKSDDILLDAWDFQGRPADRSKTGGWASAAMILCIEAVERLTTLGIGVNLVTYLTGTMHLGNATAANTVTNFLGTSFMLCLLGGFIADTFLGRYLTIAIFAAIQATGVSILTLSTIIPGLRPPRCNPTTSSHCEQASGIQLTVLYLALYLTALGTGGVKASVSGFGSDQFDETEPKERSKMTYFFNRFFFCINVGSLLAVTVLVYVQDDVGRKWGYGICAFAIVLALSVFLAGTNRYRFKKLIGSPMTQVAAVIVAAWRNRKLELPADPSYLYDVDDIIAAEGSMKGKQKLPHTEQFRSLDKAAIRDQEAGVTSNVFNKWTLSTLTDVEEVKQIVRMLPIWATCILFWTVHAQLTTLSVAQSETLDRSIGSFEIPPASMAVFYVGGLLLTTAVYDRVAIRLCKKLFNYPHGLRPLQRIGLGLFFGSMAMAVAALVELKRLRTAHAHGPTVKTLPLGFYLLIPQYLIVGIGEALIYTGQLDFFLRECPKGMKGMSTGLLLSTLALGFFFSSVLVTIVEKFTGKAHPWIADDLNKGRLYNFYWLVAVLVALNFLIFLVFSKWYVYKEKRLAEVGIELDDEPSIPMGH |
| **4OR2 (X-ra** | Low (Cryst | - | Q13255 | GRM1 | Q13255 | | A | B | 7 (GPCR) | L.{2}LF.{2}LI | L.{2}LF.{2}LI | 5 | 5 | 9 | 9 | MVGLLLFFFPAIFLEVSLLPRSPGRKVLLAGASSQRSVARMDGDVIIGALFSVHHQPPAEKVPERKCGEIREQYGIQRVEAMFHTLDKINADPVLLPNITLGSEIRDSCWHSSVALEQSIEFIRDSLISIRDEKDGINRCLPDGQSLPPGRTKKPIAGVIGPGSSSVAIQVQNLLQLFDIPQIAYSA | MVGLLLFFFPAIFLEVSLLPRSPGRKVLLAGASSQRSVARMDGDVIIGALFSVHHQPPAEKVPERKCGEIREQYGIQRVEAMFHTLDKINADPVLLPNITLGSEIRDSCWHSSVALEQSIEFIRDSLISIRDEKDGINRCLPDGQSLPPGRTKKPIAGVIGPGSSSVAIQVQNLLQLFDIPQIAYSA |

| | | | | | | | | | | | | | | | | | |
|---|---|---|---|---|---|---|---|---|---|---|---|---|---|---|---|---|---|
| **y)** | allographic) | | | | | | | | | | | | | | | TSIDLSDKTLYKYFLRVVPS<br>DTLQARAMLDIVKRYNW<br>TYVSAVHTEGNYGESGM<br>DAFKELAAQEGLCIAHSD<br>KIYSNAGEKSFDRLLRKLR<br>ERLPKARVVVCFCEGMTV<br>RGLLSAMRRLGVVGEFSL<br>IGSDGWADRDEVIEGYEV<br>EANGGITIKLQSPEVRSFD<br>DYFLKLRLDTNTRNPWFP<br>EFWQHRFQCRLPGHLLE<br>NPNFKRICTGNESLEENY<br>VQDSKMGFVINAIYAMA<br>HGLQNMHHALCPGHVG<br>LCDAMKPIDGSKLLDFLIK<br>SSFIGVSGEEVWFDEKGD<br>APGRYDIMNLQYTEANR<br>YDYVHVGTWHEGVLNID<br>DYKIQMNKSGVVRSVCSE<br>PCLKGQIKVIRKGEVSCC<br>WICTACKENEYVQDEFTC<br>KACDLGWWPNADLTGC<br>EPIPVRYLEWSNIESIIAIAF<br>SCLGILVTLFVTLIFVLYRD<br>TPVVKSSSRELCYIILAGIFL<br>GYVCPFTLIAKPTTTSCYL<br>QRLLVGLSSAMCYSALVT<br>KTNRIARILAGSKKKICTRK<br>PRFMSAWAQVIIASILISV<br>QLTLVVTLIIMEPPMPILS<br>YPSIKEVYLICNTSNLGVV<br>APLGYNGLLIMSCTYYAFK<br>TRNVPANFNEAKYIAFTM<br>YTTCIIWLAFVPIYFGSNYK<br>IITTCFAVSLSVTVALGCM<br>FTPKMYIIIAKPERNVRSA<br>FTTSDVVRMHVGDGKLP<br>CRSNTFLNIFRRKKAGAG<br>NANSNGKSVSWSEPGGG<br>QVPKGQHMWHRLSVHV<br>KTNETACNQTAVIKPLTKS<br>YQGSGKSLTFSDTSTKTLY<br>NVEEEEDAQPIRFSPPGS<br>PSMVVHRRVPSAATTPPL<br>PSHLTAEETPLFLAEPALP<br>KGLPPPLQQQQQPPPQQ<br>KSLMDQLQGVVSNFSTAI<br>PDFHAVLAGPGGPGNGL<br>RSLYPPPPPPQHLQMLPL<br>QLSTFGEELVSPPADDDD<br>DSERFKLLQEYVYEHERE<br>GNTEEDELEEEEEDLQAA<br>SKLTPDDSPALTPPSPFRD | TSIDLSDKTLYKYFLRVVPS<br>DTLQARAMLDIVKRYNW<br>TYVSAVHTEGNYGESGM<br>DAFKELAAQEGLCIAHSD<br>KIYSNAGEKSFDRLLRKLR<br>ERLPKARVVVCFCEGMTV<br>RGLLSAMRRLGVVGEFSL<br>IGSDGWADRDEVIEGYEV<br>EANGGITIKLQSPEVRSFD<br>DYFLKLRLDTNTRNPWFP<br>EFWQHRFQCRLPGHLLE<br>NPNFKRICTGNESLEENY<br>VQDSKMGFVINAIYAMA<br>HGLQNMHHALCPGHVG<br>LCDAMKPIDGSKLLDFLIK<br>SSFIGVSGEEVWFDEKGD<br>APGRYDIMNLQYTEANR<br>YDYVHVGTWHEGVLNID<br>DYKIQMNKSGVVRSVCSE<br>PCLKGQIKVIRKGEVSCC<br>WICTACKENEYVQDEFTC<br>KACDLGWWPNADLTGC<br>EPIPVRYLEWSNIESIIAIAF<br>SCLGILVTLFVTLIFVLYRD<br>TPVVKSSSRELCYIILAGIFL<br>GYVCPFTLIAKPTTTSCYL<br>QRLLVGLSSAMCYSALVT<br>KTNRIARILAGSKKKICTRK<br>PRFMSAWAQVIIASILISV<br>QLTLVVTLIIMEPPMPILS<br>YPSIKEVYLICNTSNLGVV<br>APLGYNGLLIMSCTYYAFK<br>TRNVPANFNEAKYIAFTM<br>YTTCIIWLAFVPIYFGSNYK<br>IITTCFAVSLSVTVALGCM<br>FTPKMYIIIAKPERNVRSA<br>FTTSDVVRMHVGDGKLP<br>CRSNTFLNIFRRKKAGAG<br>NANSNGKSVSWSEPGGG<br>QVPKGQHMWHRLSVHV<br>KTNETACNQTAVIKPLTKS<br>YQGSGKSLTFSDTSTKTLY<br>NVEEEEDAQPIRFSPPGS<br>PSMVVHRRVPSAATTPPL<br>PSHLTAEETPLFLAEPALP<br>KGLPPPLQQQQQPPPQQ<br>KSLMDQLQGVVSNFSTAI<br>PDFHAVLAGPGGPGNGL<br>RSLYPPPPPPQHLQMLPL<br>QLSTFGEELVSPPADDDD<br>DSERFKLLQEYVYEHERE<br>GNTEEDELEEEEEDLQAA<br>SKLTPDDSPALTPPSPFRD |

| | | | | | | | | | | | | | | | | | |
|---|---|---|---|---|---|---|---|---|---|---|---|---|---|---|---|---|---|
| | | | | | | | | | | | | | | | | SVASGSSVPSSPVSESVLCTPPNVSYASVILRDYKQSSSTL | SVASGSSVPSSPVSESVLCTPPNVSYASVILRDYKQSSSTL |
| **4TWK (X-ray)** | High (Bio) | - | O95069 | KCNK2 | O95069 | | A | B | 4 | W.{2}V.{2}IF.VV.{2}YL.{2}GATVF.{88}GGKIF.I.YALLGIPL.{2}FL.{2}G.{2}DQ.{94}W.{9}F.{3}L.{2}IG.{2}LR | W.{2}V.{2}IF.{2}VV.YLI.GATVF.{88}GGKIFCI.YALLGIPL.{2}FL.AG.{2}DQ.{94}W.{3}L.{5}F.{3}L.{2}IG.WLR | 39 | 44 | 254 | 254 | MLPSASRERPGYRAGVAAPDLLDPKSAAQNSKPRLSFSTKPTVLASRVESDTTINVMKWKTVSTIFLVVVLYLIIGATVFKALEQPHEISQRTTIVIQKQTFISQHSCVNSTELDELIQQIVAAINAGIIPLGNTSNQISHWDLGSSFFFAGTVITTIGFGNISPRTEGGKIFCIIYALLGIPLFGFLLAGVGDQLGTIFGKGIAKVEDTFIKWNVSQTKIRIISTIIFILFGCVLFVALPAIIFKHIEGWSALDAIYFVVITLTTIGFGDYVAGGSDIEYLDFYKPVVWFWILVGLAYFAAVLSMIGDWLRVISKKTKEEVGEFRAHAAEWTANVTAEFKETRRRLSVEIYDKFQRATSIKRKLSAELAGNHNQELTPCRRTLSVNHLTSERDVLPPLLKTESIYLNGLTPHCAGEEIAVIENIK | MLPSASRERPGYRAGVAAPDLLDPKSAAQNSKPRLSFSTKPTVLASRVESDTTINVMKWKTVSTIFLVVVLYLIIGATVFKALEQPHEISQRTTIVIQKQTFISQHSCVNSTELDELIQQIVAAINAGIIPLGNTSNQISHWDLGSSFFFAGTVITTIGFGNISPRTEGGKIFCIIYALLGIPLFGFLLAGVGDQLGTIFGKGIAKVEDTFIKWNVSQTKIRIISTIIFILFGCVLFVALPAIIFKHIEGWSALDAIYFVVITLTTIGFGDYVAGGSDIEYLDFYKPVVWFWILVGLAYFAAVLSMIGDWLRVISKKTKEEVGEFRAHAAEWTANVTAEFKETRRRLSVEIYDKFQRATSIKRKLSAELAGNHNQELTPCRRTLSVNHLTSERDVLPPLLKTESIYLNGLTPHCAGEEIAVIENIK |
| **4WFE (X-ray)** | High (Bio) | - | Q9NYG8 | KCNK4 | Q9NYG8 | | A | B | 4 | L.{2}L.{3}V.{2}YL.{2}GA.VF.{65}L.{2}AF.FSG.{2}I.{12}D.GR.FCI.Y.LVGIPL.{2}IL.{2}GV.{2}R.{58}L.{3}Y.{2}I.{3}T.{45}L | L.{2}LL.LVL.YL.{2}GA.VFR.L.{62}L.{2}AF.FSG.{2}I.{14}GR.FCI.Y.{2}VGI.L.{2}IL.{2}G.{3}R.{58}L.{3}Y.{2}I.{3}T.{24}W.{9}F.{3}L.{2}IG.{2}LR | 39 | 46 | 252 | 253 | MRSTTLLALLALVLLYLVSGALVFRALEQPHEQQAQRELGEVREKFLRAHPCVSDQELGLLIKEVADALGGGADPETNSTSNSSHSAWDLGSAFFFSGTIITTIGYGNVALRTDAGRLFCIFYALVGIPLFGILLAGVGDRLGSSLRHGIGHIEAIFLKWHVPPELVRVLSAMLFLLIGCLLFVLTPTFVFCYMEDWSKLEAIYFVIVTLTTVGFGDYVAGADPRQDSPAYQPLVWFWILLGLAYFASVLTTIGNWLRVVSRRTRAEMGGLTAQAASWTGTVTARVTQRAGPAAPPPEKEQPLLPPPPCPAQPLGRPRSPSPPEKAQPPSPPTASALDYPSENLAFIDESSDTQSERGCPLPRAPRGRRRPNPPRKPVRPRGPGRPRDKGVPV | MRSTTLLALLALVLLYLVSGALVFRALEQPHEQQAQRELGEVREKFLRAHPCVSDQELGLLIKEVADALGGGADPETNSTSNSSHSAWDLGSAFFFSGTIITTIGYGNVALRTDAGRLFCIFYALVGIPLFGILLAGVGDRLGSSLRHGIGHIEAIFLKWHVPPELVRVLSAMLFLLIGCLLFVLTPTFVFCYMEDWSKLEAIYFVIVTLTTVGFGDYVAGADPRQDSPAYQPLVWFWILLGLAYFASVLTTIGNWLRVVSRRTRAEMGGLTAQAASWTGTVTARVTQRAGPAAPPPEKEQPLLPPPPCPAQPLGRPRSPSPPEKAQPPSPPTASALDYPSENLAFIDESSDTQSERGCPLPRAPRGRRRPNPPRKPVRPRGPGRPRDKGVPV |
| **4WO1** | Low | 2L34o | O43939 | TYRO | O43939 | | B | C | 1 | P.{2}L.{3}V.{2}DL.{6}A.{3}Y | L.{2}I.{3}D.{3}T.{2}I.{3}V.{2}L | 7 | 7 | 23 | 22 | MGGLEPCSRLLLLPLLLAVSGLRPVQAQAQSDCSCSTVSPGVLAGIVMGDLVLTVLIALAVYFLGRLVPRGRGAAEAATRKQRITETESPYQE | MGGLEPCSRLLLLPLLLAVSGLRPVQAQAQSDCSCSTVSPGVLAGIVMGDLVLTVLIALAVYFLGRLVPRGRGAAEAATRKQRITETESPYQE |

| | | | | | | | | | | | | | | | | | |
|---|---|---|---|---|---|---|---|---|---|---|---|---|---|---|---|---|---|
| **(X-ray)** | (Medium) | r4WOL | 14 | BP | 14 | | | | | | | | | | | LQGQRSDVYSDLNTQRPYYK | LQGQRSDVYSDLNTQRPYYK |
| **4X5T (X-ray)** | Low (Bio) | 2M6B | P23415 | GLRA1 | P23415 | | A | B | 4 | Y.{2}I.{7}L.{2}IL.{16}GL.{2}T.{2}LT.{2}T.{2}SG.R | I.{3}L.{2}T.{3}S.{2}R.{19}L.{3}F.{2}L.{3}A | 14 | 9 | 50 | 46 | MYSFNTLRLYLWETIVFFSLAASKEAEAARSAPKPMSPSDFLDKLMGRTSGYDARIRPNFKGPPVNVSCNIFINSFGSIAETTMDYRVNIFLRQQWNDPRLAYNEYPDDSLDLDPSMLDSIWKPDLFFANEKGAHFHEITTDNKLLRISRNGNVLYSIRITLTLACPMDLKNFPMDVQTCIMQLESFGYTMNDLIFEWQEQGAVQVADGLTLPQFILKEEKDLRYCTKHYNTGKFTCIEARFHLERQMGYYLIQMYIPSLLIVILSWISFWINMDAAPARVGLGITTVLTMTTQSSGSRASLPKVSYVKAIDIWMAVCLLFVFSALLEYAAVNFVSRQHKELLRFRRKRRHHKSPMLNLFQEDEAGEGRFNFSAYGMGPACLQAKDGISVKGANNSNTTNPPPAPSKSPEEMRKLFIQRAKKIDKISRIGFPMAFLIFNMFYWIIYKIVRREDVHNQ | MYSFNTLRLYLWETIVFFSLAASKEAEAARSAPKPMSPSDFLDKLMGRTSGYDARIRPNFKGPPVNVSCNIFINSFGSIAETTMDYRVNIFLRQQWNDPRLAYNEYPDDSLDLDPSMLDSIWKPDLFFANEKGAHFHEITTDNKLLRISRNGNVLYSIRITLTLACPMDLKNFPMDVQTCIMQLESFGYTMNDLIFEWQEQGAVQVADGLTLPQFILKEEKDLRYCTKHYNTGKFTCIEARFHLERQMGYYLIQMYIPSLLIVILSWISFWINMDAAPARVGLGITTVLTMTTQSSGSRASLPKVSYVKAIDIWMAVCLLFVFSALLEYAAVNFVSRQHKELLRFRRKRRHHKSPMLNLFQEDEAGEGRFNFSAYGMGPACLQAKDGISVKGANNSNTTNPPPAPSKSPEEMRKLFIQRAKKIDKISRIGFPMAFLIFNMFYWIIYKIVRREDVHNQ |
| **5A63 (EM)** | High (Bio) | - | P49768 | PSEN1 | Q96BI3 | APH1A | B | C | 9(B),7(C) | DWN.{2}IA.{2}V.{2}LI.{24}F.{3}F.{2}A.{2}YLV.{2}FM | VI.{2}V.{3}F.WLV.LL.{2}SV.WF.{17}L.{6}SV.{2}Q.{44}IS.VFS.IN.{22}TS.{2}LT.{32}H.{3}S.{2}TF.N | 16 | 33 | 55 | 176 | MTELPAPLSYFQNAQMSEDNHLSNTVRSQNDNRERQEHNDRRSLGHPEPLSNGRPQGNSRQVVEQDEEEDEELTLKYGAKHVIMLFVPVTLCMVVVVATIKSVSFYTRKDGQLIYTPFTEDTETVGQRALHSILNAAIMISVIVVMTILLVVLYKYRCYKVIHAWLIISSLLLLFFFSFIYLGEVFKTYNVAVDYITVALLIWNFGVVGMISIHWKGPLRLQQAYLIMISALMALVFIKYLPEWTAWLILAVISVYDLVAVLCPKGPLRMLVETAQERNETLFPALIYSSTMVWLVNMAEGDPEAQRRVSKNSKYNAESTERESQDTVAENDDGGFSEEWEAQR | MGAAVFFGCTFVAFGPAFALFLITVAGDPLRVIILVAGAFFWLVSLLLASVVWFILVHVTDRSDARLQYGLLIFGAAVSVLLQEVFRFAYYKLLKKADEGLASLSEDGRSPISIRQMAYVSGLSFGIISGVFSVINILADALGPGVVGIHGDSPYYFLTSAFLTAAIILLHTFWGVVFFDACERRRYWALGLVVGSHLLTSGLTFLNPWYEASLLPIYAVTVSMGLWAFITAGGSLRSIQRSLLCRRQEDSRVMVYSALRIPPED |

| | | | | | | | | | | | | | | | | | |
|---|---|---|---|---|---|---|---|---|---|---|---|---|---|---|---|---|---|
| | | | | | | | | | | | | | | | | DSHLGPHRSTPESRAAVQELSSSILAGEDPEERGVKLGLGDFIFYSVLVGKASATASGDWNTTIACFVAILIGLCLTLLLLAIFKKALPALPISITFGLVFYFATDYLVQPFMDQLAFHQFYI | |
| **5A63 (EM)** | High (Bio) | 2N7Q or 2N7R for Q92542 | Q92542 | NCSTN | Q96BI3 | APH1A | A | C | 1(A),7(C) | E.{2}TL.VGF.IL.{2}SL.{2}TY | AV.{2}G.{2}FV.{2}GP.{2}AL.{2}I.{112}NI.{19}FL.{3}F | 12 | 15 | 20 | 159 | MATAGGGSGADPGSRGLLRLLSFCVLLAGLCRGNSVERKIYIPLNKTAPCVRLLNATHQIGCQSSISGDTGVIHVVEKEEDLQWVLTDGPNPPYMVLLESKHFTRDLMEKLKGRTSRIAGLAVSLTKPSPASGFSPSVQCPNDGFGVYSNSYGPEFAHCREIQWNSLGNGLAYEDFSFPIFLLEDENETKVIKQCYQDHNLSQNGSAPTFPLCAMQLFSHMHAVISTATCMRRSSIQSTFSINPEIVCDPLSDYNVWSMLKPINTTGTLKPDDRVVVAATRLDSRSFFWNVAPGAESAVASFVTQLAAAEALQKAPDVTTLPRNVMFVFFQGETFDYIGSSRMVYDMEKGKFPVQLENVDSFVELGQVALRTSLELWMHTDPVSQKNESVRNQVEDLLATLEKSGAGVPAVILRRPNQSQPLPPSSLQRFLRARNISGVVLADHSGAFHNKYYQSIYDTAENINVSYPEWLSPEEDLNFVTDTAKALADVATVLGRALYELAGGTNFSDTVQADPQTVTRLLYGFLIKANNSWFQSILRQDLRSYLGDGPLQHYIAVSSPTNTTYVVQYALANLTGTVVNLTREQCQDPSKVPSENKDLYEYSWVQGPLHSNETDRLPRCVRSTARLARALSPAFELSQWSSTEYSTWTESRWKDIRARIFLIASKELELITLTVGFGILIFSLIVTYCINAKADVLFIAPREPGAVSY | MGAAVFFGCTFVAFGPAFALFLITVAGDPLRVIILVAGAFFWLVSLLLASVVWFILVHVTDRSDARLQYGLLIFGAAVSVLLQEVFRFAYYKLLKKADEGLASLSEDGRSPISIRQMAYVSGLSFGIISGVFSVINILADALGPGVVGIHGDSPYYFLTSAFLTAAIILLHTFWGVVFFDACERRRYWALGLVVGSHLLTSGLTFLNPWYEASLLPIYAVTVSMGLWAFITAGGSLRSIQRSLLCRRQEDSRVMVYSALRIPPED |
| **5A63** | High | - | P49768 | PSEN1 | Q9NZ42 | PSEN | B | D | 9(B),1 | F.{3}G.{2}F.{8}YITV.L.{2}WN.{2}VV.{2}I.{9}L | W.{3}L.{2}W.{3}FQ | 14 | 5 | 43 | 13 | MTELPAPLSYFQNAQMSEDNHLSNTVRSQNDNRERQEHNDRRSLGHPEPLSNGRPQGNSRQVVEQDEEEDEELTLKYGAKHVIMLFVPVTLCMVVVVATIKSVSF | MNLERVSNEEKLNLCRKYYLGGFAFLPFLWLVNIFWFFREAFLVPAYTEQSQIKGYVWRSAVGFLFWVIVLTSWITIFQIYRPRWGALGDYLSFTIPLGTP |

| | | | | | | | | | | | | | | | | | |
|---|---|---|---|---|---|---|---|---|---|---|---|---|---|---|---|---|---|
| **(EM)** | (Bio) | | | | | | | | (D) | | | | | | | YTRKDGQLIYTPFTEDTETVGQRALHSILNAAIMISVIVVMTILLVVLYKYRCYKVIHAWLIISSLLLLFFFSFIYLGEVFKTYNVAVDYITVALLIWNFGVVGMISIHWKGPLRLQQAYLIMISALMALVFIKYLPEWTAWLILAVISVYDLVAVLCPKGPLRMLVETAQERNETLFPALIYSSTMVWLVNMAEGDPEAQRRVSKNSKYNAESTERESQDTVAENDDGGFSEEWEAQRDSHLGPHRSTPESRAAVQELSSSILAGEDPEERGVKLGLGDFIFYSVLVGKASATASGDWNTTIACFVAILIGLCLTLLLLAIFKKALPALPISITFGLVFYFATDYLVQPFMDQLAFHQFYI | |
| **5AEX (X-ray)** | High (Bio) | - | P41948 | MEP2 | P41948 | | A | B | 10 | G.{3}A.{2}WI.{2}P.{18}LW.{2}MM.SA.{2}I.{3}F.{45}V.{3}M.{18}LF.{2}M | VT.{2}V.{3}V.{2}WF.{2}MF.{17}M.{2}NL.{2}A.{6}M.{12}T.{2}L.{23}V | 18 | 16 | 119 | 89 | MSYNFTGTPTGEGTGGNSLTTDLNTQFDLANMGWIGVASAGVWIMVPGIGLLYSGLSRKKHALSLLWASMMASAVCIFQWFFWGYSLAFSHNTRGNGFIGTLEFFGFRNVLGAPSSVSSLPDILFAVYQGMFAAVTGALMLGGACERARLFPMMVFLFLWMTIVYCPIACWVWNAEGWLVKLGSLDYAGGLCVHLTSGHGGLVYALILGKRNDPVTRKGMPKYKPHSVTSVVLGTVFLWFGWMFFNGGSAGNATIRAWYSIMSTNLAAACGGLTWMVIDYFRCGRKWTTVGLCSGIIAGLVGITPAAGFVPIWSAVVIGVVTGAGCNLAVDLKSLLRIDDGLDCYSIHGVGGCIGSVLTGIFAADYVNATAGSYISPIDGGWINHHYKQVGYQLAGICAALAWTVTVTSILLLTMNAIPFLKLRLSADEEELGTDAAQIGEFTYEESTAYIPEPIRSKTSAQMPPPHENIDDKIVGNTDAEKNSTPSDASSTKNTDHIV | MSYNFTGTPTGEGTGGNSLTTDLNTQFDLANMGWIGVASAGVWIMVPGIGLLYSGLSRKKHALSLLWASMMASAVCIFQWFFWGYSLAFSHNTRGNGFIGTLEFFGFRNVLGAPSSVSSLPDILFAVYQGMFAAVTGALMLGGACERARLFPMMVFLFLWMTIVYCPIACWVWNAEGWLVKLGSLDYAGGLCVHLTSGHGGLVYALILGKRNDPVTRKGMPKYKPHSVTSVVLGTVFLWFGWMFFNGGSAGNATIRAWYSIMSTNLAAACGGLTWMVIDYFRCGRKWTTVGLCSGIIAGLVGITPAAGFVPIWSAVVIGVVTGAGCNLAVDLKSLLRIDDGLDCYSIHGVGGCIGSVLTGIFAADYVNATAGSYISPIDGGWINHHYKQVGYQLAGICAALAWTVTVTSILLLTMNAIPFLKLRLSADEEELGTDAAQIGEFTYEESTAYIPEPIRSKTSAQMPPPHENIDDKIVGNTDAEKNSTPSDASSTKNTDHIV |
| **5CFB** | High | - | O753 | GLRA3 | O753 | | A | B | 4 | I.{3}L.{2}T.{6}R.{12}D.{2}MA.{6}F.{2}LL | GYY.IQ.{6}L.{2}IL.{2}V.{13}A.{3} | 10 | 13 | 43 | 46 | MAHVRHFRTLVSGFYFWEAALLLSLVATKETDSARSRSAPMSPSDFLDKLMGRTSGYDARIRPNFKGPPVNVTCNIFINSFGSIAETTMD | MAHVRHFRTLVSGFYFWEAALLLSLVATKETDSARSRSAPMSPSDFLDKLMGRTSGYDARIRPNFKGPPVNVTCNIFINSFGSIAETTMD |

| | | | | | | | | | | | | | | | | | |
|---|---|---|---|---|---|---|---|---|---|---|---|---|---|---|---|---|---|
| **(X-ray)** | (Bio) | | 11 | | 11 | | | | | | T.{3}T.{3}Q | | | | | YRVNIFLRQKWNDPRLAYSEYPDDSLDLDPSMLDSIWKPDLFFANEKGANFHEVTTDNKLLRIFKNGNVLYSIRLTLTLSCPMDLKNFPMDVQTCIMQLESFGYTMNDLIFEWQDEAPVQVAEGLTLPQFLLKEEKDLRYCTKHYNTGKFTCIEVRFHLERQMGYYLIQMYIPSLLIVILSWVSFWINMDAAPARVALGITTVLTMTTQSSGSRASLPKVSYVKAIDIWMAVCLLFVFSALLEYAAVNFVSRQHKELLRFRRKRKNKTEAFALEKFYRFSDMDDEVRESRFSFTAYGMGPCLQAKDGMTPKGPNHPVQVMPKSPDEMRKVFIDRAKKIDTISRACFPLAFLIFNIFYWVIYKILRHEDIHQQQD | YRVNIFLRQKWNDPRLAYSEYPDDSLDLDPSMLDSIWKPDLFFANEKGANFHEVTTDNKLLRIFKNGNVLYSIRLTLTLSCPMDLKNFPMDVQTCIMQLESFGYTMNDLIFEWQDEAPVQVAEGLTLPQFLLKEEKDLRYCTKHYNTGKFTCIEVRFHLERQMGYYLIQMYIPSLLIVILSWVSFWINMDAAPARVALGITTVLTMTTQSSGSRASLPKVSYVKAIDIWMAVCLLFVFSALLEYAAVNFVSRQHKELLRFRRKRKNKTEAFALEKFYRFSDMDDEVRESRFSFTAYGMGPCLQAKDGMTPKGPNHPVQVMPKSPDEMRKVFIDRAKKIDTISRACFPLAFLIFNIFYWVIYKILRHEDIHQQQD |
| **5CTG (X-ray)** | Low (Bio) | - | Q5N8J1 | SWEET2B | Q5N8J1 | | A | B | 7 (GPCR) | M.{3}G.{3}L.{2}C.{3}LV.HA.{2}FF.{8}F | L.{4}F.{2}YF.{2}GL.{2}I.{3}M.{3}L.{2}Y | 11 | 10 | 33 | 28 | MDSLYDISCFAAGLAGNIFALALFLSPVTTFKRILKAKSTERFDGLPYLFSLLNCLICLWYGLPWVADGRLLVATVNGIGAVFQLAYICLFIFYADSRKTRMKIIGLLVLVVCGFALVSHASVFFFDQPLRQQFVGAVSMASLISMFASPLAVMGVVIRSESVEFMPFYLSLSTFLMSASFALYGLLLRDFFIYFPNGLGLILGAMQLALYAYYSRKWRGQDSSAPLLLA | MDSLYDISCFAAGLAGNIFALALFLSPVTTFKRILKAKSTERFDGLPYLFSLLNCLICLWYGLPWVADGRLLVATVNGIGAVFQLAYICLFIFYADSRKTRMKIIGLLVLVVCGFALVSHASVFFFDQPLRQQFVGAVSMASLISMFASPLAVMGVVIRSESVEFMPFYLSLSTFLMSASFALYGLLLRDFFIYFPNGLGLILGAMQLALYAYYSRKWRGQDSSAPLLLA |
| **5IRX (EM)** | High (Bio) | - | O35433 | Trpv1 | O35433 | | A | C | 6 | R.{3}V.{2}VF.FG.{2}TA.VTL.{59}V.{2}IL.L.{2}VI.{2}YILL.NM.{2}AL | AY.{82}E.VA.{2}VF.LA.{2}WT.{124}L.{3}I.{2}M | 25 | 14 | 103 | 231 | MEQRASLDSEESESPPQENSCLDPPDRDPNCKPPPVKPHIFTTRSRTRLFGKGDSEEASPLDCPYEEGGLASCPIITVSSVLTIQRPGDGPASVRPSSQDSVSAGEKPPRLYDRRSIFDAVAQSNCQELESLLPFLQRSKKRLTDSEFKDPETGKTCLLKAMLNLHNGQNDTIALLLDVARKTDSLKQFVNASYTDSYYKGQTALHIAIERRNMTLVTLLVENGADVQAAANGDFFKKTKGRPGFYFGELPLSLAACTNQLAIVKFLLQNSWQPADISARDSVGNTVLHALVEVADNTVDNTKFVTSMYNEILILGAKLHPTLKLEEITNRKGLTPLALAASSGKIGVLA | MEQRASLDSEESESPPQENSCLDPPDRDPNCKPPPVKPHIFTTRSRTRLFGKGDSEEASPLDCPYEEGGLASCPIITVSSVLTIQRPGDGPASVRPSSQDSVSAGEKPPRLYDRRSIFDAVAQSNCQELESLLPFLQRSKKRLTDSEFKDPETGKTCLLKAMLNLHNGQNDTIALLLDVARKTDSLKQFVNASYTDSYYKGQTALHIAIERRNMTLVTLLVENGADVQAAANGDFFKKTKGRPGFYFGELPLSLAACTNQLAIVKFLLQNSWQPADISARDSVGNTVLHALVEVADNTVDNTKFVTSMYNEILILGAKLHPTLKLEEITNRKGLTPLALAASSGKIGVLA |

| | | | | | | | | | | | | | | | | | |
|---|---|---|---|---|---|---|---|---|---|---|---|---|---|---|---|---|---|
| | | | | | | | | | | | | | | | | YILQREIHEPECRHLSRKFTEWAYGPVHSSLYDLSCIDTCEKNSVLEVIAYSSSETPNRHDMLLVEPLNRLLQDKWDRFVKRIFYFNFFVYCLYMIIFTAAAYYRPVEGLPPYKLKNTVGDYFRVTGEILSVSGGVYFFFRGIQYFLQRRPSLKSLFVDSYSEILFFVQSLFMLVSVVLYFSQRKEYVASMVFSLAMGWTNMLYYTRGFQQMGIYAVMIEKMILRDLCRFMFVYLVFLFGFSTAVVTLIEDGKNNSLPMESTPHKCRGSACKPGNSYNSLYSTCLELFKFTIGMGDLEFTENYDFKAVFIILLLAYVILTYILLLNMLIALMGETVNKIAQESKNIWKLQRAITILDTEKSFLKCMRKAFRSGKLLQVGFTPDGKDDYRWCFRVDEVNWTTWNTNVGIINEDPGNCEGVKRTLSFSLRSGRVSGRNWKNFALVPLLRDASTRDRHATQQEEVQLKHYTGSLKPEDAEVFKDSMVPGEK | YILQREIHEPECRHLSRKFTEWAYGPVHSSLYDLSCIDTCEKNSVLEVIAYSSSETPNRHDMLLVEPLNRLLQDKWDRFVKRIFYFNFFVYCLYMIIFTAAAYYRPVEGLPPYKLKNTVGDYFRVTGEILSVSGGVYFFFRGIQYFLQRRPSLKSLFVDSYSEILFFVQSLFMLVSVVLYFSQRKEYVASMVFSLAMGWTNMLYYTRGFQQMGIYAVMIEKMILRDLCRFMFVYLVFLFGFSTAVVTLIEDGKNNSLPMESTPHKCRGSACKPGNSYNSLYSTCLELFKFTIGMGDLEFTENYDFKAVFIILLLAYVILTYILLLNMLIALMGETVNKIAQESKNIWKLQRAITILDTEKSFLKCMRKAFRSGKLLQVGFTPDGKDDYRWCFRVDEVNWTTWNTNVGIINEDPGNCEGVKRTLSFSLRSGRVSGRNWKNFALVPLLRDASTRDRHATQQEEVQLKHYTGSLKPEDAEVFKDSMVPGEK |
| **5O8F (X-ray)** | High (Bio) | - | P28472 | GABRB3 | P31644 | GABRA5 | A | B | 4(A),4(B) Note: chimera! | V.{3}T.{3}V.{3}L.{6}I.{22}Y.{3}F.{2}L.{3}A | IQ.{6}M.{2}IL.{2}V.{3}L.{6}A.{3}F.{2}T.{3}T.{130}R | 9 | 12 | 54 | 169 | MWGLAGGRLFGIFSAPVLVAVVCCAQSVNDPGNMSFVKETVDKLLKGYDIRLRPDFGGPPVCVGMNIDIASIDMVSEVNMDYTLTMYFQQYWRDKRLAYSGIPLNLTLDNRVADQLWVPDTYFLNDKKSFVHGVTVKNRMIRLHPDGTVLYGLRITTTAACMMDLRRYPLDEQNCTLEIESYGYTTDDIEFYWRGGDKAVTGVERIELPQFSIVEHRLVSRNVVFATGAYPRLSLSFRLKRNIGYFILQTYMPSILITILSWVSFWINYDASAARVALGITTVLTMTTINTHLRETLPKIPYVKAIDMYLMGCFVFVFLALLEYAFVNYIFFGRGPQRQKKLAEKTAKAKNDRSKSESNRVDAHGNILLTSLEVHNEMNEVSGGIGDTRNSAISFDNSGIQYRKQSMPREGHGRFLGDRSLPHKKTHLRRRSSQLKIKIPDLTDVNAIDR | MDNGMFSGFIMIKNLLLFCISMNLSSHFGFSQMPTSSVKDETNDNITIFTRILDGLLDGYDNRLRPGLGERITQVRTDIYVTSFGPVSDTEMEYTIDVFFRQSWKDERLRFKGPMQRLPLNNLLASKIWTPDTFFHNGKKSIAHNMTTPNKLLRLEDDGTLLYTMRLTISAECPMQLEDFPMDAHACPLKFGSYAYPNSEVVYVWTNGSTKSVVVAEDGSRLNQYHLMGQTVGTENISTSTGEYTIMTAHFHLKRKIGYFVIQTYLPCIMTVILSQVSFWLNRESVPARTVFGVTTVLTMTTLSISARNSLPKVAYATAMDWFIAVCYAFVFSALIEFATVNYFTKRGWAWDGKKALEAAKIKKKREVILNKSTNAFTTGKMSHPPNIPKEQTPAGTSNTTSVSVKPSEEKTSESKKTYNSISKIDKMSRIVFPVLFGTFNLVYWATYLNREPVIKGAASPK |

| | | | | | | | | | | | | | | | | | |
|---|---|---|---|---|---|---|---|---|---|---|---|---|---|---|---|---|---|
| | | | | | | | | | | | | | | | | WSRIVFPFTFSLFNLVYWLYYVN | |
| **5O9H (X-ray)** | Low (Bio/Crystallographic) | 6C1Q or 6C1R | P21730 | C5AR1 | P21730 | | A | B | 7 (GPCR) | L.{2}L.{6}LY.{23}R.{10}L | LL.{5}LY.{22}RR | 6 | 6 | 47 | 33 | MDSFNYTTPDYGHYDDKDTLDLNTPVDKTSNTLRVPDILALVIFAVVFLVGVLGNALVVWVTAFEAKRTINAIWFLNLAVADFLSCLALPILFTSIVQHHHWPFGGAACSILPSLILLNMYASILLLATISADRFLLVFKPIWCQNFRGAGLAWIACAVAWGLALLLTIPSFLYRVVREEYFPPKVLCGVDYSHDKRRERAVAIVRLVLGFLWPLLTLTICYTFILLRTWSRRATRSTKTLKVVVAVVASFFIFWLPYQVTGIMMSFLEPSSPTFLLLKKLDSLCVSFAYINCCINPIIYVVAGQGFQGRLRKSLPSLLRNVLTEESVVRESKSFTRSTVDTMAQKTQAV | MDSFNYTTPDYGHYDDKDTLDLNTPVDKTSNTLRVPDILALVIFAVVFLVGVLGNALVVWVTAFEAKRTINAIWFLNLAVADFLSCLALPILFTSIVQHHHWPFGGAACSILPSLILLNMYASILLLATISADRFLLVFKPIWCQNFRGAGLAWIACAVAWGLALLLTIPSFLYRVVREEYFPPKVLCGVDYSHDKRRERAVAIVRLVLGFLWPLLTLTICYTFILLRTWSRRATRSTKTLKVVVAVVASFFIFWLPYQVTGIMMSFLEPSSPTFLLLKKLDSLCVSFAYINCCINPIIYVVAGQGFQGRLRKSLPSLLRNVLTEESVVRESKSFTRSTVDTMAQKTQAV |
| **5OEK (NMR)** | Low (~Bio) | - | P10912 | GHR | P10912 | | A | B | 1 | FG.FG.{2}VM | FG.FG.{2}VM | 6 | 6 | 9 | 9 | MDLWQLLLTLALAGSSDAFSGSEATAAILSRAPWSLQSVNPGLKTNSSKEPKFTKCRSPERETFSCHWTDEVHHGTKNLGPIQLFYTRRNTQEWTQEWKECPDYVSAGENSCYFNSSFTSIWIPYCIKLTSNGGTVDEKCFSVDEIVQPDPPIALNWTLLNVSLTGIHADIQVRWEAPRNADIQKGWMVLEYELQYKEVNETKWKMMDPILTTSVPVYSLKVDKEYEVRVRSKQRNSGNYGEFSEVLYVTLPQMSQFTCEEDFYFPWLLIIIFGIFGLTVMLFVFLFSKQQRIKMLILPPVPVPKIKGIDPDLLKEGKLEEVNTILAIHDSYKPEFHSDDSWVEFIELDIDEPDEKTEESDTDRLLSSDHEKSHSNLGVKDGDSGRTSCCEPDILETDFNANDIHEGTSEVAQPQRLKGEADLLCLDQKNQNNSPYHDACPATQQPSVIQAEKNKPQPLPTEGAESTHQAAHI | MDLWQLLLTLALAGSSDAFSGSEATAAILSRAPWSLQSVNPGLKTNSSKEPKFTKCRSPERETFSCHWTDEVHHGTKNLGPIQLFYTRRNTQEWTQEWKECPDYVSAGENSCYFNSSFTSIWIPYCIKLTSNGGTVDEKCFSVDEIVQPDPPIALNWTLLNVSLTGIHADIQVRWEAPRNADIQKGWMVLEYELQYKEVNETKWKMMDPILTTSVPVYSLKVDKEYEVRVRSKQRNSGNYGEFSEVLYVTLPQMSQFTCEEDFYFPWLLIIIFGIFGLTVMLFVFLFSKQQRIKMLILPPVPVPKIKGIDPDLLKEGKLEEVNTILAIHDSYKPEFHSDDSWVEFIELDIDEPDEKTEESDTDRLLSSDHEKSHSNLGVKDGDSGRTSCCEPDILETDFNANDIHEGTSEVAQPQRLKGEADLLCLDQKNQNNSPYHDACPATQQPSVIQAEKNKPQPLPTEGAESTHQAAHI |

| | | | | | | | | | | | | | | | | | |
|---|---|---|---|---|---|---|---|---|---|---|---|---|---|---|---|---|---|
| | | | | | | | | | | | | | | | | QLSNPSSLSNIDFYAQVSDITPAGSVVLSPGQKNKAGMSQCDMHPEMVSLCQENFLMDNAYFCEADAKKCIPVAPHIKVESHIQPSLNQEDIYITTESLTTAAGRPGTGEHVPGSEMPVPDYTSIHIVQSPQGLILNATALPLPDKEFLSSCGYVSTDQLNKIMP | QLSNPSSLSNIDFYAQVSDITPAGSVVLSPGQKNKAGMSQCDMHPEMVSLCQENFLMDNAYFCEADAKKCIPVAPHIKVESHIQPSLNQEDIYITTESLTTAAGRPGTGEHVPGSEMPVPDYTSIHIVQSPQGLILNATALPLPDKEFLSSCGYVSTDQLNKIMP |
| **5OHD (NMR)** | Low (~Bio) | - | P10912 | GHR | P10912 | | A | B | 1 | W.{3}II.{2}I.{3}T.{3}F | W.{3}II.{2}I.{3}T.{3}F | 6 | 6 | 17 | 17 | MDLWQLLLTLALAGSSDAFSGSEATAAILSRAPWSLQSVNPGLKTNSSKEPKFTKCRSPERETFSCHWTDEVHHGTKNLGPIQLFYTRRNTQEWTQEWKECPDYVSAGENSCYFNSSFTSIWIPYCIKLTSNGGTVDEKCFSVDEIVQPDPPIALNWTLLNVSLTGIHADIQVRWEAPRNADIQKGWMVLEYELQYKEVNETKWKMMDPILTTSVPVYSLKVDKEYEVRVRSKQRNSGNYGEFSEVLYVTLPQMSQFTCEEDFYFPWLLIIIFGIFGLTVMLFVFLFSKQQRIKMLILPPVPVPKIKGIDPDLLKEGKLEEVNTILAIHDSYKPEFHSDDSWVEFIELDIDEPDEKTEESDTDRLLSSDHEKSHSNLGVKDGDSGRTSCCEPDILETDFNANDIHEGTSEVAQPQRLKGEADLLCLDQKNQNNSPYHDACPATQQPSVIQAEKNKPQPLPTEGAESTHQAAHIQLSNPSSLSNIDFYAQVSDITPAGSVVLSPGQKNKAGMSQCDMHPEMVSLCQENFLMDNAYFCEADAKKCIPVAPHIKVESHIQPSLNQEDIYITTESLTTAAGRPGTGEHVPGSEMPVPDYTSIHIVQSPQGLILNATALPLPDKEFLSSCGYVSTDQLNKIMP | MDLWQLLLTLALAGSSDAFSGSEATAAILSRAPWSLQSVNPGLKTNSSKEPKFTKCRSPERETFSCHWTDEVHHGTKNLGPIQLFYTRRNTQEWTQEWKECPDYVSAGENSCYFNSSFTSIWIPYCIKLTSNGGTVDEKCFSVDEIVQPDPPIALNWTLLNVSLTGIHADIQVRWEAPRNADIQKGWMVLEYELQYKEVNETKWKMMDPILTTSVPVYSLKVDKEYEVRVRSKQRNSGNYGEFSEVLYVTLPQMSQFTCEEDFYFPWLLIIIFGIFGLTVMLFVFLFSKQQRIKMLILPPVPVPKIKGIDPDLLKEGKLEEVNTILAIHDSYKPEFHSDDSWVEFIELDIDEPDEKTEESDTDRLLSSDHEKSHSNLGVKDGDSGRTSCCEPDILETDFNANDIHEGTSEVAQPQRLKGEADLLCLDQKNQNNSPYHDACPATQQPSVIQAEKNKPQPLPTEGAESTHQAAHIQLSNPSSLSNIDFYAQVSDITPAGSVVLSPGQKNKAGMSQCDMHPEMVSLCQENFLMDNAYFCEADAKKCIPVAPHIKVESHIQPSLNQEDIYITTESLTTAAGRPGTGEHVPGSEMPVPDYTSIHIVQSPQGLILNATALPLPDKEFLSSCGYVSTDQLNKIMP |
| **5T4D (E** | High (B | - | Q13563 | PKD2 | Q13563 | | A | B | 6 | K.{3}GF.{2}M.{2}II.{2}AY.QL.YL.{38}L.P.{2}F.{3}VF.{3}FI.{ | N.{2}A.{2}VF.{2}W.{3}F.{2}I.{27}F.{67}L.{3}L.{2}IN | 25 | 12 | 87 | 122 | MVNSSRVQPQQPGDAKRPPAPRAPDPGRLMAGCAAVGASLAAPGGLCEQRGLEIEMQRIRQAAARDPPAGAAASPSPPLSSCSRQAWSRDNPGFEAEEEEEEVEGEEGGMVVEMDVEWRP | MVNSSRVQPQQPGDAKRPPAPRAPDPGRLMAGCAAVGASLAAPGGLCEQRGLEIEMQRIRQAAARDPPAGAAASPSPPLSSCSRQAWSRDNPGFEAEEEEEEVEGEEGGMVVEMDVEWRP |

| | | | | | | | | | | | | | | | | | |
|---|---|---|---|---|---|---|---|---|---|---|---|---|---|---|---|---|---|
| **M)** | io) | | | | | | | | | 2}NM.LAI.N | | | | | | GSRRSAASSAVSSVGARSRGLGGYHGAGHPSGRRRRREDQGPPCPSPVGGGDPLHRHLPLEGQPPRVAWAERLVRGLRGLWGTRLMEESSTNREKYLKSVLRELVTYLLFLIVLCILTYGMMSSNVYYYTRMMSQLFLDTPVSKTEKTNFKTLSSMEDFWKFTEGSLLDGLYWKMQPSNQTEADNRSFIFYENLLLGVPRIRQLRVRNGSCSIPQDLRDEIKECYDVYSVSSEDRAPFGPRNGTAWIYTSEKDLNGSSHWGIIATYSGAGYYLDLSRTREETAAQVASLKKNVWLDRGTRATFIDFSVYNANINLFCVVRLLVEFPATGGVIPSWQFQPLKLIRYVTTFDFFLAACEIIFCFFIFYYVVEEILEIRIHKLHYFRSFWNCLDVVIVVLSVVAIGINIYRTSNVEVLLQFLEDQNTFPNFEHLAYWQIQFNNIAAVTVFFVWIKLFKFINFNRTMSQLSTTMSRCAKDLFGFAIMFFIIFLAYAQLAYLVFGTQVDDFSTFQECIFTQFRIILGDINFAEIEEANRVLGPIYFTTFVFFMFFILLNMFLAIINDTYSEVKSDLAQQKAEMELSDLIRKGYHKALVKLKLKKNTVDDISESLRQGGGKLNFDELRQDLKGKGHTDAEIEAIFTKYDQDGDQELTEHEHQQMRDDLEKEREDLDLDHSSLPRPMSSRSFPRSLDDSEEDDDEDSGHSSRRRGSISSGVSYEEFQVLVRRVDRMEHSIGSIVSKIDAVIVKLEIMERAKLKRREVLGRLLDGVAEDERLGRDSEIHREQMERLVREELERWESDDAASQISHGLGTPVGLNGQPRPRSSRPSSSQSTEGMEGAGGNGSSNVHV | GSRRSAASSAVSSVGARSRGLGGYHGAGHPSGRRRRREDQGPPCPSPVGGGDPLHRHLPLEGQPPRVAWAERLVRGLRGLWGTRLMEESSTNREKYLKSVLRELVTYLLFLIVLCILTYGMMSSNVYYYTRMMSQLFLDTPVSKTEKTNFKTLSSMEDFWKFTEGSLLDGLYWKMQPSNQTEADNRSFIFYENLLLGVPRIRQLRVRNGSCSIPQDLRDEIKECYDVYSVSSEDRAPFGPRNGTAWIYTSEKDLNGSSHWGIIATYSGAGYYLDLSRTREETAAQVASLKKNVWLDRGTRATFIDFSVYNANINLFCVVRLLVEFPATGGVIPSWQFQPLKLIRYVTTFDFFLAACEIIFCFFIFYYVVEEILEIRIHKLHYFRSFWNCLDVVIVVLSVVAIGINIYRTSNVEVLLQFLEDQNTFPNFEHLAYWQIQFNNIAAVTVFFVWIKLFKFINFNRTMSQLSTTMSRCAKDLFGFAIMFFIIFLAYAQLAYLVFGTQVDDFSTFQECIFTQFRIILGDINFAEIEEANRVLGPIYFTTFVFFMFFILLNMFLAIINDTYSEVKSDLAQQKAEMELSDLIRKGYHKALVKLKLKKNTVDDISESLRQGGGKLNFDELRQDLKGKGHTDAEIEAIFTKYDQDGDQELTEHEHQQMRDDLEKEREDLDLDHSSLPRPMSSRSFPRSLDDSEEDDDEDSGHSSRRRGSISSGVSYEEFQVLVRRVDRMEHSIGSIVSKIDAVIVKLEIMERAKLKRREVLGRLLDGVAEDERLGRDSEIHREQMERLVREELERWESDDAASQISHGLGTPVGLNGQPRPRSSRPSSSQSTEGMEGAGGNGSSNVHV |
| **5TIN** | High | - | O75311 | GLRA3 | O75311 | | A | B | 4 | A.{3}V.{3}I.{3}L.{6}S.{2}R.{19}L.{3}F.{2}L | IQ.{6}LI.IL.{5}WI.{6}A.{2}AL.{2}T.{3}T.{64}R | 9 | 14 | 50 | 103 | MAHVRHFRTLVSGFYFWEAALLLSLVATKETDSARSRSAPMSPSDFLDKLMGRTSGYDARIRPNFKGPPVNVTCNIFINSFGSIAETTMDYRVNIFLRQKWNDPRLAY | MAHVRHFRTLVSGFYFWEAALLLSLVATKETDSARSRSAPMSPSDFLDKLMGRTSGYDARIRPNFKGPPVNVTCNIFINSFGSIAETTMDYRVNIFLRQKWNDPRLAY |

| | | | | | | | | | | | | | | | | | |
|---|---|---|---|---|---|---|---|---|---|---|---|---|---|---|---|---|---|
| **(X-ray)** | (Bio) | | | | | | | | | | | | | | | SEYPDDSLDLDPSMLDSIWKPDLFFANEKGANFHEVTTDNKLLRIFKNGNVLYSIRLTLTLSCPMDLKNFPMDVQTCIMQLESFGYTMNDLIFEWQDEAPVQVAEGLTLPQFLLKEEKDLRYCTKHYNTGKFTCIEVRFHLERQMGYYLIQMYIPSLLIVILSWVSFWINMDAAPARVALGITTVLTMTTQSSGSRASLPKVSYVKAIDIWMAVCLLFVFSALLEYAAVNFVSRQHKELLRFRRKRKNKTEAFALEKFYRFSDMDDEVRESRFSFTAYGMGPCLQAKDGMTPKGPNHPVQVMPKSPDEMRKVFIDRAKKIDTISRACFPLAFLIFNIFYWVIYKILRHEDIHQQQD | SEYPDDSLDLDPSMLDSIWKPDLFFANEKGANFHEVTTDNKLLRIFKNGNVLYSIRLTLTLSCPMDLKNFPMDVQTCIMQLESFGYTMNDLIFEWQDEAPVQVAEGLTLPQFLLKEEKDLRYCTKHYNTGKFTCIEVRFHLERQMGYYLIQMYIPSLLIVILSWVSFWINMDAAPARVALGITTVLTMTTQSSGSRASLPKVSYVKAIDIWMAVCLLFVFSALLEYAAVNFVSRQHKELLRFRRKRKNKTEAFALEKFYRFSDMDDEVRESRFSFTAYGMGPCLQAKDGMTPKGPNHPVQVMPKSPDEMRKVFIDRAKKIDTISRACFPLAFLIFNIFYWVIYKILRHEDIHQQQD |
| **5U6O (EM)** | High (Bio) | - | O60741 | HCN1 | O60741 | | A | B | 6 | Y.{2}FV.{3}T.{2}IQ | R.{74}LW.{2}M.{2}MI.{2}A.{2}YAM.VGH.T.{3}Q | 6 | 15 | 13 | 102 | MEGGGKPNSSSNSRDDGNSVFPAKASATGAGPAAAEKRLGTPPGGGGAGAKEHGNSVCFKVDGGGGGGGGGGGGGEEPAGGFEDAEGPRRQYGFMQRQFTSMLQPGVNKFSLRMFGSQKAVEKEQERVKTAGFWIIHPYSDFRFYWDLIMLIMMVGNLVIIPVGITFFTEQTTTPWIIFNVASDTVFLLDLIMNFRTGTVNEDSSEIILDPKVIKMNYLKSWFVVDFISSIPVDYIFLIVEKGMDSEVYKTARALRIVRFTKILSLLRLLRLSRLIRYIHQWEEIFHMTYDLASAVVRIFNLIGMMLLLCHWDGCLQFLVPLLQDFPPDCWVSLNEMVNDSWGKQYSYALFKAMSHMLCIGYGAQAPVSMSDLWITMLSMIVGATCYAMFVGHATALIQSLDSSRRQYQEKYKQVEQYMSFHKLPADMRQKIHDYYEHRYQGKIFDEENILNELNDPLREEIVNFNCRKLVATMPLFANADPNFVTAMLSKLRFEVFQPGDYIIREGAVGKKMYFIQHGVAGVITKSSKEMKLTDGSYFGEICLLTKGRRTASVRADTYCRLYSLSVDNFNEVLEEYPMMRRAFETV | MEGGGKPNSSSNSRDDGNSVFPAKASATGAGPAAAEKRLGTPPGGGGAGAKEHGNSVCFKVDGGGGGGGGGGGGGEEPAGGFEDAEGPRRQYGFMQRQFTSMLQPGVNKFSLRMFGSQKAVEKEQERVKTAGFWIIHPYSDFRFYWDLIMLIMMVGNLVIIPVGITFFTEQTTTPWIIFNVASDTVFLLDLIMNFRTGTVNEDSSEIILDPKVIKMNYLKSWFVVDFISSIPVDYIFLIVEKGMDSEVYKTARALRIVRFTKILSLLRLLRLSRLIRYIHQWEEIFHMTYDLASAVVRIFNLIGMMLLLCHWDGCLQFLVPLLQDFPPDCWVSLNEMVNDSWGKQYSYALFKAMSHMLCIGYGAQAPVSMSDLWITMLSMIVGATCYAMFVGHATALIQSLDSSRRQYQEKYKQVEQYMSFHKLPADMRQKIHDYYEHRYQGKIFDEENILNELNDPLREEIVNFNCRKLVATMPLFANADPNFVTAMLSKLRFEVFQPGDYIIREGAVGKKMYFIQHGVAGVITKSSKEMKLTDGSYFGEICLLTKGRRTASVRADTYCRLYSLSVDNFNEVLEEYPMMRRAFETV |

| | | | | | | | | | | | | | | | | | |
|---|---|---|---|---|---|---|---|---|---|---|---|---|---|---|---|---|---|
| | | | | | | | | | | | | | | | | AIDRLDRIGKKNSILLQKFQKDLNTGVFNNQENEILKQIVKHDREMVQAIAPINYPQMTTLNSTSSTTTPTSRMRTQSPPVYTATSLSHSNLHSPSPSTQTPQPSAILSPCSYTTAVCSPPVQSPLAARTFHYASPTASQLSLMQQQPQQQVQQSQPPQTQPQQPSPQPQTPGSSTPKNEVHKSTQALHNTNLTREVRPLSASQPSLPHEVSTLISRPHPTVGESLASIPQPVTAVPGTGLQAGGRSTVPQRVTLFRQMSSGAIPPNRGVPPAPPPPAAALPRESSSVLNTDPDAEKPRFASNL | AIDRLDRIGKKNSILLQKFQKDLNTGVFNNQENEILKQIVKHDREMVQAIAPINYPQMTTLNSTSSTTTPTSRMRTQSPPVYTATSLSHSNLHSPSPSTQTPQPSAILSPCSYTTAVCSPPVQSPLAARTFHYASPTASQLSLMQQQPQQQVQQSQPPQTQPQQPSPQPQTPGSSTPKNEVHKSTQALHNTNLTREVRPLSASQPSLPHEVSTLISRPHPTVGESLASIPQPVTAVPGTGLQAGGRSTVPQRVTLFRQMSSGAIPPNRGVPPAPPPPAAALPRESSSVLNTDPDAEKPRFASNL |
| **5WJ9 (EM)** | High (Bio) | - | Q9GZU1 | MCOLN1 | Q9GZU1 | | A | B | 6 | F.{2}C.{2}VI.{2}GY.FC.WIV.{2}P.{39}LVW.F.Q.{3}Y.{2}I.{3}IYM.{2}S | T.{2}LI.{302}D.{2}SI.{2}GT.{2}L.{2}WV.VI.{2}L | 23 | 14 | 84 | 329 | MTAPAGPRGSETERLLTPNPGYGTQAGPSPAPPTPPEEEDLRRRLKYFFMSPCDKFRAKGRKPCKLMLQVVKILVVTVQLILFGLSNQLAVTFREENTIAFRHLFLLGYSDGADDTFAAYTREQLYQAIFHAVDQYLALPDVSLGRYAYVRGGGDPWTNGSGLALCQRYYHRGHVDPANDTFDIDPMVVTDCIQVDPPERPPPPPSDDLTLLESSSSYKNLTLKFHKLVNVTIHFRLKTINLQSLINNEIPDCYTFSVLITFDNKAHSGRIPISLETQAHIQECKHPSVFQHGDNSFRLLFDVVVILTCSLSFLLCARSLLRGFLLQNEFVGFMWRQRGRVISLWERLEFVNGWYILLVTSDVLTISGTIMKIGIEAKNLASYDVCSILLGTSTLLVWVGVIRYLTFFHNYNILIATLRVALPSVMRFCCCVAVIYLGYCFCGWIVLGPYHVKFRSLSMVSECLFSLINGDDMFVTFAAMQAQQGRSSLVWLFSQLYLYSFISLFIYMVLSLFIALITGAYDTIKHPGGAGAEESELQAYIAQCQDSPTSGKFRRGSGSACSLLCCCGRDPSEEHSLLVN | MTAPAGPRGSETERLLTPNPGYGTQAGPSPAPPTPPEEEDLRRRLKYFFMSPCDKFRAKGRKPCKLMLQVVKILVVTVQLILFGLSNQLAVTFREENTIAFRHLFLLGYSDGADDTFAAYTREQLYQAIFHAVDQYLALPDVSLGRYAYVRGGGDPWTNGSGLALCQRYYHRGHVDPANDTFDIDPMVVTDCIQVDPPERPPPPPSDDLTLLESSSSYKNLTLKFHKLVNVTIHFRLKTINLQSLINNEIPDCYTFSVLITFDNKAHSGRIPISLETQAHIQECKHPSVFQHGDNSFRLLFDVVVILTCSLSFLLCARSLLRGFLLQNEFVGFMWRQRGRVISLWERLEFVNGWYILLVTSDVLTISGTIMKIGIEAKNLASYDVCSILLGTSTLLVWVGVIRYLTFFHNYNILIATLRVALPSVMRFCCCVAVIYLGYCFCGWIVLGPYHVKFRSLSMVSECLFSLINGDDMFVTFAAMQAQQGRSSLVWLFSQLYLYSFISLFIYMVLSLFIALITGAYDTIKHPGGAGAEESELQAYIAQCQDSPTSGKFRRGSGSACSLLCCCGRDPSEEHSLLVN |
| **5Z1W** | High | - | A2A2 | Pkd2 | A2A2 | | A | C | 6 | Y.{2}T.{317}N.{2}A.{2}L | M.{6}AY.QL.{2}LL.G.{32}NR.{3}P | 8 | 14 | 336 | 63 | MNSMESPKNQELQTLGNRAWDNPAYSDPPSPNRTLRICTVSSVALPETQPKKPEVRCQEKTQRTLVSSCCL | MNSMESPKNQELQTLGNRAWDNPAYSDPPSPNRTLRICTVSSVALPETQPKKPEVRCQEKTQRTLVSSCCL |

| | | | | | | | | | | | | | | | | | |
|---|---|---|---|---|---|---|---|---|---|---|---|---|---|---|---|---|---|
| **(EM)** | (Bio) | | 59 | l1 | 59 | | | | | F.{2}W.{3}F | .{2}FV.{2}V | | | | | HICRSIRGLWGTTLTENTAENRELYVKTTLRELVVYIVFLVDICLLTYGMTSSSAYYYTKVMSELFLHTPSDSGVSFQTISSMSDFWDFAQGPLLDSLYWTKWYNNQSLGRGSHSFIYYENLLLGAPRLRQLRVRNDSCVVHEDFREDILNCYDVYSPDKEDQLPFGPQNGTAWTYHSQNELGGSSHWGRLTSYSGGGYYLDLPGSRQASAEALQGLQEGLWLDRGTRVVFIDFSVYNANINLFCILRLVVEFPATGGTIPSWQIRTVKLIRYVNNWDFFIVGCEVVFCVFIFYYVVEEILEIHLHRLRYLSSVWNILDLVVILLSIVAVGFHIFRTLEVNRLMGKLLQQPDTYADFEFLAFWQTQYNNMNAVNLFFAWIKIFKYISFNKTMTQLSSTLARCAKDILGFAIMFFIVFFAYAQLGYLLFGTQVENFSTFVKCIFTQFRIILGDFDYNAIDNANRILGPVYFVTYVFFVFFVLLNMFLAIINDTYSEVKEELAGQKDQLQLSDFLKQSYNKTLLRLRLRKERVSDVQKVLKGGEPEIQFEDFTSTLRELGHEEHEITAAFTRFDQDGDHILDEEEQEQMRQGLEEERVTLNAEIENLGRSVGHSPPGELGAEAARGQSWVSGEEFDMLTRRVLQLQCVLEGVVSQIDAVGSKLKMLERKGELAPSPGMGEPAVWENLYNPS | HICRSIRGLWGTTLTENTAENRELYVKTTLRELVVYIVFLVDICLLTYGMTSSSAYYYTKVMSELFLHTPSDSGVSFQTISSMSDFWDFAQGPLLDSLYWTKWYNNQSLGRGSHSFIYYENLLLGAPRLRQLRVRNDSCVVHEDFREDILNCYDVYSPDKEDQLPFGPQNGTAWTYHSQNELGGSSHWGRLTSYSGGGYYLDLPGSRQASAEALQGLQEGLWLDRGTRVVFIDFSVYNANINLFCILRLVVEFPATGGTIPSWQIRTVKLIRYVNNWDFFIVGCEVVFCVFIFYYVVEEILEIHLHRLRYLSSVWNILDLVVILLSIVAVGFHIFRTLEVNRLMGKLLQQPDTYADFEFLAFWQTQYNNMNAVNLFFAWIKIFKYISFNKTMTQLSSTLARCAKDILGFAIMFFIVFFAYAQLGYLLFGTQVENFSTFVKCIFTQFRIILGDFDYNAIDNANRILGPVYFVTYVFFVFFVLLNMFLAIINDTYSEVKEELAGQKDQLQLSDFLKQSYNKTLLRLRLRKERVSDVQKVLKGGEPEIQFEDFTSTLRELGHEEHEITAAFTRFDQDGDHILDEEEQEQMRQGLEEERVTLNAEIENLGRSVGHSPPGELGAEAARGQSWVSGEEFDMLTRRVLQLQCVLEGVVSQIDAVGSKLKMLERKGELAPSPGMGEPAVWENLYNPS |
| **5Z96 (EM)** | High (Bio) | - | Q9QUQ5 | Trpc4 | Q9QUQ5 | | A | B | 6 | KF.FI.{2}L.{3}AF.NG.NQL.F.{52}F.{2}FV.A.MFG.{2}NV.{2}L.{4}NM | LL.AS.{90}LV.EA.FAI.NI.{2}S.{2}L.{3}F.{19}I.{100}I | 25 | 18 | 98 | 238 | MAQFYYKRNVNAPYRDRIPLRIVRAESELSPSEKAYLNAVEKGDYASVKKSLEEAEIYFKININCIDPLGRTALLIAIENENLELIELLLSFNVYVGDALLHAIRKEVVGAVELLLNHKKPSGEKQVPPILLDKQFSEFTPDITPIILAAHTNNYEIIKLLVQKGVSVPRPHEVRCNCVECVSSSDVDSLRHSRSRLNIYKALASPSLIALSSEDPFLTAFQLSWELQELSKVENEFKSEYEELSRQCKQFAKDLLDQTRSSRELEIILNYRDDNSLIEEQSGNDL | MAQFYYKRNVNAPYRDRIPLRIVRAESELSPSEKAYLNAVEKGDYASVKKSLEEAEIYFKININCIDPLGRTALLIAIENENLELIELLLSFNVYVGDALLHAIRKEVVGAVELLLNHKKPSGEKQVPPILLDKQFSEFTPDITPIILAAHTNNYEIIKLLVQKGVSVPRPHEVRCNCVECVSSSDVDSLRHSRSRLNIYKALASPSLIALSSEDPFLTAFQLSWELQELSKVENEFKSEYEELSRQCKQFAKDLLDQTRSSRELEIILNYRDDNSLIEEQSGNDL |

| | | | | | | | | | | | | | | | | | |
|---|---|---|---|---|---|---|---|---|---|---|---|---|---|---|---|---|---|
| | | | | | | | | | | | | | | | | ARLKLAIKYRQKEFVAQPNCQQLLASRWYDEFPGWRRRHWAVKMVTCFIIGLLFPVFSVCYLIAPKSPLGLFIRKPFIKFICHTASYLTFLFLLLLASQHIDRSDLNRQGPPPTIVEWMILPWVLGFIWGEIKQMWDGGLQDYIHDWWNLMDFVMNSLYLATISLKIVAFVKYSALNPRESWDMWHPTLVAEALFAIANIFSSLRLISLFTANSHLGPLQISLGRMLLDILKFLFIYCLVLLAFANGLNQLYFYYEETKGLSCKGIRCEKQNNAFSTLFETLQSLFWSIFGLINLYVTNVKAQHEFTEFVGATMFGTYNVISLVVLLNMLIAMMNNSYQLIADHADIEWKFARTKLWMSYFEEGGTLPTPFNVIPSPKSLWYLVKWIWTHLCKKKMRRKPESFGTIGRRAADNLRRHHQYQEVMRNLVKRYVAAMIREAKTEEGLTEENVKELKQDISSFRFEVLGLLRGSKLSTIQSANAASSADSDEKSQSEGNGKDKRKNLSLFDLTTLIHPRSAAIASERHNLSNGSALVVQEPPREKQRKVNFVADIKNFGLFHRRSKQNAAEQNANQIFSVSEEITRQQAAGALERNIELESKGLASRGDRSIPGLNEQCVLVDHRERNTDTLGLQVGKRVCSTFKSEKVVVEDTVPIIPKEKHAHEEDSSIDYDLSPTDTAAHEDYVVTTRL | ARLKLAIKYRQKEFVAQPNCQQLLASRWYDEFPGWRRRHWAVKMVTCFIIGLLFPVFSVCYLIAPKSPLGLFIRKPFIKFICHTASYLTFLFLLLLASQHIDRSDLNRQGPPPTIVEWMILPWVLGFIWGEIKQMWDGGLQDYIHDWWNLMDFVMNSLYLATISLKIVAFVKYSALNPRESWDMWHPTLVAEALFAIANIFSSLRLISLFTANSHLGPLQISLGRMLLDILKFLFIYCLVLLAFANGLNQLYFYYEETKGLSCKGIRCEKQNNAFSTLFETLQSLFWSIFGLINLYVTNVKAQHEFTEFVGATMFGTYNVISLVVLLNMLIAMMNNSYQLIADHADIEWKFARTKLWMSYFEEGGTLPTPFNVIPSPKSLWYLVKWIWTHLCKKKMRRKPESFGTIGRRAADNLRRHHQYQEVMRNLVKRYVAAMIREAKTEEGLTEENVKELKQDISSFRFEVLGLLRGSKLSTIQSANAASSADSDEKSQSEGNGKDKRKNLSLFDLTTLIHPRSAAIASERHNLSNGSALVVQEPPREKQRKVNFVADIKNFGLFHRRSKQNAAEQNANQIFSVSEEITRQQAAGALERNIELESKGLASRGDRSIPGLNEQCVLVDHRERNTDTLGLQVGKRVCSTFKSEKVVVEDTVPIIPKEKHAHEEDSSIDYDLSPTDTAAHEDYVVTTRL |
| **6BCO (EM)** | High (Bio) | - | Q7TN37 | Trpm4 | Q7TN37 | | A | B | 6 | KD.{2}F.{2}FF.{2}VW.VA.{2}VA.EGI.{67}L.V.L.I.{2}LL.{2}NIL.{2}NL.I | L.{86}D.{2}RT.{2}CL.FM.{2}TL.{2}L.{2}F.{19}V.{2}F.{105}LI | 26 | 16 | 113 | 239 | MVGPEKEQSWIPKIFRKKVCTTFIVDLSDDAGGTLCQCGQPRDAHPSVAVEDAFGAAVVTEWNSDEHTTEKPTDAYGDLDFTYSGRKHSNFLRLSDRTDPATVYSLVTRSWGFRAPNLVVSVLGGSGGPVLQTWLQDLLRRGLVRAAQSTGAWIVTGGLHTGIGRHVGVAVRDHQTASTGSSKVVAMGVAPWGVVRNRDMLINPKGSFPARYRWRGDPEDGVEFPLDYNYSAFFLVDDGTYGRLGGENRFRLRFESYVAQQK | MVGPEKEQSWIPKIFRKKVCTTFIVDLSDDAGGTLCQCGQPRDAHPSVAVEDAFGAAVVTEWNSDEHTTEKPTDAYGDLDFTYSGRKHSNFLRLSDRTDPATVYSLVTRSWGFRAPNLVVSVLGGSGGPVLQTWLQDLLRRGLVRAAQSTGAWIVTGGLHTGIGRHVGVAVRDHQTASTGSSKVVAMGVAPWGVVRNRDMLINPKGSFPARYRWRGDPEDGVEFPLDYNYSAFFLVDDGTYGRLGGENRFRLRFESYVAQQK |

| | | | | | | | | | | | | | | | | | |
|---|---|---|---|---|---|---|---|---|---|---|---|---|---|---|---|---|---|
| | | | | | | | | | | | | | | | | TGVGGTGIDIPVLLLLIDG<br>DEKMLKRIEDATQAQLPC<br>LLVAGSGGAADCLVETLE<br>DTLAPGSGGLRRGEARDR<br>IRRYFPKGDPEVLQAQVE<br>RIMTRKELLTVYSSEDGSE<br>EFETIVLRALVKACGSSEA<br>SAYLDELRLAVAWNRVDI<br>AQSELFRGDIQWRSFHLE<br>ASLMDALLNDRPEFVRLLI<br>SHGLSLGHFLTPVRLAQLY<br>SAVSPNSLIRNLLDQASH<br>ASSSKSPPVNGTVELRPP<br>NVGQVLRTLLGETCAPRY<br>PARNTRDSYLGQDHREN<br>DSLLMDWANKQPSTDAS<br>FEQAPWSDLLIWALLLNR<br>AQMAIYFWEKGSNSVAS<br>ALGACLLLRVMARLESEA<br>EEAARRKDLAATFESMSV<br>DLFGECYHNSEERAARLLL<br>RRCPLWGEATCLQLAMQ<br>ADARAFFAQDGVQSLLT<br>QKWWGEMDSTTPIWAL<br>LLAFFCPPLIYTNLIVFRKS<br>EEEPTQKDLDFDMDSSIN<br>GAGPPGTVEPSAKVALER<br>RQRRRPGRALCCGKFSKR<br>WSDFWGAPVTAFLGNV<br>VSYLLFLLLFAHVLLVDFQ<br>PTKPSVSELLLYFWAFTLL<br>CEELRQGLGGGWGSLAS<br>GGRGPDRAPLRHRLHLYL<br>SDTWNQCDLLALTCFLLG<br>VGCRLTPGLFDLGRTVLCL<br>DFMIFTLRLLHIFTVNKQL<br>GPKIVIVSKMMKDVFFFL<br>FFLCVWLVAYGVATEGIL<br>RPQDRSLPSILRRVFYRPY<br>LQIFGQIPQEEMDVALMI<br>PGNCSMERGSWAHPEG<br>PVAGSCVSQYANWLVVL<br>LLIVFLLVANILLLNLLIAM<br>FSYTFSKVHGNSDLYWKA<br>QRYSLIREFHSRPALAPPLI<br>IISHVRLLIKWLRRCRRCR<br>RANLPASPVFEHFRVCLS<br>KEAERKLLTWESVHKENF<br>LLAQARDKRDSDSERLKR<br>TSQKVDTALKQLGQIREY<br>DRRLRGLEREVQHCSRVL<br>TWMAEALSHSALLPPGA<br>PPPPSPTGSKD | TGVGGTGIDIPVLLLLIDG<br>DEKMLKRIEDATQAQLPC<br>LLVAGSGGAADCLVETLE<br>DTLAPGSGGLRRGEARDR<br>IRRYFPKGDPEVLQAQVE<br>RIMTRKELLTVYSSEDGSE<br>EFETIVLRALVKACGSSEA<br>SAYLDELRLAVAWNRVDI<br>AQSELFRGDIQWRSFHLE<br>ASLMDALLNDRPEFVRLLI<br>SHGLSLGHFLTPVRLAQLY<br>SAVSPNSLIRNLLDQASH<br>ASSSKSPPVNGTVELRPP<br>NVGQVLRTLLGETCAPRY<br>PARNTRDSYLGQDHREN<br>DSLLMDWANKQPSTDAS<br>FEQAPWSDLLIWALLLNR<br>AQMAIYFWEKGSNSVAS<br>ALGACLLLRVMARLESEA<br>EEAARRKDLAATFESMSV<br>DLFGECYHNSEERAARLLL<br>RRCPLWGEATCLQLAMQ<br>ADARAFFAQDGVQSLLT<br>QKWWGEMDSTTPIWAL<br>LLAFFCPPLIYTNLIVFRKS<br>EEEPTQKDLDFDMDSSIN<br>GAGPPGTVEPSAKVALER<br>RQRRRPGRALCCGKFSKR<br>WSDFWGAPVTAFLGNV<br>VSYLLFLLLFAHVLLVDFQ<br>PTKPSVSELLLYFWAFTLL<br>CEELRQGLGGGWGSLAS<br>GGRGPDRAPLRHRLHLYL<br>SDTWNQCDLLALTCFLLG<br>VGCRLTPGLFDLGRTVLCL<br>DFMIFTLRLLHIFTVNKQL<br>GPKIVIVSKMMKDVFFFL<br>FFLCVWLVAYGVATEGIL<br>RPQDRSLPSILRRVFYRPY<br>LQIFGQIPQEEMDVALMI<br>PGNCSMERGSWAHPEG<br>PVAGSCVSQYANWLVVL<br>LLIVFLLVANILLLNLLIAM<br>FSYTFSKVHGNSDLYWKA<br>QRYSLIREFHSRPALAPPLI<br>IISHVRLLIKWLRRCRRCR<br>RANLPASPVFEHFRVCLS<br>KEAERKLLTWESVHKENF<br>LLAQARDKRDSDSERLKR<br>TSQKVDTALKQLGQIREY<br>DRRLRGLEREVQHCSRVL<br>TWMAEALSHSALLPPGA<br>PPPPSPTGSKD |

| | | | | | | | | | | | | | | | | | |
|---|---|---|---|---|---|---|---|---|---|---|---|---|---|---|---|---|---|
| **6BQV (EM)** | High (Bio) | - | Q8TD43 | TRPM4 | Q8TD43 | | A | B | 6 | D.{2}F.{2}FF.{2}VW.VA.{2}VA.EGL.{67}L.V.LLV.{2}LL.{2}NIL.{2}NL.IAM | S.{3}L.{87}L.RT.{2}CI.FM.{2}T.{2}LL.{2}F.{19}V.{2}FL.{104}LI.{2}FS | 28 | 20 | 114 | 247 | MVVPEKEQSWIPKIFKKKTCTTFIVDSTDPGGTLCQCGRPRTAHPAVAMEDAFGAAVVTVWDSDAHTTEKPTDAYGELDFTGAGRKHSNFLRLSDRTDPAAVYSLVTRTWGFRAPNLVVSVLGGSGGPVLQTWLQDLLRRGLVRAAQSTGAWIVTGGLHTGIGRHVGVAVRDHQMASTGGTKVVAMGVAPWGVVRNRDTLINPKGSFPARYRWRGDPEDGVQFPLDYNYSAFFLVDDGTHGCLGGENRFRLRLESYISQQKTGVGGTGIDIPVLLLLIDGDEKMLTRIENATQAQLPCLLVAGSGGAADCLAETLEDTLAPGSGGARQGEARDRIRRFFPKGDLEVLQAQVERIMTRKELLTVYSSEDGSEEFETIVLKALVKACGSSEASAYLDELRLAVAWNRVDIAQSELFRGDIQWRSFHLEASLMDALLNDRPEFVRLLISHGLSLGHFLTPMRLAQLYSAAPSNSLIRNLLDQASHSAGTKAPALKGGAAELRPPDVGHVLRMLLGKMCAPRYPSGGAWDPHPGQGFGESMYLLSDKATSPLSLDAGLGQAPWSDLLLWALLLNRAQMAMYFWEMGSNAVSSALGACLLLRVMARLEPDAEEAARRKDLAFKFEGMGVDLFGECYRSSEVRAARLLLRRCPLWGDATCLQLAMQADARAFFAQDGVQSLLTQKWWGDMASTTPIWALVLAFFCPPLIYTRLITFRKSEEEPTREELEFDMDSVINGEGPVGTADPAEKTPLGVPRQSGRPGCCGGRCGGRRCLRRWFHFWGAPVTIFMGNVVSYLLFLLLFSRVLLVDFQPAPPGSLELLLYFWAFTLLCEELRQGLSGGGGSLASGGPGPGHASLSQRLRLYLADSWNQCDLVALTCFLLGVGCRLTPGLYHLGRTVLCIDFMVFTVRLLHIFTVNKQLGPKIVIVSKMMKDVFFFLFFLGVWLVAY | MVVPEKEQSWIPKIFKKKTCTTFIVDSTDPGGTLCQCGRPRTAHPAVAMEDAFGAAVVTVWDSDAHTTEKPTDAYGELDFTGAGRKHSNFLRLSDRTDPAAVYSLVTRTWGFRAPNLVVSVLGGSGGPVLQTWLQDLLRRGLVRAAQSTGAWIVTGGLHTGIGRHVGVAVRDHQMASTGGTKVVAMGVAPWGVVRNRDTLINPKGSFPARYRWRGDPEDGVQFPLDYNYSAFFLVDDGTHGCLGGENRFRLRLESYISQQKTGVGGTGIDIPVLLLLIDGDEKMLTRIENATQAQLPCLLVAGSGGAADCLAETLEDTLAPGSGGARQGEARDRIRRFFPKGDLEVLQAQVERIMTRKELLTVYSSEDGSEEFETIVLKALVKACGSSEASAYLDELRLAVAWNRVDIAQSELFRGDIQWRSFHLEASLMDALLNDRPEFVRLLISHGLSLGHFLTPMRLAQLYSAAPSNSLIRNLLDQASHSAGTKAPALKGGAAELRPPDVGHVLRMLLGKMCAPRYPSGGAWDPHPGQGFGESMYLLSDKATSPLSLDAGLGQAPWSDLLLWALLLNRAQMAMYFWEMGSNAVSSALGACLLLRVMARLEPDAEEAARRKDLAFKFEGMGVDLFGECYRSSEVRAARLLLRRCPLWGDATCLQLAMQADARAFFAQDGVQSLLTQKWWGDMASTTPIWALVLAFFCPPLIYTRLITFRKSEEEPTREELEFDMDSVINGEGPVGTADPAEKTPLGVPRQSGRPGCCGGRCGGRRCLRRWFHFWGAPVTIFMGNVVSYLLFLLLFSRVLLVDFQPAPPGSLELLLYFWAFTLLCEELRQGLSGGGGSLASGGPGPGHASLSQRLRLYLADSWNQCDLVALTCFLLGVGCRLTPGLYHLGRTVLCIDFMVFTVRLLHIFTVNKQLGPKIVIVSKMMKDVFFFLFFLGVWLVAY |

| | | | | | | | | | | | | | | | | | |
|---|---|---|---|---|---|---|---|---|---|---|---|---|---|---|---|---|---|
| | | | | | | | | | | | | | | | | GVATEGLLRPRDSDFPSILRRVFYRPYLQIFGQIPQEDMDVALMEHSNCSSEPGFWAHPPGAQAGTCVSQYANWLVVLLLVIFLLVANILLVNLLIAMFSYTFGKVQGNSDLYWKAQRYRLIREFHSRPALAPPFIVISHLRLLLRQLCRRPRSPQPSSPALEHFRVYLSKEAERKLLTWESVHKENFLLARARDKRESDSERLKRTSQKVDLALKQLGHIREYEQRLKVLEREVQQCSRVLGWVAEALSRSALLPPGGPPPPDLPGSKD | GVATEGLLRPRDSDFPSILRRVFYRPYLQIFGQIPQEDMDVALMEHSNCSSEPGFWAHPPGAQAGTCVSQYANWLVVLLLVIFLLVANILLVNLLIAMFSYTFGKVQGNSDLYWKAQRYRLIREFHSRPALAPPFIVISHLRLLLRQLCRRPRSPQPSSPALEHFRVYLSKEAERKLLTWESVHKENFLLARARDKRESDSERLKRTSQKVDLALKQLGHIREYEQRLKVLEREVQQCSRVLGWVAEALSRSALLPPGGPPPPDLPGSKD |
| **6CQ9 (X-ray)** | High (Bio) | - | P97438 | Kcnk2 | P97438 | | A | B | 4 | KW.{2}V.{3}F.{2}V.{2}YLI.GA.VF.{87}EGGKIFC.{2}YALLG.PL.{2}FL.{2}G.{2}DQ.{94}W.{13}L.{2}IG.{3}R | W.{2}V.{2}IF.{2}V.{2}YLI.GA.VFK.{87}GGK.FC.{2}YAL.GIPL.{2}FL.{2}G.{2}DQ.{108}L.{2}IG.{2}L | 36 | 34 | 255 | 253 | MLASASRERPGYTAGVAAPDLLDPKSAAQNSKPRLSFSSKPTVLASRVESDSAINVMKWKTVSTIFLVVVLYLIIGATVFKALEQPQEISQRTTIVIQKQTFIAQHACVNSTELDELIQQIVAAINAGIIPLGNSSNQVSHWDLGSSFFFAGTVITTIGFGNISPRTEGGKIFCIIYALLGIPLFGFLLAGVGDQLGTIFGKGIAKVEDTFIKWNVSQTKIRIISTIIFILFGCVLFVALPAVIFKHIEGWSALDAIYFVVITLTTIGFGDYVAGGSDIEYLDFYKPVVWFWILVGLAYFAAVLSMIGDWLRVISKKTKEEVGEFRAHAAEWTANVTAEFKETRRRLSVEIYDKFQRATSVKRKLSAELAGNHNQELTPCRRTLSVNHLTSEREVLPPLLKAESIYLNGLTPHCAGEDIAVIENMK | MLASASRERPGYTAGVAAPDLLDPKSAAQNSKPRLSFSSKPTVLASRVESDSAINVMKWKTVSTIFLVVVLYLIIGATVFKALEQPQEISQRTTIVIQKQTFIAQHACVNSTELDELIQQIVAAINAGIIPLGNSSNQVSHWDLGSSFFFAGTVITTIGFGNISPRTEGGKIFCIIYALLGIPLFGFLLAGVGDQLGTIFGKGIAKVEDTFIKWNVSQTKIRIISTIIFILFGCVLFVALPAVIFKHIEGWSALDAIYFVVITLTTIGFGDYVAGGSDIEYLDFYKPVVWFWILVGLAYFAAVLSMIGDWLRVISKKTKEEVGEFRAHAAEWTANVTAEFKETRRRLSVEIYDKFQRATSVKRKLSAELAGNHNQELTPCRRTLSVNHLTSEREVLPPLLKAESIYLNGLTPHCAGEDIAVIENMK |
| **6CUD (EM)** | High (Bio) | - | Q13507 | TRPC3 | Q13507 | | A | B | 6 | KF.VL.{5}FAF.{2}G.FIL.S.{41}FI.{2}I.Y.{2}YG.{2}N.{7}LN | A.{2}RFE.{126}I.{2}EGL.AI.{2}V.{2}F.{115}L | 21 | 13 | 86 | 263 | MREKGRRQAVRGPAFMFNDRGTSLTAEEERFLDAAEYGNIPVVRKMLEESKTLNVNCVDYMGQNALQLAVGNEHLEVTELLLKKENLARIGDALLLAISKGYVRIVEAILNHPGFAASKRLTLSPCEQELQDDDFYAYDEDGTRFSPDITPIILAAHCQKYEVVHMLLMKGARIERPHDYFCKCGDCMEKQRHDSFSHSRSRINAYKGLASPAYLSLSSEDPVLTALELSNELAKLANIEKEFKNDYRKLSMQCKDFVVGVLDLCRDSEEVE | MREKGRRQAVRGPAFMFNDRGTSLTAEEERFLDAAEYGNIPVVRKMLEESKTLNVNCVDYMGQNALQLAVGNEHLEVTELLLKKENLARIGDALLLAISKGYVRIVEAILNHPGFAASKRLTLSPCEQELQDDDFYAYDEDGTRFSPDITPIILAAHCQKYEVVHMLLMKGARIERPHDYFCKCGDCMEKQRHDSFSHSRSRINAYKGLASPAYLSLSSEDPVLTALELSNELAKLANIEKEFKNDYRKLSMQCKDFVVGVLDLCRDSEEVE |

| | | | | | | | | | | | | | | | | | |
|---|---|---|---|---|---|---|---|---|---|---|---|---|---|---|---|---|---|
| | | | | | | | | | | | | | | | | AILNGDLESAEPLEVHRHKASLSRVKLAIKYEVKKFVAHPNCQQQLLTIWYENLSGLREQTIAIKCLVVLVVALGLPFLAIGYWIAPCSRLGKILRSPFMKFVAHAASFIIFLGLLVFNASDRFEGITTLPNITVTDYPKQIFRVKTTQFTWTEMLIMVWVLGMMWSECKELWLEGPREYILQLWNVLDFGMLSIFIAAFTARFLAFLQATKAQQYVDSYVQESDLSEVTLPPEIQYFTYARDKWLPSDPQIISEGLYAIAVVLSFSRIAYILPANESFGPLQISLGRTVKDIFKFMVLFIMVFFAFMIGMFILYSYYLGAKVNAAFTTVEESFKTLFWSIFGLSEVTSVVLKYDHKFIENIGYVLYGIYNVTMVVVLLNMLIAMINSSYQEIEDDSDVEWKFARSKLWLSYFDDGKTLPPPFSLVPSPKSFVYFIMRIVNFPKCRRRRLQKDIEMGMGNSKSRLNLFTQSNSRVFESHSFNSILNQPTRYQQIMKRLIKRYVLKAQVDKENDEVNEGELKEIKQDISSLRYELLEDKSQATEELAILIHKLSEKLNPSMLRCE | AILNGDLESAEPLEVHRHKASLSRVKLAIKYEVKKFVAHPNCQQQLLTIWYENLSGLREQTIAIKCLVVLVVALGLPFLAIGYWIAPCSRLGKILRSPFMKFVAHAASFIIFLGLLVFNASDRFEGITTLPNITVTDYPKQIFRVKTTQFTWTEMLIMVWVLGMMWSECKELWLEGPREYILQLWNVLDFGMLSIFIAAFTARFLAFLQATKAQQYVDSYVQESDLSEVTLPPEIQYFTYARDKWLPSDPQIISEGLYAIAVVLSFSRIAYILPANESFGPLQISLGRTVKDIFKFMVLFIMVFFAFMIGMFILYSYYLGAKVNAAFTTVEESFKTLFWSIFGLSEVTSVVLKYDHKFIENIGYVLYGIYNVTMVVVLLNMLIAMINSSYQEIEDDSDVEWKFARSKLWLSYFDDGKTLPPPFSLVPSPKSFVYFIMRIVNFPKCRRRRLQKDIEMGMGNSKSRLNLFTQSNSRVFESHSFNSILNQPTRYQQIMKRLIKRYVLKAQVDKENDEVNEGELKEIKQDISSLRYELLEDKSQATEELAILIHKLSEKLNPSMLRCE |
| **6ETI (EM)** | High (Bio) | - | Q9UNQ0 | ABCG2 | Q9UNQ0 | | A | B | 6 | L.{2}VI.{2}IY.{14}G.{2}F.{115}F.{2}IF.GLLV | L.{2}VI.{2}IY.{14}G.{2}F.{115}F.{2}IF.GLLV | 14 | 14 | 152 | 152 | MSSSNVEVFIPVSQGNTNGFPATASNDLKAFTEGAVLSFHNICYRVKLKSGFLPCRKPVEKEILSNINGIMKPGLNAILGPTGGGKSSLLDVLAARKDPSGLSGDVLINGAPRPANFKCNSGYVVQDDVVMGTLTVRENLQFSAALRLATTMTNHEKNERINRVIQELGLDKVADSKVGTQFIRGVSGGERKRTSIGMELITDPSILFLDEPTTGLDSSTANAVLLLLKRMSKQGRTIIFSIHQPRYSIFKLFDSLTLLASGRLMFHGPAQEALGYFESAGYHCEAYNNPADFFLDIINGDSTAVALNREEDFKATEIIEPSKQDKPLIEKLAEIYVNSSFYKETKAELHQLSGGEKKKKITVFKEISYTTSFCHQLRWVSKRSFKNLLGNPQASIAQIIVTVVLGLVIG | MSSSNVEVFIPVSQGNTNGFPATASNDLKAFTEGAVLSFHNICYRVKLKSGFLPCRKPVEKEILSNINGIMKPGLNAILGPTGGGKSSLLDVLAARKDPSGLSGDVLINGAPRPANFKCNSGYVVQDDVVMGTLTVRENLQFSAALRLATTMTNHEKNERINRVIQELGLDKVADSKVGTQFIRGVSGGERKRTSIGMELITDPSILFLDEPTTGLDSSTANAVLLLLKRMSKQGRTIIFSIHQPRYSIFKLFDSLTLLASGRLMFHGPAQEALGYFESAGYHCEAYNNPADFFLDIINGDSTAVALNREEDFKATEIIEPSKQDKPLIEKLAEIYVNSSFYKETKAELHQLSGGEKKKKITVFKEISYTTSFCHQLRWVSKRSFKNLLGNPQASIAQIIVTVVLGLVIG |

| | | | | | | | | | | | | | | | | | |
|---|---|---|---|---|---|---|---|---|---|---|---|---|---|---|---|---|---|
| | | | | | | | | | | | | | | | | AIYFGLKNDSTGIQNRAG<br>VLFFLTTNQCFSSVSAVEL<br>FVVEKKLFIHEYISGYYRVS<br>SYFLGKLLSDLLPMRMLP<br>SIIFTCIVYFMLGLKPKADA<br>FFVMMFTLMMVAYSASS<br>MALAIAAGQSVVSVATLL<br>MTICFVFMMIFSGLLVNL<br>TTIASWLSWLQYFSIPRYG<br>FTALQHNEFLGQNFCPGL<br>NATGNNPCNYATCTGEE<br>YLVKQGIDLSPWGLWKN<br>HVALACMIVIFLTIAYLKLL<br>FLKKYS | AIYFGLKNDSTGIQNRAG<br>VLFFLTTNQCFSSVSAVEL<br>FVVEKKLFIHEYISGYYRVS<br>SYFLGKLLSDLLPMRMLP<br>SIIFTCIVYFMLGLKPKADA<br>FFVMMFTLMMVAYSASS<br>MALAIAAGQSVVSVATLL<br>MTICFVFMMIFSGLLVNL<br>TTIASWLSWLQYFSIPRYG<br>FTALQHNEFLGQNFCPGL<br>NATGNNPCNYATCTGEE<br>YLVKQGIDLSPWGLWKN<br>HVALACMIVIFLTIAYLKLL<br>FLKKYS |

Column

A: PDB accession code of validated structures of membrane protein complexes

B: Biological EPPIC assembly probability (PRODIGY Predicted Interface Biological vs Crystallographic), High or Low

C: PDB accession code for the unbound structures of the components of complexes, used for defining a benchmark of membrane protein-protein interactions (Table 1 main text)

D (F): Uniprot accession number of protein A (B)

E (G): Gene name of protein A (B when different from A)

H (I): Chain name of protein A (B)

J: number of TM helices of protomer in complex

K (L): Binding motif of protein A (B)

M (N): Number of interface contact residues of protein A (B)

O (P): Total length of matched motif, including variable residues, of protein A (B)

Q (R): Sequence of protein A (B)

Table X Paired/coupled contact motifs in which motif A is different from motif B

Deductions?

| Motif A | Motif B |
|---|---|
| A.{3}V.{3}I.{3}L.{6}S.{2}R.{19}L.{3}F.{2}L | Q.{6}LI.IL.{5}WI.{6}A.{2}AL.{2}T |
| DWN.{2}IA.{2}V.{2}LI.{24}F.{3}F.{2}A.{2}YLV.{2}FM | VI.{2}V.{3}F.WLV.LL.{2}SV.WF.{17}L.{6}SV.{2}Q.{44}IS.VFS.IN.{22}TS.{2}LT.{32}H.{3}S.{2}TF.N |
| E.{2}TL.VGF.IL.{2}SL.{2}TY | AV.{2}G.{2}FV.{2}GP.{2}AL.{2}I.{112}NI.{19}FL.{3}F |
| I.{3}L.{2}T.{6}R.{12}D.{2}MA.{6}F.{2}LL | GYY.IQ.{6}L.{2}IL.{2}V.{13}A.{3}T.{3}T.{3}Q |
| K.{3}GF.{2}M.{2}II.{2}AY.QL.YL.{38}L.P.{2}F.{3}VF.{3}FI.{2}NM.LAI.N | N.{2}A.{2}VF.{2}W.{3}F.{2}I.{27}F.{67}L.{3}L.{2}IN |
| K.{6}L.{6}FF.{2}GS.{11}LQ.{2}LA.{2}LA.{2}T.{2}QAL.PV.{40}I.{58}L | L.{86}Q.{3}V.{2}IL.{2}Q.{3}C.{2}AS.{14}S.{2}LS.TL.{2}LV.{44}L.{3}L |
| KF.FI.{2}L.{3}AF.NG.NQL.F.{52}F.{2}FV.A.MFG.{2}NV.{2}L.{4}NM | LL.AS.{90}LV.EA.FAI.NI.{2}S.{2}L.{3}F.{19}I.{100}I |
| KF.VL.{5}FAF.{2}G.FIL.S.{41}FI.{2}I.Y.{2}YG.{2}N.{7}LN | A.{2}RFE.{126}I.{2}EGL.AI.{2}V.{2}F.{115}L |
| L.{3}V.{3}T.{2}W.{81}V.{2}MF | LL.{3}VL.SI.{2}AF.{2}G |
| Q.{6}LI.IL.{5}WI.{6}A.{2}AL.{2}T | S.{3}V.{3}I.{3}L.{6}T.{19}M.{2}F.{3}F.{2}LL.{5}N |
| R.{2}FA.{2}L.{3}L.{2}FF.{2}GS.{11}LQ.{2}MA.{2}LG.{2}T.VQALG.{42}LL.{57}L | VL.{86}Q.{2}TV.{2}FL.LQ.{2}LC.{2}AS.{14}S.{2}FS.AL.{2}LL.{43}LY |
| V.{3}T.{3}V.{3}L.{6}I.{22}Y.{3}F.{2}L.{3}A | IQ.{6}M.{2}IL.{2}V.{3}L.{6}A.{3}F.{2}T.{3}T.{130}R |
| W.{2}V.{3}F.{2}M.{3}V.{3}EV.{143}E.{2}VF.{2}F | IR.{3}L.{2}IF.{2}T.{3}L |
| Y.{2}FV.{3}T.{2}IQ | R.{74}LW.{2}M.{2}MI.{2}A.{2}YAM.VGH.T.{3}Q |
| Y.{2}I.{7}L.{2}IL.{16}GL.{2}T.{2}LT.{2}T.{2}SG.R | I.{3}L.{2}T.{3}S.{2}R.{19}L.{3}F.{2}L.{3}A |

### Table S4

Set of 417 human PPIMem MPs involved in predicted complexes

| UniProKB Accession | UniProtKB Name | Mendelian Inheritance in Man Number (MIM) |
|---|---|---|
| A0PJK1 | Sodium/glucose cotransporter 5 | |
| A6NHA9 | Olfactory receptor 4C46 | |
| A6NI61 | Protein myomaker {ECO:0000250\|UniProtKB:Q9D1N4} | 254940 |
| A6NM45 | Putative claudin-24 {ECO:0000305} | |
| A8MPY1 | Gamma-aminobutyric acid receptor subunit rho-3 | |
| O00155 | Probable G-protein coupled receptor 25 | |
| O00270 | 12-(S)-hydroxy-5,8,10,14-eicosatetraenoic acid receptor | |
| O00337 | Sodium/nucleoside cotransporter 1 | 618477 |
| O00590 | Atypical chemokine receptor 2 | |
| O14494 | Phospholipid phosphatase 1 {ECO:0000312\|HGNC:HGNC:9228} | |
| O14495 | Phospholipid phosphatase 3 {ECO:0000312\|HGNC:HGNC:9229} | |
| O14764 | Gamma-aminobutyric acid receptor subunit delta | 613060 |
| O15118 | NPC intracellular cholesterol transporter 1 {ECO:0000312\|HGNC:HGNC:7897} | 257220 |
| O15121 | Sphingolipid delta(4)-desaturase DES1 | 618404 |
| O15126 | Secretory carrier-associated membrane protein 1 | |
| O15218 | G-protein coupled receptor 182 | |
| O15244 | Solute carrier family 22 member 2 | |
| O15245 | Solute carrier family 22 member 1 | |
| O15260 | Surfeit locus protein 4 | |
| O15399 | Glutamate receptor ionotropic, NMDA 2D | 617162 |
| O15427 | Monocarboxylate transporter 4 | |
| O15438 | Canalicular multispecific organic anion transporter 2 | |
| O15440 | Multidrug resistance-associated protein 5 | |
| O43155 | Leucine-rich repeat transmembrane protein FLRT2 | |
| O43157 | Plexin-B1 | |
| O43184 | Disintegrin and metalloproteinase domain-containing protein 12 | |
| O43193 | Motilin receptor | |
| O43306 | Adenylate cyclase type 6 | 616287 |
| O43451 | Maltase-glucoamylase, intestinal | |
| O43603 | Galanin receptor type 2 | |
| O43657 | Tetraspanin-6 | |
| O43688 | Phospholipid phosphatase 2 {ECO:0000312\|HGNC:HGNC:9230} | |
| O60266 | Adenylate cyclase type 3 | 601665 |
| O60669 | Monocarboxylate transporter 2 | |
| O60906 | Sphingomyelin phosphodiesterase 2 | |

| | | |
|---|---|---|
| O75022 | Leukocyte immunoglobulin-like receptor subfamily B member 3 | |
| O75056 | Syndecan-3 | |
| O75096 | Low-density lipoprotein receptor-related protein 4 | 212780 |
| O75311 | Glycine receptor subunit alpha-3 | |
| O75387 | Large neutral amino acids transporter small subunit 3 | |
| O94759 | Transient receptor potential cation channel subfamily M member 2 | |
| O94823 | Probable phospholipid-transporting ATPase VB | |
| O94910 | Adhesion G protein-coupled receptor L1 {ECO:0000312\|HGNC:HGNC:20973} | |
| O95069 | Potassium channel subfamily K member 2 | |
| O95180 | Voltage-dependent T-type calcium channel subunit alpha-1H | 611942 |
| O95255 | Multidrug resistance-associated protein 6 | 264800 |
| O95427 | GPI ethanolamine phosphate transferase 1 | 614080 |
| O95477 | ATP-binding cassette sub-family A member 1 | 205400 |
| O95490 | Adhesion G protein-coupled receptor L2 {ECO:0000303\|PubMed:25713288} | |
| O95857 | Tetraspanin-13 | |
| O95866 | Megakaryocyte and platelet inhibitory receptor G6b {ECO:0000312\|HGNC:HGNC:13937} | 617441 |
| O95907 | Monocarboxylate transporter 3 | |
| P00533 | Epidermal growth factor receptor | 211980 |
| P01130 | Low-density lipoprotein receptor | 143890 |
| P04626 | Receptor tyrosine-protein kinase erbB-2 | 137800 |
| P05067 | Amyloid-beta precursor protein {ECO:0000305} | 104300 |
| P06734 | Low affinity immunoglobulin epsilon Fc receptor | |
| P06756 | Integrin alpha-V | |
| P07204 | Thrombomodulin | 614486 |
| P07510 | Acetylcholine receptor subunit gamma | 253290 |
| P08172* | Muscarinic acetylcholine receptor M2 | 608516 |
| P08173 | Muscarinic acetylcholine receptor M4 | |
| P08912 | Muscarinic acetylcholine receptor M5 | |
| P09619 | Platelet-derived growth factor receptor beta | 131440 |
| P0C604 | Olfactory receptor 4A8 | |
| P0C645 | Olfactory receptor 4E1 | |
| P10912 | Growth hormone receptor | 262500 |
| P11229 | Muscarinic acetylcholine receptor M1 | |
| P11230 | Acetylcholine receptor subunit beta | 616313 |
| P11836* | B-lymphocyte antigen CD20 | 613495 |
| P14867 | Gamma-aminobutyric acid receptor subunit alpha-1 | 611136 |
| P16109 | P-selectin | 601367 |
| P16410 | Cytotoxic T-lymphocyte protein 4 | 152700 |
| P16422 | Epithelial cell adhesion molecule | 613217 |
| P17152 | Transmembrane protein 11, mitochondrial | |
| P18505 | Gamma-aminobutyric acid receptor subunit beta-1 | 617153 |
| P18507 | Gamma-aminobutyric acid receptor subunit gamma-2 | 618396 |
| P20963* | T-cell surface glycoprotein CD3 zeta chain | 610163 |

| | | |
|---|---|---|
| P21452 | Substance-K receptor | |
| P21728 | D(1A) dopamine receptor | |
| P21730 | C5a anaphylatoxin chemotactic receptor 1 | |
| P21754 | Zona pellucida sperm-binding protein 3 | 617712 |
| P21860 | Receptor tyrosine-protein kinase erbB-3 | 607598 |
| P21917 | D(4) dopamine receptor | |
| P21964 | Catechol O-methyltransferase | 181500 |
| P22223 | Cadherin-3 | 601553 |
| P23415 | Glycine receptor subunit alpha-1 | 149400 (Hyperekplexia) Arg299Leu, Arg299Gln, Leu319Pro |
| P23416 | Glycine receptor subunit alpha-2 | |
| P23634 | Plasma membrane calcium-transporting ATPase 4 {ECO:0000305} | |
| P24046 | Gamma-aminobutyric acid receptor subunit rho-1 | |
| P25024 | C-X-C chemokine receptor type 1 | |
| P25025 | C-X-C chemokine receptor type 2 | |
| P25090 | N-formyl peptide receptor 2 | |
| P25103 | Substance-P receptor | |
| P25106 | Atypical chemokine receptor 3 | |
| P28472 | Gamma-aminobutyric acid receptor subunit beta-3 | 612269 617113 (Encephalopathy) Gln249Lys, Leu256Gln |
| P28476 | Gamma-aminobutyric acid receptor subunit rho-2 | |
| P28566 | 5-hydroxytryptamine receptor 1E | |
| P28827 | Receptor-type tyrosine-protein phosphatase mu | |
| P28906 | Hematopoietic progenitor cell antigen CD34 | |
| P28908 | Tumor necrosis factor receptor superfamily member 8 {ECO:0000312\|HGNC:HGNC:11923} | |
| P29033 | Gap junction beta-2 protein | 220290 |
| P29274 | Adenosine receptor A2a | |
| P29317 | Ephrin type-A receptor 2 | 116600 |
| P30273 | High affinity immunoglobulin epsilon receptor subunit gamma | |
| P30301 | Lens fiber major intrinsic protein | 615274 |
| P30542 | Adenosine receptor A1 | |
| P30939 | 5-hydroxytryptamine receptor 1F | |
| P31644 | Gamma-aminobutyric acid receptor subunit alpha-5 | 618559 |
| P32239 | Gastrin/cholecystokinin type B receptor {ECO:0000303\|PubMed:8415658} | |
| P32246 | C-C chemokine receptor type 1 | |
| P32248 | C-C chemokine receptor type 7 | |
| P33032 | Melanocortin receptor 5 | |
| P34903 | Gamma-aminobutyric acid receptor subunit alpha-3 | |

| | | |
|---|---|---|
| P35368 | Alpha-1B adrenergic receptor | |
| P35916 | Vascular endothelial growth factor receptor 3 | 153100 |
| P35968 | Vascular endothelial growth factor receptor 2 | 602089 |
| P36021 | Monocarboxylate transporter 8 | 300523 |
| P36382 | Gap junction alpha-5 protein | 108770 |
| P41181 | Aquaporin-2 | 125800 |
| P41231 | P2Y purinoceptor 2 | |
| P41597 | C-C chemokine receptor type 2 | |
| P43004 | Excitatory amino acid transporter 2 | 617105 |
| P43007 | Neutral amino acid transporter A | 616657 |
| P43629 | Killer cell immunoglobulin-like receptor 3DL1 | |
| P45844 | ATP-binding cassette sub-family G member 1 | |
| P46092 | C-C chemokine receptor type 10 | |
| P47869 | Gamma-aminobutyric acid receptor subunit alpha-2 | 618557 |
| P47870 | Gamma-aminobutyric acid receptor subunit beta-2 | 617829 (Encephalopathy) Leu277Ser |
| P47871 | Glucagon receptor | |
| P48165 | Gap junction alpha-8 protein | 116200 |
| P48169 | Gamma-aminobutyric acid receptor subunit alpha-4 | |
| P48549 | G protein-activated inward rectifier potassium channel 1 | |
| P48751 | Anion exchange protein 3 | |
| P50406 | 5-hydroxytryptamine receptor 6 | |
| P51677 | C-C chemokine receptor type 3 | |
| P51686 | C-C chemokine receptor type 9 | |
| P51693 | Amyloid-like protein 1 | |
| P51790 | H(+)/Cl(-) exchange transporter 3 | |
| P51795 | H(+)/Cl(-) exchange transporter 5 | 300554 |
| P51801 | Chloride channel protein ClC-Kb | 607364 |
| P51828 | Adenylate cyclase type 7 {ECO:0000305} | |
| P52569 | Cationic amino acid transporter 2 | |
| P55017 | Solute carrier family 12 member 3 | 263800 |
| P55061 | Bax inhibitor 1 | |
| P55064 | Aquaporin-5 | 600231 |
| P55087 | Aquaporin-4 | |
| P56747 | Claudin-6 | |
| P57789 | Potassium channel subfamily K member 10 | |
| P59541 | Taste receptor type 2 member 30 | |
| P59542 | Taste receptor type 2 member 19 | |
| P59551 | Taste receptor type 2 member 60 | |
| P61073* | C-X-C chemokine receptor type 4 | 193670 |
| P78324 | Tyrosine-protein phosphatase non-receptor type substrate 1 | |
| P78334 | Gamma-aminobutyric acid receptor subunit epsilon | |
| P78348 | Acid-sensing ion channel 1 {ECO:0000303\|PubMed:10798398} | |

| | | |
|---|---|---|
| P78382* | CMP-sialic acid transporter | 603585 |
| P82251 | b(0,+)-type amino acid transporter 1 | 220100 |
| Q01814 | Plasma membrane calcium-transporting ATPase 2 | |
| Q02094 | Ammonium transporter Rh type A | 268150 |
| Q02223 | Tumor necrosis factor receptor superfamily member 17 | |
| Q03395 | Rod outer segment membrane protein 1 | 608133 |
| Q04844 | Acetylcholine receptor subunit epsilon | 605809 |
| Q05901 | Neuronal acetylcholine receptor subunit beta-3 | |
| Q06432 | Voltage-dependent calcium channel gamma-1 subunit | |
| Q07011 | Tumor necrosis factor receptor superfamily member 9 | |
| Q08357 | Sodium-dependent phosphate transporter 2 | 213600 |
| Q08462 | Adenylate cyclase type 2 | |
| Q08828 | Adenylate cyclase type 1 | 610154 |
| Q12879 | Glutamate receptor ionotropic, NMDA 2A | 245570 |
| Q13224 | Glutamate receptor ionotropic, NMDA 2B | 613970 |
| Q13258 | Prostaglandin D2 receptor | 607277 |
| Q13291 | Signaling lymphocytic activation molecule | |
| Q13507 | Short transient receptor potential channel 3 | 616410 |
| Q13563 | Polycystin-2 {ECO:0000305} | 613095 |
| Q13571 | Lysosomal-associated transmembrane protein 5 | |
| Q13733 | Sodium/potassium-transporting ATPase subunit alpha-4 | |
| Q13936 | Voltage-dependent L-type calcium channel subunit alpha-1C | 601005 |
| Q14542 | Equilibrative nucleoside transporter 2 | |
| Q14573 | Inositol 1,4,5-trisphosphate receptor type 3 | |
| Q14761 | Protein tyrosine phosphatase receptor type C-associated protein | |
| Q14773 | Intercellular adhesion molecule 4 | |
| Q14916 | Sodium-dependent phosphate transport protein 1 | |
| Q14957 | Glutamate receptor ionotropic, NMDA 2C | |
| Q15077 | P2Y purinoceptor 6 | |
| Q15303 | Receptor tyrosine-protein kinase erbB-4 | 615515 |
| Q15722 | Leukotriene B4 receptor 1 | |
| Q15758 | Neutral amino acid transporter B(0) | |
| Q16099 | Glutamate receptor ionotropic, kainate 4 | |
| Q16288 | NT-3 growth factor receptor | |
| Q16445 | Gamma-aminobutyric acid receptor subunit alpha-6 | |
| Q16478 | Glutamate receptor ionotropic, kainate 5 | |
| Q16515 | Acid-sensing ion channel 2 {ECO:0000303\|PubMed:10842183} | |
| Q16563 | Synaptophysin-like protein 1 | |
| Q16581 | C3a anaphylatoxin chemotactic receptor | |
| Q16720 | Plasma membrane calcium-transporting ATPase 3 | 302500 |
| Q16849 | Receptor-type tyrosine-protein phosphatase-like N | |
| Q1HG43 | Dual oxidase maturation factor 1 | |
| Q3KNW5 | Solute carrier family 10 member 6 | |

| | | |
|---|---|---|
| Q4VNC1 | Probable cation-transporting ATPase 13A4 | |
| Q5GH70 | XK-related protein 9 | |
| Q5JW98 | Calcium homeostasis modulator protein 4 | |
| Q5JXX5 | Glycine receptor subunit alpha-4 | |
| Q5R3K3 | Calcium homeostasis modulator protein 6 | |
| Q5T197 | E3 ubiquitin-protein ligase DCST1 {ECO:0000305} | |
| Q5T3U5 | Multidrug resistance-associated protein 7 | |
| Q68CP4 | Heparan-alpha-glucosaminide N-acetyltransferase | 252930 |
| Q6DN72 | Fc receptor-like protein 6 | |
| Q6IF82 | Olfactory receptor 4A47 | |
| Q6P5W5 | Zinc transporter ZIP4 | 201100 |
| Q6PJF5 | Inactive rhomboid protein 2 | 148500 |
| Q6UVM3 | Potassium channel subfamily T member 2 | 617771 |
| Q6ZMH5 | Zinc transporter ZIP5 | 615946 |
| Q6ZN44 | Netrin receptor UNC5A | |
| Q6ZSA7 | Leucine-rich repeat-containing protein 55 | |
| Q6ZUX7 | LHFPL tetraspan subfamily member 2 protein {ECO:0000312\|HGNC:HGNC:6588} | |
| Q7L1I2 | Synaptic vesicle glycoprotein 2B | |
| Q7LBE3 | Solute carrier family 26 member 9 | |
| Q7RTX0 | Taste receptor type 1 member 3 | |
| Q7RTX9 | Monocarboxylate transporter 14 | |
| Q7RTY0* | Monocarboxylate transporter 13 | 125853 |
| Q7Z403 | Transmembrane channel-like protein 6 | 226400 |
| Q86SJ6 | Desmoglein-4 | 607903 |
| Q86SM8 | Mas-related G-protein coupled receptor member E | |
| Q86UK0 | ATP-binding cassette sub-family A member 12 | 601277 |
| Q86UQ4 | ATP-binding cassette sub-family A member 13 | |
| Q86VB7 | Scavenger receptor cysteine-rich type 1 protein M130 | |
| Q86VE9 | Serine incorporator 5 {ECO:0000305} | |
| Q86VZ4 | Low-density lipoprotein receptor-related protein 11 | |
| Q86W47 | Calcium-activated potassium channel subunit beta-4 | |
| Q86Y34 | Adhesion G protein-coupled receptor G3 {ECO:0000303\|PubMed:25713288} | |
| Q86YD5 | Low-density lipoprotein receptor class A domain-containing protein 3 | |
| Q8IU99 | Calcium homeostasis modulator protein 1 | |
| Q8IVJ1 | Solute carrier family 41 member 1 | |
| Q8IY26 | Phospholipid phosphatase 6 {ECO:0000312\|HGNC:HGNC:23682} | |
| Q8IYS0 | Protein Aster-C {ECO:0000250\|UniProtKB:Q8CI52} | |
| Q8IZ08 | G-protein coupled receptor 135 {ECO:0000305} | |
| Q8IZU9 | Kin of IRRE-like protein 3 | |
| Q8N1C3 | Gamma-aminobutyric acid receptor subunit gamma-1 | |
| Q8N370 | Large neutral amino acids transporter small subunit 4 | |
| Q8N7P3 | Claudin-22 | |
| Q8N8Q9 | Magnesium transporter NIPA2 | |

| | | |
|---|---|---|
| Q8NCK7 | Monocarboxylate transporter 11 | 125853 |
| Q8NCM2 | Potassium voltage-gated channel subfamily H member 5 | |
| Q8NDV2 | G-protein coupled receptor 26 | |
| Q8NFM4 | Adenylate cyclase type 4 | |
| Q8NFR9 | Interleukin-17 receptor E | |
| Q8NGA1 | Olfactory receptor 1M1 | |
| Q8NGB4 | Olfactory receptor 4S1 | |
| Q8NGC8 | Olfactory receptor 11H7 | |
| Q8NGE2 | Olfactory receptor 2AP1 | |
| Q8NGH9 | Olfactory receptor 52E4 | |
| Q8NGJ6 | Olfactory receptor 51A4 | |
| Q8NGJ7 | Olfactory receptor 51A2 | |
| Q8NGK0 | Olfactory receptor 51G2 | |
| Q8NGK1 | Olfactory receptor 51G1 | |
| Q8NGK3 | Olfactory receptor 52K2 | |
| Q8NGK4 | Olfactory receptor 52K1 | |
| Q8NGK9 | Olfactory receptor 5D16 | |
| Q8NGL9 | Olfactory receptor 4C16 | |
| Q8NGN1 | Olfactory receptor 6T1 | |
| Q8NGN8 | Putative olfactory receptor 4A4 | |
| Q8NGP3 | Olfactory receptor 5M9 | |
| Q8NGP4 | Olfactory receptor 5M3 | |
| Q8NGQ3 | Olfactory receptor 1S2 | |
| Q8NGQ4 | Olfactory receptor 10Q1 | |
| Q8NGS1 | Olfactory receptor 1J4 | |
| Q8NGS2 | Olfactory receptor 1J2 | |
| Q8NGS3 | Olfactory receptor 1J1 | |
| Q8NGW6 | Olfactory receptor 6K6 | |
| Q8NH08 | Olfactory receptor 10AC1 | |
| Q8NH09 | Olfactory receptor 8S1 | |
| Q8NH40 | Olfactory receptor 6S1 | |
| Q8NH48 | Olfactory receptor 5B3 | |
| Q8NH59 | Olfactory receptor 51Q1 | |
| Q8NH64 | Olfactory receptor 51A7 | |
| Q8NH70 | Olfactory receptor 4A16 | |
| Q8NH93 | Olfactory receptor 1L3 | |
| Q8TAB3 | Protocadherin-19 | 300088 |
| Q8TCC7 | Solute carrier family 22 member 8 | |
| Q8TCT6 | Signal peptide peptidase-like 3 {ECO:0000303\|PubMed:12077416, ECO:0000312\|HGNC:HGNC:30424} | |
| Q8TCT9 | Minor histocompatibility antigen H13 | |
| Q8TD46 | Cell surface glycoprotein CD200 receptor 1 | |
| Q8TDS5 | Oxoeicosanoid receptor 1 | |
| Q8TDS7 | Mas-related G-protein coupled receptor member D | |

| | | |
|---|---|---|
| Q8TDT2 | Probable G-protein coupled receptor 152 | |
| Q8TDY8 | Immunoglobulin superfamily DCC subclass member 4 | |
| Q8WUX1 | Sodium-coupled neutral amino acid transporter 5 | |
| Q8WWG1 | Pro-neuregulin-4, membrane-bound isoform | |
| Q8WWT9 | Solute carrier family 13 member 3 | 618384 |
| Q8WZ84 | Olfactory receptor 8D1 | |
| Q92806 | G protein-activated inward rectifier potassium channel 3 | |
| Q92911 | Sodium/iodide cotransporter | 274400 |
| Q969I6 | Sodium-coupled neutral amino acid transporter 4 | |
| Q96AM1 | Mas-related G-protein coupled receptor member F | |
| Q96AP7 | Endothelial cell-selective adhesion molecule | |
| Q96B33 | Claudin-23 | |
| Q96D53 | Atypical kinase COQ8B, mitochondrial {ECO:0000305} | 615573 |
| Q96D96 | Voltage-gated hydrogen channel 1 | |
| Q96DU3 | SLAM family member 6 | |
| Q96FT7 | Acid-sensing ion channel 4 | |
| Q96G91 | P2Y purinoceptor 11 | |
| Q96GZ6 | Solute carrier family 41 member 3 | |
| Q96J84 | Kin of IRRE-like protein 1 | |
| Q96LA5 | Fc receptor-like protein 2 | |
| Q96LA9 | Mas-related G-protein coupled receptor member X4 | |
| Q96LB0 | Mas-related G-protein coupled receptor member X3 | |
| Q96LB1 | Mas-related G-protein coupled receptor member X2 | |
| Q96LB2 | Mas-related G-protein coupled receptor member X1 | |
| Q96N87 | Inactive sodium-dependent neutral amino acid transporter B(0)AT3 | |
| Q96P88 | Putative gonadotropin-releasing hormone II receptor | |
| Q96RD9 | Fc receptor-like protein 5 | |
| Q96SJ8 | Tetraspanin-18 | |
| Q96T54 | Potassium channel subfamily K member 17 | |
| Q96TA0 | Putative protocadherin beta-18 | |
| Q99075 | Proheparin-binding EGF-like growth factor | |
| Q99466 | Neurogenic locus notch homolog protein 4 | |
| Q99624 | Sodium-coupled neutral amino acid transporter 3 | |
| Q99884 | Sodium-dependent proline transporter | |
| Q99928 | Gamma-aminobutyric acid receptor subunit gamma-3 | |
| Q9BPV8 | P2Y purinoceptor 13 | |
| Q9BQS7 | Hephaestin | |
| Q9BSK0 | MARVEL domain-containing protein 1 | |
| Q9BX84 | Transient receptor potential cation channel subfamily M member 6 | 602014 |
| Q9BXR5 | Toll-like receptor 10 | |
| Q9BY07 | Electrogenic sodium bicarbonate cotransporter 4 | |
| Q9BY15 | Adhesion G protein-coupled receptor E3 {ECO:0000303\|PubMed:25713288} | |
| Q9BZJ6 | Probable G-protein coupled receptor 63 | |

| | | |
|---|---|---|
| Q9BZJ7 | G-protein coupled receptor 62 {ECO:0000305} | |
| Q9C0K1 | Zinc transporter ZIP8 | 616721 |
| Q9H158 | Protocadherin alpha-C1 | |
| Q9H195 | Mucin-3B {ECO:0000305} | |
| Q9H228 | Sphingosine 1-phosphate receptor 5 | |
| Q9H295 | Dendritic cell-specific transmembrane protein | |
| Q9H2C8 | Olfactory receptor 51V1 | |
| Q9H2S1 | Small conductance calcium-activated potassium channel protein 2 | |
| Q9H313 | Protein tweety homolog 1 | |
| Q9H427 | Potassium channel subfamily K member 15 | |
| Q9HAR2 | Adhesion G protein-coupled receptor L3 {ECO:0000312\|HGNC:HGNC:20974} | |
| Q9HBW0 | Lysophosphatidic acid receptor 2 | |
| Q9HCC8 | Glycerophosphoinositol inositolphosphodiesterase GDPD2 | |
| Q9HCN3 | Post-GPI attachment to proteins factor 6 {ECO:0000303\|PubMed:27881714} | |
| Q9HCX4 | Short transient receptor potential channel 7 | |
| Q9NP94 | Zinc transporter ZIP2 | |
| Q9NPC1 | Leukotriene B4 receptor 2 | |
| Q9NPY3 | Complement component C1q receptor | |
| Q9NQ40 | Solute carrier family 52, riboflavin transporter, member 3 | 211530 |
| Q9NQS3 | Nectin-3 | |
| Q9NRJ7 | Protocadherin beta-16 | |
| Q9NRR2 | Tryptase gamma | |
| Q9NRU3 | Metal transporter CNNM1 | |
| Q9NS66 | Probable G-protein coupled receptor 173 | |
| Q9NS67 | Probable G-protein coupled receptor 27 | |
| Q9NSA0 | Solute carrier family 22 member 11 | |
| Q9NW15 | Anoctamin-10 | 613728 |
| Q9NXK6 | Membrane progestin receptor gamma {ECO:0000303\|PubMed:23763432} | |
| Q9NY15 | Stabilin-1 | |
| Q9NY26 | Zinc transporter ZIP1 | |
| Q9NYQ7 | Cadherin EGF LAG seven-pass G-type receptor 3 | |
| Q9NYW0 | Taste receptor type 2 member 10 | |
| Q9NZD1 | G-protein coupled receptor family C group 5 member D | |
| Q9NZQ8 | Transient receptor potential cation channel subfamily M member 5 | |
| Q9P0L9 | Polycystic kidney disease 2-like 1 protein | |
| Q9P1Z3 | Potassium/sodium hyperpolarization-activated cyclic nucleotide-gated channel 3 | |
| Q9UBK5 | Hematopoietic cell signal transducer | |
| Q9UBN4 | Short transient receptor potential channel 4 | |
| Q9UBS5 | Gamma-aminobutyric acid type B receptor subunit 1 | |
| Q9UEF7 | Klotho | 617994 |
| Q9UGI6 | Small conductance calcium-activated potassium channel protein 3 | 618658 |
| Q9UGN4 | CMRF35-like molecule 8 | |
| Q9UGQ3 | Solute carrier family 2, facilitated glucose transporter member 6 | |

| | | |
|---|---|---|
| Q9UHC3 | Acid-sensing ion channel 3 | |
| Q9UHI7 | Solute carrier family 23 member 1 | |
| Q9UHW9 | Solute carrier family 12 member 6 | 218000 |
| Q9UHX3 | Adhesion G protein-coupled receptor E2 {ECO:0000303\|PubMed:25713288} | 125630 |
| Q9UI40 | Sodium/potassium/calcium exchanger 2 | |
| Q9UJ99 | Cadherin-22 | |
| Q9UKL2 | Olfactory receptor 52A1 | |
| Q9UL51 | Potassium/sodium hyperpolarization-activated cyclic nucleotide-gated channel 2 | |
| Q9UL62 | Short transient receptor potential channel 5 | |
| Q9UM47 | Neurogenic locus notch homolog protein 3 | 125310 |
| Q9UM73 | ALK tyrosine kinase receptor | 613014 |
| Q9UMD9 | Collagen alpha-1(XVII) chain | 226650 |
| Q9UN66 | Protocadherin beta-8 {ECO:0000305} | |
| Q9UN67 | Protocadherin beta-10 | |
| Q9UN71 | Protocadherin gamma-B4 | |
| Q9UN76 | Sodium- and chloride-dependent neutral and basic amino acid transporter B(0+) | |
| Q9UN88 | Gamma-aminobutyric acid receptor subunit theta | |
| Q9UNQ0 | ATP-binding cassette sub-family G member 2 | |
| Q9UP95 | Solute carrier family 12 member 4 | |
| Q9UPZ6 | Thrombospondin type-1 domain-containing protein 7A | |
| Q9Y210 | Short transient receptor potential channel 6 | 603965 |
| Q9Y289 | Sodium-dependent multivitamin transporter | |
| Q9Y2T6 | G-protein coupled receptor 55 | |
| Q9Y5E1 | Protocadherin beta-9 | |
| Q9Y5E2 | Protocadherin beta-7 | |
| Q9Y5E3 | Protocadherin beta-6 {ECO:0000305} | |
| Q9Y5E4 | Protocadherin beta-5 | |
| Q9Y5E5 | Protocadherin beta-4 | |
| Q9Y5E6 | Protocadherin beta-3 | |
| Q9Y5E7 | Protocadherin beta-2 | |
| Q9Y5E8 | Protocadherin beta-15 | |
| Q9Y5E9 | Protocadherin beta-14 | |
| Q9Y5F0 | Protocadherin beta-13 | |
| Q9Y5F1 | Protocadherin beta-12 | |
| Q9Y5F2 | Protocadherin beta-11 | |
| Q9Y5F8 | Protocadherin gamma-B7 | |
| Q9Y5F9 | Protocadherin gamma-B6 | |
| Q9Y5G0 | Protocadherin gamma-B5 | |
| Q9Y5G1 | Protocadherin gamma-B3 | |
| Q9Y5G2 | Protocadherin gamma-B2 | |
| Q9Y5G3 | Protocadherin gamma-B1 | |
| Q9Y5I7 | Claudin-16 | 248250 |
| Q9Y5Y4 | Prostaglandin D2 receptor 2 | |

| | | |
|---|---|---|
| Q9Y5Z0 | Beta-secretase 2 | |
| Q9Y6C5 | Protein patched homolog 2 | 155255 |
| Q9Y6W8* | Inducible T-cell costimulator | 607594 |

* Absent from the humsavar dataset of human variants

The 3d column shows:

- The phenotype number of a disease in the Mendelian Inheritance in Man (MIM) compendium of human genes and genetic phenotypes
- When applicable, the variant associated to the disease
